# Transgressive Hybrids as Hopeful Holobionts

**DOI:** 10.1101/2024.08.17.607925

**Authors:** Benjamin Thomas Camper, Andrew Stephen Kanes, Zachary Tyler Laughlin, Riley Tate Manuel, Sharon Anne Bewick

## Abstract

**Background:** Hybridization between evolutionary lineages has profound impacts on the fitness and ecology of hybrid progeny. In extreme cases, the effects of hybridization can transcend ecological timescales by introducing trait novelty upon which evolution can act. Indeed, hybridization can even have macroevolutionary consequences, for example, as a driver of adaptive radiations and evolutionary innovations. Accordingly, hybridization is now recognized as a motor for macrobial evolution. By contrast, there has been substantially less progress made towards understanding the positive eco-evolutionary consequences of hybridization on holobionts. Rather, the emerging paradigm in holobiont literature is that hybridization disrupts symbiosis between a host lineage and its microbiota, leaving hybrids at a fitness deficit. These conclusions, however, have been drawn based on results from predominantly low-fitness hybrid organisms. Studying ‘dead-end’ hybrids all but guarantees finding that hybridization is detrimental. This is the pitfall that Dobzhansky fell into over 80 years ago when he used hybrid sterility and inviability to conclude that hybridization hinders evolution. Goldschmidt, however, argued that rare saltational successes—so-called ‘hopeful monsters’—disproportionately drive positive evolutionary outcomes. Goldschmidt’s view is now becoming a widely accepted explanation for the prevalence of historical hybridization in extant macrobial lineages. Aligning holobiont research with this broader evolutionary perspective requires recognizing the importance of similar patterns in host–microbiome systems. That is, rare and successful ‘hopeful holobionts’ (i.e., hopeful monsters at the holobiont scale) might be disproportionately responsible for holobiont evolution. If true, then it is these successful systems that we should be studying to assess impacts of hybridization on the macroevolutionary trajectories of host– microbiome symbioses.

**Results:** In this paper, we explore the effects of hybridization on the gut (cloacal) and skin microbiota in an ecologically successful hybrid lizard, *Aspidoscelis neomexicanus*. Specifically, we test the hypothesis that hybrid lizards have host-associated (HA) microbiota traits strongly differentiated from their progenitor species. Across numerous hybrid microbiota phenotypes, we find widespread evidence of transgressive segregation. Further, microbiota restructuring broadly correlates with niche restructuring during hybridization. This suggests a relationship between HA microbiota traits and ecological success.

**Conclusion:** Transgressive segregation of HA microbiota traits is not limited to hybrids at a fitness deficit but also occurs in ecologically successful hybrids. This suggests that hybridization may be a mechanism for generating novel and potentially beneficial holobiont phenotypes. Supporting such a conclusion, the correlations that we find between hybrid microbiota and the hybrid niche indicate that hybridization might change host microbiota in ways that promote a shift or an expansion in host niche space. If true, hybrid microbiota restructuring may underly ecological release from progenitors. This, in turn, could drive evolutionary diversification. Using our system as an example, we elaborate on the evolutionary implications of host hybridization within the context of holobiont theory and then outline the next steps for understanding the role of hybridization in holobiont research.

## Introduction

Hybridization in macrobial systems facilitates evolution through lineage unification, typically via unidirectional introgression or bidirectional admixture. This contrasts with more traditional mechanisms of evolution involving lineage diversification. Whereas genomic change due to divergent processes tends to be incremental, hybridization yields immediate, extensive change across an organism’s entire genome. For this reason, hybrid systems are often cradles of phenotypic novelty^1,2^ and evolutionary innovation.^3–6^ In the past, hybrid systems have been studied as generators of phenotypic novelty in behavior,^7,8^ fecundity,^9–13^ morphology,^14–17^ and physiology.^18–22^ Recently, however, there has been growing recognition that many ecologically relevant organismal traits are not determined by an organism’s genes alone. Instead, macroorganisms function as ‘holobionts’ – collectives comprising both a host organism and all of its host-associated (HA) microbes.^23–25^ As a consequence, many host traits are jointly or even solely governed by the actions of HA microbes.^26–28^ How host hybridization impacts the HA microbiome, the extent to which these changes influence other host traits, and whether these changes serve as motors or brakes for holobiont evolution remain open questions.

The importance of HA microbiomes to host ecology and evolution has been recognized since at least the early 1900’s.^29,30^ However, recent advances in low-cost, next-generation sequencing have dramatically improved our understanding of how extensively and intimately HA microbiomes interact with their hosts to influence host function. Gut microbiomes, for instance, affect everything from host obesity to host metabolic phenotype and energy balance.^31–33^ As an example, colonization of germ-free mice with ‘obese’ versus ‘lean’ microbiota results in differential weight gain even when food consumption is identical.^34^ Additionally, gut microbiota can provision hosts with key nutrients or detoxify plant defensive compounds, impacting dietary niche.^35,36^ Some phloem-feeding aphids, for instance, rely on *Buchnera aphidicola* to supply tryptophan that is not available in plant sap. Similarly, *Pantoea* spp. bacteria in the guts of white-spectacled bulbuls detoxify *Ochradenus baccatus* seeds, making them palatable to the birds.^37^ Gut, skin, and vaginal microbiota can also provide pathogen resistance to their host. The amphibian skin microbiota, for instance, is an important defense mechanism against the widespread fungal pathogens *Batrachochytrium dendrobatidis* and *B. salamandrivorans*.^38–42^ Likewise, some human vaginal microbiota compositions are associated with trichomonal, gonococcal, and chlamydial infection risk,^43^ while certain gut microbial taxa in roundworms protect their hosts from pathogenic *Staphylococcus aureus* infections.^44^

The importance of host–microbiome interactions to host health and ecology has led to intense study of how HA microbiomes interact with host evolution.^45^ One shortcoming of this research is that it has primarily explored host–microbiome relationships from the perspective of host lineage diversification. Indeed, within holobiont literature, host hybridization has been viewed as a disruptive force, leading to host–microbiome mismatches and breakdowns of mutualistic relationships.^45–48^ The resulting narrow focus of holobiont research on lineage diversification is noticeably at odds with broader evolutionary perspectives that increasingly accept hybridization as an important mode of adaptive change.^49^ Indeed, in macrobial systems, hybridization is now considered something of a double-edged sword. In many cases, hybridization serves as a brake, slowing the rate of evolutionary change. But in other cases, it is actually a motor, promoting host lineage diversification.^50–52^ In the systems where hybridization is beneficial, evolutionary success is often attributed to transgressive segregation (i.e., extreme or novel phenotypes). Among tiger salamanders, for example, thermal stress often limits fitness. However, some first-generation (F_1_) hybrids between barred tiger salamanders (*Ambystoma mavortium*) and California tiger salamanders (*A. californiense*) exhibit transgressive segregation in their critical thermal maxima.^21^ This allows them to tolerate temperatures higher than either progenitor, and thus persist in locations where their progenitors cannot. Similarly, within the well-studied *Helianthus* sunflower system, hybrid flowers often gain the ability to colonize extreme environments, including desert sand dunes and salt marshes. These are habitats where neither progenitor can survive, and likely emerge as hybrid environments due to selection on transgressive phenotypes.^53,54^ Though transgressive segregation receives most of the attention, hybrid phenotypes intermediate to progenitor phenotypes may also have positive evolutionary implications. Intermediate phenotypes could, for example, be beneficial in ecotones between progenitor habitats. Given the myriad of systems where hybridization acts as a source of phenotypic novelty on host traits and, in particular, host traits that contribute to ecological success, it would be surprising if hybridization could not generate the same beneficial phenotypic novelty in holobionts. Despite this, very little effort has been invested in testing the hypothesis that hybridization can generate beneficial novelty in holobiont traits.

The potential for host hybridization to positively impact holobiont evolution has likely been ignored because the majority of holobiont hybridization studies have considered hybrids at a fitness deficit.^47^ One of the earliest examinations of microbiota in hybrid hosts, for instance, was a study of *Nasonia* wasps. In this system, the gut microbiota of hybrid *Nasonia* exhibited phenotypic novelty, differing in both microbial abundance and diversity as compared to progenitors.^55^ However, the gut microbiota of hybrid wasps were also detrimental to the host. In fact, hybrid wasps with intact gut microbiota experienced up to 90% lethality that was not observed in germ-free animals. This led to the conclusion that hybridization disrupts host–microbiome interactions. The first study of hybrid microbiota in vertebrates, an analysis of a central European *Mus musculus* hybrid zone, revealed similar outcomes. In this system, hybridization also induced large shifts in gut microbiota composition. As with *Nasonia*, however, these compositional shifts were largely disruptive, contributing to immunoregulation defects in the hybrid mice.^56^

Like the *Nasonia* and *Mus musculus* systems, most other work on hybrid HA microbiomes, at least amongst animals,^57–59^ has considered hybrid hosts that exhibit fitness deficits. Focusing on ‘dead-end’ hybrids, however, strongly biases results towards finding the negative impacts of host hybridization on host–microbiome symbioses. Thus, while holobiont fitness deficits may be the most likely outcomes of host hybridization, dead-end hybrids are not the most fruitful for understanding the positive evolutionary consequences of hybridization. Notably, associating prevalence with significance was the mistake made by Dobzhansky who, during early hybridization research, argued that hybridization hinders macrobial evolution based on the fact that hybrid organisms are frequently sterile and/or inviable.^11,60–63^ Goldschmidt, on the other hand, avoided this logical fallacy by recognizing that rare positive outcomes— so-called ‘hopeful monsters’—could disproportionately drive evolutionary diversification^64,65^ (but see discussion of criticisms preventing the early adoption of Goldschmidt’s hopeful monsters^66,67^). Although Goldschmidt’s arguments were formulated in a broader ‘macromutation’ context, the pattern that he described has since been extensively applied to hybridization. In particular, recent authors have outlined how genetic restructuring through hybridization serves as a common mechanism for generating macromutations^68–73^ that can drive evolutionary diversification.^5,51,74–79^

Shortly after the postulation of Goldschmidt’s hopeful monsters, biologists such as Anderson^80–82^ and Stebbins^83,84^ introduced hybridization into the lexicon of macrobial evolution, offering it as a mechanism for generating beneficial genetic variation in populations. Since then, appreciation for the role of hybridization has accelerated, driven in part by the prevalence of DNA sequencing techniques in the field of evolution.^85^ Thus, hybridization is now understood as a common and important mode of evolution. Indeed, many successful clades show widespread evidence of historical introgression, and in some taxa, hybridization has even been associated with increased diversification rates, such as adaptive radiations.^5,51,74–76,85,86^ The prominence of historical hybridization across disparate taxa has led to a new understanding of how macrobial evolution proceeds: not always as a divergent tree, but as a ‘reticulate network’ comprising both lineage divergence and unification.^87^ Aligning holobiont research with these broader evolutionary perspectives requires recognizing the importance of similar patterns in host– microbiome systems. Like Goldschmidt’s hopeful monsters, rare, successful ‘hopeful holobionts’ might be disproportionately responsible for holobiont evolution, even when most hybrid holobionts suffer fitness deficits (e.g., disruptions to host–microbe symbioses). If this hypothesis is true, then the impact of holobiont trait novelty on evolutionary success will likely only be apparent by focusing on successful hybrid systems. Several recent studies have examined HA microbiomes in hybrid systems without apparent fitness deficits.^88,89^ Similar to the studies of *Nasonia* wasps and hybrid mice,^55,56^ these studies have identified hybrid HA microbiomes that are distinct from their progenitor species. Thus, there is opportunity for positive evolutionary outcomes of holobiont hybridization, including novel microbial phenotypes that might enable shifts and/or expansions of the holobiont into new ecological space. Unfortunately, most studies of HA microbiomes in hosts without an obvious fitness deficit have taken place in the lab or have used captive animals. Because of the strong effects of captivity on HA microbiomes,^90–92^ interpretation of these systems and their implications for wild animal populations and *in situ* eco-evolutionary processes are less clear.

Whiptail lizards (genus=*Aspidoscelis*) of the southwestern USA and northern Central America serve as unique examples of ecologically successful hybrid vertebrates. Among *Aspidoscelis*, there are a total of 8 diploid and 7 triploid parthenogenetic species^93–95^ (although these numbers may change based on new discoveries and species definitions^96–98)^. All were spawned as a result of hybridization between different progenitor species (although see Cole et al.^99,100^ for lab-reared parthenogenetic tetraploid species), and many exhibit ecological shifts^101–103^ or expansion^104–107^ into novel habitats and locations. Beyond being hallmark examples of successful hybridization, whiptail lizards have several other advantages. Unlike many traditional hybrid systems that backcross with their progenitor populations, parthenogenetic whiptails emerge through single hybridization events that give rise to independently reproducing clonal lineages.^107,108^ Thus, populations (commonly termed ‘arrays’^97,109^ though we will refer to them as ‘populations’ to avoid confusion) of hybrid parthenogens maintain very little interindividual genetic variation and effectively represent the ‘frozen’ F_1_ generation of a hybrid cross. This means that, unlike introgression zones, hybrid parthenogens have a fixed genetic composition that is almost fully intermediate (i.e., 50% maternal progenitor genome and 50% paternal progenitor genome) to either progenitor species. Further, because of their intermediate genetic status, hybrid parthenogens exhibit maximum heterozygosity at alleles that are differentially fixed in progenitor populations.^110^ Additionally, because parthenogens resemble F_1_ generations, the ‘front-end’ effects of hybridization are effectively preserved. This permits better study of these effects than most other types of hybrid systems. A final and important advantage of whiptails is that many parthenogenetic *Aspidoscelis* species occur syntopically or, at least, sympatrically with one or both progenitor species.^105,106,111–113^ As a result, *Aspidoscelis* systems are excellent natural laboratories for comparing traits, including microbial phenotypes, between progenitor species and their hybrid offspring while controlling for the effects of varying habitat. Despite these many advantages, few published studies have addressed the effects of hybridization on HA microbiomes of parthenogenetic whiptails or even hybrid vertebrate parthenogens in general.

In this paper, we use the ecologically successful diploid parthenogen, *Aspidoscelis neomexicanus*, to test the hypothesis that successful hybrid holobionts exhibit HA microbiota traits that are differentiated from those of their progenitors. The alternative hypothesis is that hybrid holobionts are only successful when HA microbiota traits are conserved (i.e., when traits are similar to one or both parents). Finding that most *A. neomexicanus* HA microbiota traits are, in fact, distinct from their progenitors, we further test the hypothesis that successful hybrid holobionts exhibit HA microbiota traits that are transgressive to their progenitors. The alternative hypothesis is that successful hybrid holobionts are characterized by intermediate HA microbiota traits. We expect *A. neomexicanus* HA microbiota traits to be transgressive rather than intermediate for several reasons. First, *A. neomexicanus* exhibits some transgressive ecological traits.^106,114,115^ Thus, to the extent that HA microbiota traits either cause or reflect an altered ecological niche, the HA microbiota traits of *A. neomexicanus* should be transgressive. Second, existing data from macrobial systems suggests that transgressive phenotypes are more common than intermediate phenotypes among ecologically and/or evolutionarily successful hybrids.^116^ To test our hypotheses, we sampled both the skin (cutaneous) and cloacal (representative of ‘gut’) microbiota of *Aspidoscelis neomexicanus*, where it occurs syntopically with each of its progenitor species, *A. marmoratus* (maternal) and *A. inornatus* (paternal).^93^ We then considered a range of different HA microbiota traits, including both diversity and composition measures. For each trait, we classified *A. neomexicanus* as transgressive (novel), intermediate (between each progenitor), or conserved (resembling one or both progenitors) relative to its two progenitor species. Our findings suggest that most HA microbiota traits in *A. neomexicanus* are indeed distinct from and transgressive to the traits of both progenitor species. By relating observed differences between hybrid and progenitor microbiota to known differences in ecological niche, we use these results to speculate on potential mechanisms by which hybridization produces phenotypic novelty in holobionts and the possible ecological consequences of such novelty. More broadly, we use our results to discuss the largely overlooked potential for ecological differences that emerge in the holobiont to influence holobiont macroevolution. Consistent with early evolutionary theory, the field of holobiont evolution has adopted a paradigm wherein hybridization is viewed as a primarily negative phenomenon. By shifting focus to successful hybrid systems, we aim to align holobiont research with modern evolutionary theory that recognizes hybridization as an important mode of evolution.

## Results

Throughout the ‘Results’ section, we apply phylogenetically aware incidence-based metrics (i.e., Faith’s Phylogenetic Diversity^117^ and unweighted UniFrac^118^) to amplicon sequence variants (ASVs, i.e. - sequences that differ by at least one base pair, regardless of taxonomic assignment) to characterize microbiota structure and composition. We choose an incidence-based approach because, unlike an abundance-based approach, it can be applied to all of the HA microbiota traits that we consider, including traits that are calculated across multiple lizards (e.g., *β*-diversity, *γ*-diversity). We choose a phylogenetically aware approach because it captures differentiation in a less arbitrary manner than using a specific taxonomic level (e.g., species, genera) or similarity threshold (e.g., 97% similar operational taxonomic units). However, we also provide numerous abundance-based and phylogenetically agnostic measures, including genus-level analyses, in the Additional files. As expected, most of our results are consistent across metrics, though there are occasional differences. Throughout the remainder of the manuscript, we refer to ‘population’ as a single lizard species from a single study location (e.g., *A. neomexicanus* at SBluG).

### Microbiota Diversity

To test our hypotheses that HA microbiota diversity traits are 1) distinct and 2) transgressive in hybrid animals, we compare hybrid and progenitor HA microbiota diversity as follows:

- If hybrid HA microbiota diversity is significantly lower or significantly higher than both progenitors, hybrid HA microbiota diversity is transgressive.
- If hybrid HA microbiota diversity is significantly lower than one progenitor and significantly higher than the other progenitor, hybrid HA microbiota diversity is intermediate.
- If hybrid HA microbiota diversity is not significantly different from one or both progenitors, hybrid HA microbiota diversity is conserved.

#### *⍺*-diversity

The median *⍺*-diversity of the gut (see Fig. 1A) and skin (see Fig. 1E) microbiota of hybrid lizards (*A. neomexicanus*) is higher than the median *⍺*-diversity of the gut and skin microbiota of progenitors (*A. marmoratus* at SNW and *A. inornatus* at SBluG). This is true not only when comparing syntopic lizards (*A. neomexicanus* with *A. inornatus* at SBluG or *A. neomexicanus* with *A. marmoratus* at SNW), but also when making comparisons across locations (*A. neomexicanus* at SNW with *A. inornatus* at SBluG or *A. neomexicanus* at SBluG with *A. marmoratus* at SNW). However, differences in *⍺*-diversity are not significant with either progenitor for gut microbiota and are only significant for skin microbiota at one location, SBluG.(see Additional file 1: Figs. 1.1-1.6 for additional metrics). That said, combining the two *A. neomexicanus* populations into a single ‘asexual’ class and the two progenitor populations (*A. inornatus* + *A. marmoratus*) into a single ‘bisexual’ class results in significantly greater hybrid *⍺*-diversity (see Additional file 1: Figs. 1.2 and 1.4). Thus, *⍺*-diversity of hybrid microbiota appears to be conserved with both progenitors but may be weakly transgressive. If *⍺*-diversity is transgressive for either gut or skin microbiota, then the effect size is small and difficult to detect without a larger sample size (i.e., by pooling the progenitor populations and the two hybrid populations).

**Figure 1.**
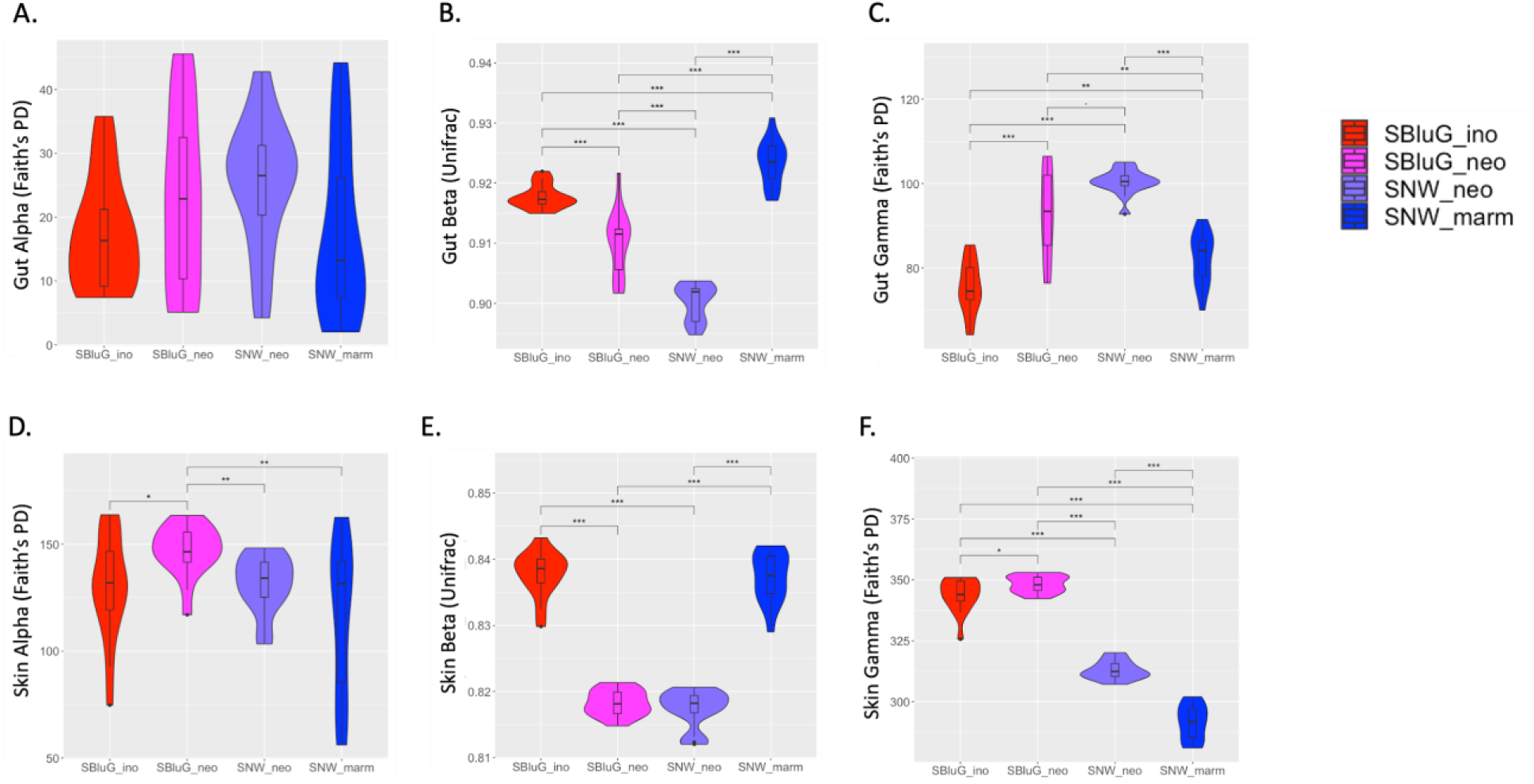
Comparison of diversity between populations of *Aspidoscelis inornatus* from SBluG (red), *A. neomexicanus* from SBluG (magenta), *A. neomexicanus* from SNW (purple), and *A. marmoratus* from SNW (blue) for both gut (A–C) and skin (D–F) microbiota (ASVs) and considering *⍺*-diversity (A, D), *β*-diversity (B, E), and *γ*-diversity (C, F). Panels (A, D) are based on the microbiota of individual lizards from each population; panels (B, C, E, F) are based on 15 subsamples of 12 individuals from each population. Significant differences in diversity between groups, as determined by a Kruskal-Wallis test followed by post hoc Pairwise Wilcox tests using a Benjamini-Hochberg correction, are indicated as follows: p-value ≤ 0.001 (***), p-value ≤ 0.01 (**), p-value ≤ 0.05 (*), p-value ≤ 0.1 (.) Additional analyses using alternate diversity metrics can be found in Additional file 1.

#### *β*-diversity

*β*-diversity of the gut and skin microbiota of the two hybrid lizard (*A. neomexicanus*) populations is significantly lower than *β*-diversity of the gut and skin microbiota of the two progenitor populations (*A. marmoratus* at SNW and *A. inornatus* at SBluG). Again, this trend holds both within and across locations. Thus, *A. inornatus* microbiota at SBluG have higher *β*-diversity than *A. neomexicanus* microbiota at either SBluG or SNW, while *A. marmoratus* microbiota at SNW have higher *β*-diversity than *A. neomexicanus* microbiota at either SBluG or SNW (see Additional file 1: Figs. 1.7–1.8 for additional metrics). Thus, *β*-diversity of hybrid gut and skin microbiota is transgressive.

#### *γ*-diversity

*γ*-diversity of the gut microbiota of both hybrid lizard (*A. neomexicanus*) populations is higher than *γ*-diversity of the gut microbiota of either progenitor population. For skin microbiota, however, there are large differences in *γ*-diversity across locations, with both lizard populations at SBluG exhibiting significantly higher *γ*-diversity than both lizard populations at SNW. As a result, while skin microbiota *γ*-diversity of each hybrid lizard (*A. neomexicanus*) population is higher than skin microbiota *γ*-diversity of the syntopic progenitor population, skin microbiota *γ*-diversity of *A. neomexicanus* at SNW is lower than skin microbiota *γ*-diversity of *A. inornatus* at SBluG (see Additional file 1: Fig. 1.9 for additional metrics). Thus, *γ*-diversity of hybrid gut microbiota is transgressive. While *γ*-diversity of skin microbiota appears to be transgressive as well, this result is complicated by location-specific effects (see ‘Discussion’ section for details).

#### Core diversity

Using a core threshold of 50% (i.e., to be part of the core, a microbial taxon must be present on at least half of the lizards in a population), the gut and skin core microbiota of hybrid lizard (*A. neomexicanus*) populations at both SBluG and SNW are more diverse than the gut and skin core microbiota of progenitor populations (see Fig. 2A, E). Again, this is true for comparisons both within and between locations and is also true for most other core thresholds (see Additional file 2: Figs. 2.3 and 2.6, see also Additional file 2: Figs. 2.1-2.7 for additional metrics). For gut microbiota, the high diversity of the hybrid core relative to either progenitor appears to be driven not only by the overall higher *γ*-diversity of hybrid microbiota but also by the larger percentage of hybrid *γ*-diversity that is part of the core (see Fig. 2B). For skin, this is not the case, as both hybrid populations and the *A. marmoratus* population have similar percentages of *γ*-diversity within their core (see Additional file 2: Figs. 2.8-2.14 for additional metrics). Thus, the absolute core diversities of hybrid gut and skin microbiota are <u>transgressive</u>. The percentage of *γ*-diversity that is part of the hybrid core is also <u>transgressive</u> for gut microbiota but appears to be <u>conserved</u> for skin microbiota.

**Figure 2.**
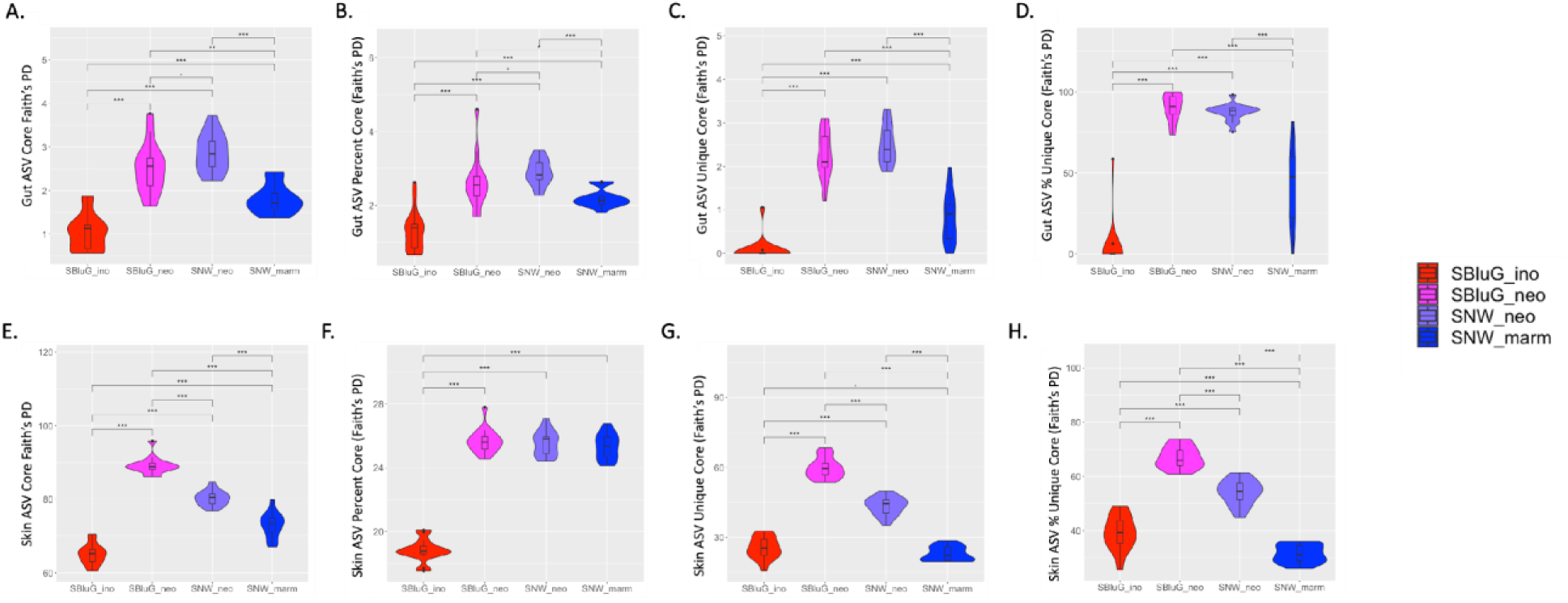
Diversity of the core gut (A–D) and skin (E–H) microbiota (ASVs) for *Aspidoscelis inornatus* from SBluG (red), *A. neomexicanus* from SBluG (magenta), *A. neomexicanus* from SNW (purple), and *A. marmoratus* from SNW (blue), assuming a 50% core threshold (i.e., to be considered part of the core, a microbial taxon must be present on at least half of the animals in a population). Panels (A, E) show the diversity of the core microbiota for each population. Panels (B, F) show the percentage of the overall *γ*-diversity of each population that is part of the respective core microbiota. Panels (C, G) show the diversity of the component of the core microbiota that is unique to each population. Panels (D, H) show the percentage of the core microbiota that is unique to each population. Additional information on the core microbiota, including analyses at alternative core thresholds and Venn Diagrams, can be found in Additional file 2.

#### Unique core diversity

At a core threshold of 50%, many of the microbial taxa and much of the microbial phylogenetic diversity comprising the core gut and skin microbiota of the four lizard populations are unique to their respective species (i.e., not shared with other lizard species, see Fig. 2 C, G as well as Additional file 2: Figs. 2.15–2.21 for additional metrics; see also the ‘Materials and Methods’ section for how we define ‘unique’). Like overall core diversity, the diversity of the unique component of core microbiota is higher amongst hybrids than amongst progenitors. This is true of both skin and gut microbiota and holds in both an absolute sense (see Figure 2C,G) and when measured as the percentage of a population’s core diversity (see Fig. 2D,H, see also Additional file 2: Figs. 2.22-2.28 for additional metrics as well as Additional file 2: Fig 2.29 for a Venn Diagram summary of shared microbial taxa). Thus, both absolute and relative diversity of the unique components of the hybrid gut and skin core microbiota are transgressive.

### Microbiota Composition

To test our hypotheses that HA microbiota composition traits are 1) distinct and 2) transgressive in hybrid animals, we use two ordination space techniques. First, we consider applied in Merot et al.^16^ to consider the position of the hybrid centroid in ordination ordination based on maximizing system-wide variation (Principal Coordinate Analysis, PCoA). Second, we use the method space relative to and perpendicular to the main axis of variation between progenitors (‘Triangle plots’).^16^ For each ordination method, we separately consider the first two ordination axes (PCoA) or the two ordination axes that emerge from the analysis (Triangle plots) and compare the positions of hybrid and progenitor HA microbiota as follows:

- If the positions of hybrid HA microbiota lie significantly above or below the positions of both progenitors, hybrid HA microbiota composition is transgressive.
- If the positions of hybrid HA microbiota lie significantly above one progenitor and significantly below the other the progenitor, hybrid HA microbiota composition is intermediate.
- If the positions of hybrid HA microbiota are not significantly different from one or both progenitors, hybrid HA microbiota composition is conserved.

#### PCoA

Hybrid gut microbiota are conserved along both the first and second UniFrac PCoA axes (see Fig. 3A and Additional file 3: Tables 3.3 and 3.4). Indeed, there are no significant differences along either PCoA axis when comparing hybrids to their syntopic progenitor, and there are only two significant differences when comparing hybrids to their non-syntopic progenitor (*A. neomexicanus* at SBluG is significantly different from *A. marmoratus* at SNW along the first PCoA axis, and *A. neomexicanus* at SNW is significantly different from *A. inornatus* at SBluG along the second PCoA axis). Results for skin microbiota are more complicated (see Fig. 3C and Additional file 3: Tables 3.11 and 3.12). *A. neomexicanus* at SBluG lies significantly higher along the first PCoA axis than either progenitor. However, *A. neomexicanus* at SNW, though also lying higher than both progenitors along the first Unifrac PCoA axis, is not significantly different from either progenitor. Along the second UniFrac PCoA axis, neither hybrid population is significantly different from its syntopic progenitor, though both hybrid populations show significant differences with the non-syntopic progenitor. The latter is perhaps not surprising, given the role of *Fodinibacter luteus* as a driver of skin microbiota variation, and the fact that *Fodinibacter luteus* appears to be location-specific, rather than species-specific (see Additional file 3: Figures 3.1-3.4 and Tables 3.1-3.16 for additional metrics). Thus, we conclude that gut microbiota composition of the hybrid is <u>conserved</u> along the first two UniFrac PCoA axes, and skin microbiota composition of the hybrid is <u>conserved</u> along the second UniFrac PCoA axis. While skin microbiota composition of the hybrid along the first UniFrac PCoA axis appears to be <u>transgressive</u>, like *γ*-diversity, interpretation of this result is complicated by location-specific effects (see ‘Discussion’ section for details; see also Additional file 4 for PERMANOVA results that are broadly consistent with these analyses).

**Figure 3.**
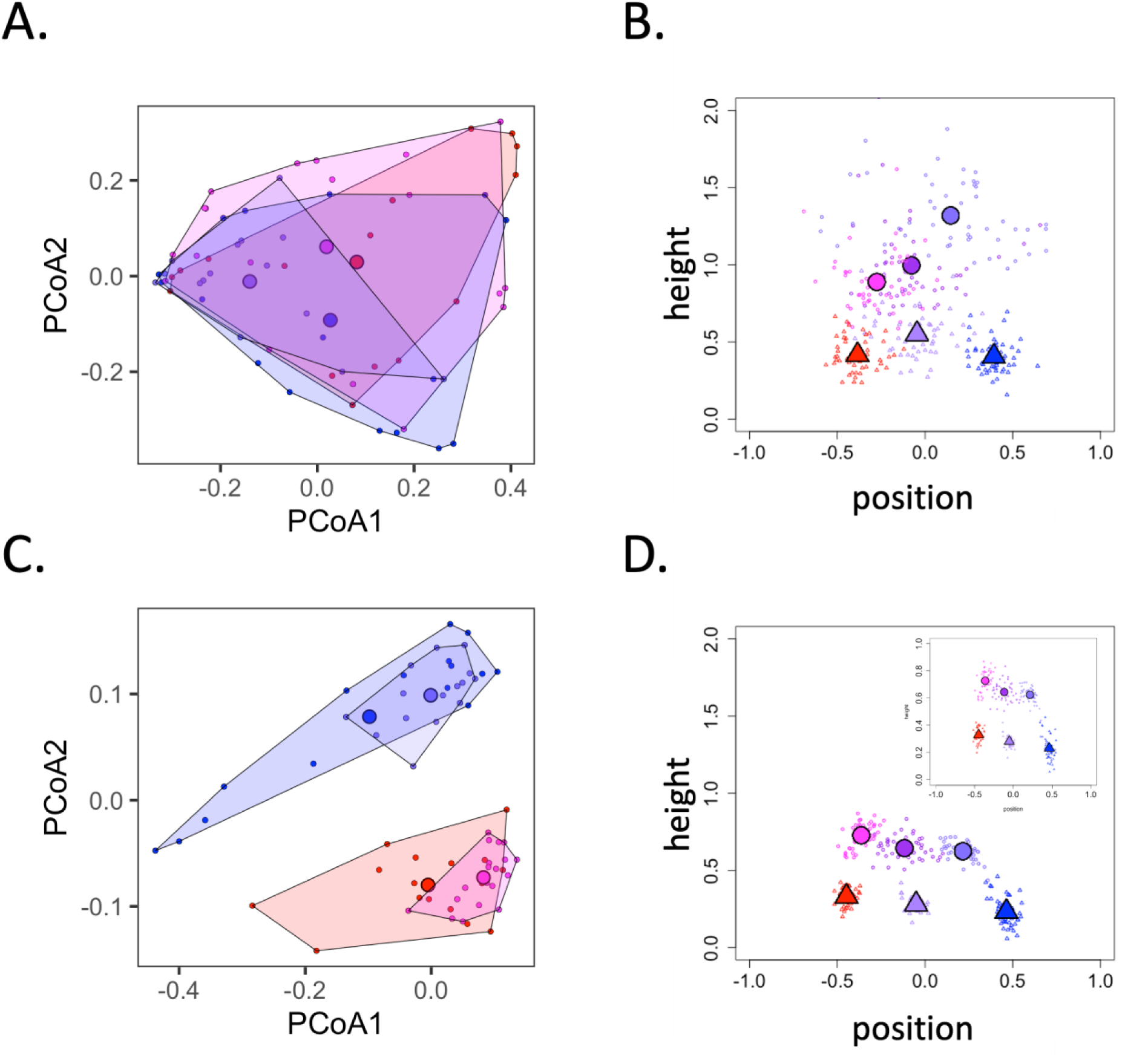
Two-dimensional PCoA (unweighted UniFrac distances) plots for gut (A) and skin (C) ASVs of *Aspidoscelis neomexicanus* from SBluG (magenta), *A. neomexicanus* from SNW (blue-purple), *A. marmoratus* (blue), and *A. inornatus* (red). Triangle plots of the hybrid mean microbiota for gut (B) and skin (D) ASVs based on unweighted UniFrac distances. The position of the hybrid microbiota projection (x-axis) represents a measure of distance towards each progenitor along the main axis of parental variation, while the height of the triangle (y-axis) represents the deviation of the mean hybrid microbiota relative to main axis of parental variation. All values are calculated in the ‘microbiota space’ defined by the first 20 principal coordinates (>95% of variance) and normalized by the distance between progenitors. Large circles are the mean value of 50 subsamples on observed data and small circles represent the outcome of each individual subsample on observed data. Large triangles are the mean value of 50 subsamples on null models and small triangles represent the outcome of each individual subsample on a null model. For observed data, we considered the *A. neomexicanus* populations from both locations (purple), only the *A. neomexicanus* location from SNW (blue-purple), and only the *A. neomexicanus* location from SBluG (magenta). For null models, we consider subsamples of the *A. inornatus* population (red), subsamples of the *A. marmoratus* population (blue), and samples based on combining *A. inornatus* + *A. marmoratus* individuals (periwinkle). For both observed data and null models, subsamples consisted of 12 randomly sampled individuals from each progenitor population (see ‘Materials and Methods’ section). Additional triangle plots can be found in Additional file 5; additional ordination plots can be found in Additional file 3.

#### Triangle Plots

Based on a 20-dimensional UniFrac projection, hybrid gut microbiota at SBluG are intermediate to both parent null models (see ‘Materials and Methods’ section for a description of the progenitor null models) along the main axis of parental variation (see Fig. 3B). By contrast, hybrid gut microbiota at SNW are not significantly different from *A. marmoratus* null models at SNW. Like the hybrid population at SBluG, however, the centroid for the hybrid population at SNW does lie intermediate to the centroids of the two progenitor null models. Along the perpendicular axis - ‘height of the triangle’ – the gut microbiota of both hybrid populations are significantly different from progenitor null models. Notice that, because of the way the height of the triangle is defined (i.e., distance from a line drawn between the parental centroids), the height of the triangle will either be conserved (not different from progenitors) or transgressive (different from progenitors). It can never be intermediate. Similar to gut microbiota, hybrid skin microbiota are intermediate to both parents along the main axis of parental variation (see Fig. 3D). However, for skin, both hybrid populations are significantly different from progenitors, making the skin microbiota of hybrid animals clearly intermediate for this trait. Also, like gut microbiota, hybrid skin microbiota are significantly different along the perpendicular height of the triangle. Thus, hybrid gut and skin microbiota are intermediate (though there is some evidence for conservation in gut microbiota at SNW) along the main axis of parental variation, and transgressive along the perpendicular height of the triangle (see Additional file 5 for additional metrics, 2D projections, and the microbial taxa that comprise the two axes of the triangle plots in Euclidean space).

### Microbial abundances

To test our hypotheses that microbial abundance traits are 1) distinct and 2) transgressive in hybrid animals, we first use Principal Component Analysis (PCA) to identify the microbial taxa responsible for variation in our system (see Additional File 3: Figs 3.1 and 3.3 and Tables 3.1, 3.2, 3.9 and 3.10). Specifically, we select the microbial taxa with the highest loadings on the first two Principal Component axes for gut and skin microbiota. We then compare the relative abundances of microbial taxa as follows:

- If the relative abundance of the microbial taxon in hybrid HA microbiota is significantly lower or significantly higher than it is in both progenitor microbiota, taxon abundance on the hybrid is transgressive.
- If the relative abundance of the microbial taxon in hybrid HA microbiota is significantly lower than it is in one progenitor microbiota and significantly higher than it is in the other progenitor microbiota, taxon abundance on the hybrid is intermediate.
- If the relative abundance of the microbial taxon in hybrid HA microbiota is not significantly different from its relative abundance in one or both progenitor microbiota, taxon abundance on the hybrid is conserved.

#### Dietzia maris

An ASV mapping to *Dietzia maris* is the dominant loading (97.5%) on PC1 (49.98% of variation) for gut microbiota and is the dominant loading (97.33%) on PC2 (8.78% of variation) for skin microbiota. The median relative abundance of *Dietzia maris* in gut (see Fig. 4A) and skin (see Fig. 4B) microbiota of hybrid lizards (*A. neomexicanus*) is lower than the median relative abundance of *Dietzia maris* in gut and skin microbiota of either progenitor (*A. marmoratus* at SNW and *A. inornatus* at SBluG). This is true not only when comparing syntopic lizards (*A. neomexicanus* with *A. inornatus* at SBluG or *A. neomexicanus* with *A. marmoratus* at SNW), but also when making comparisons across locations (*A. neomexicanus* at SNW with *A. inornatus* at SBluG or *A. neomexicanus* at SBluG with *A. marmoratus* at SNW). Indeed, in both *A. neomexicanus* gut and skin microbiota, the median relative abundance of *Dietzia maris* is zero, while it is non-zero in both *A. inornatus* gut and skin microbiota and non-zero in both *A. marmoratus* gut and skin microbiota. Thus, the relative abundances of *Dietzia maris* in hybrid gut and skin microbiota are transgressive.

**Figure 4.**
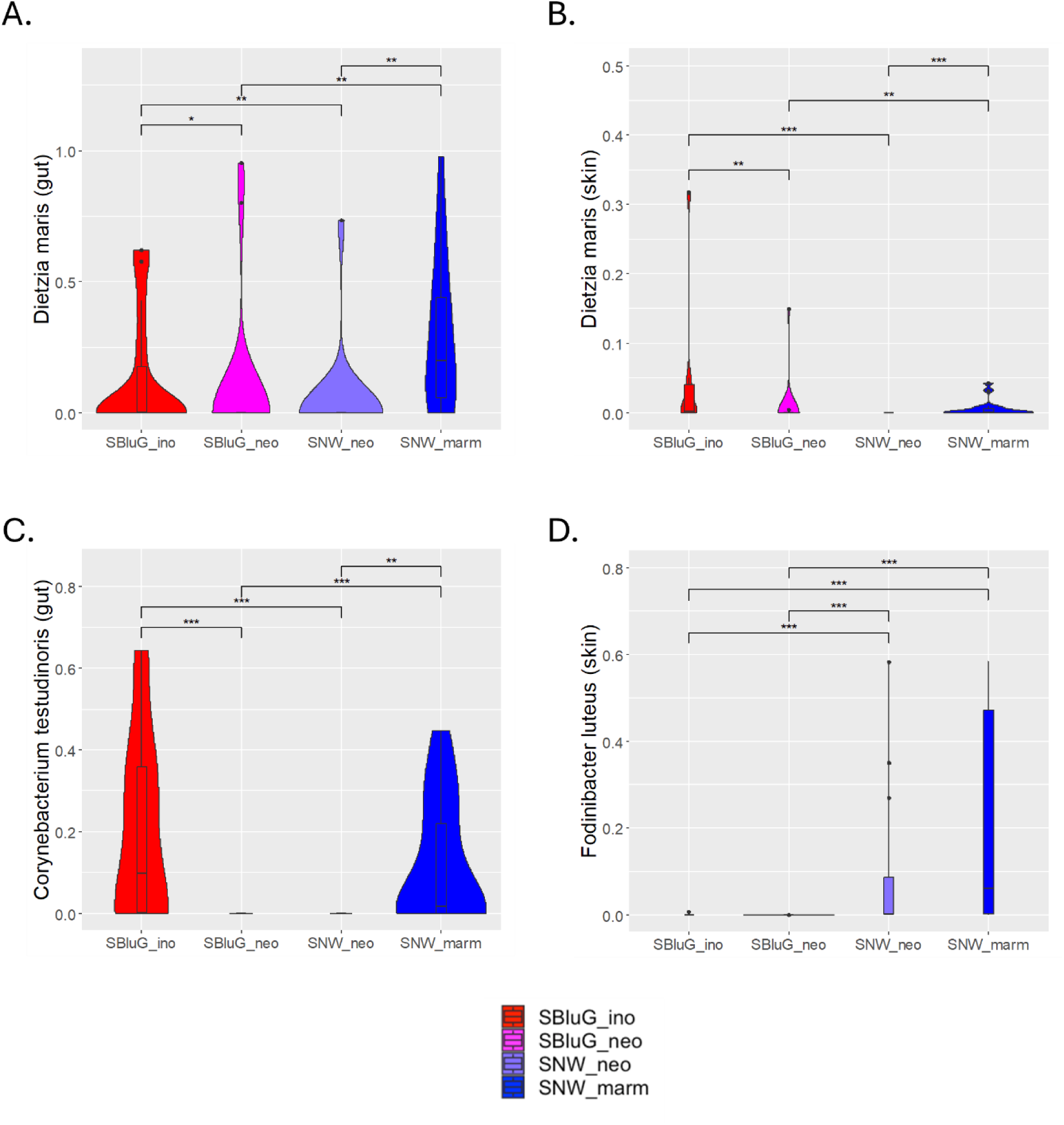
Relative abundances of ASVs representing the dominant loadings on gut and skin Principal Component Axes (PCA). Taxa considered are as follows: *Dietzia maris* in gut (A, dominant loading on PC1) and skin (B, dominant loading on PC2) microbiota, *Corynebacterium testudinoris* in gut microbiota (C, dominant loading on PC2) and *Fodinibacter luteus* in skin microbiota (D, dominant loading on PC1) for Aspidoscelis inornatus from SBluG (red), *A. neomexicanus* from SBluG (magenta), *A. neomexicanus* from SNW (purple), and *A. marmoratus* from SNW (blue).

#### Corynebacterium testudinoris

An ASV mapping to *Corynebacterium testudinoris* is the dominant loading (94.27%) on PC2 (17.53% of variation) for gut microbiota. The median relative abundance of *Corynebacterium testudinoris* in gut microbiota of hybrid lizards (*A. neomexicanus*) is lower than the median relative abundance of *Corynebacterium testudinoris* in gut microbiota of either progenitor (*A. marmoratus* at SNW and *A. inornatus* at SBluG, see Fig. 4C). Again, this is true not only when comparing syntopic lizards (*A. neomexicanus* with *A. inornatus* at SBluG or *A. neomexicanus* with *A. marmoratus* at SNW), but also when making comparisons across locations (*A. neomexicanus* at SNW with *A. inornatus* at SBluG or *A. neomexicanus* at SBluG with *A. marmoratus* at SNW). Similar to *Dietzia maris*, the median relative abundance of *Corynebacterium testudinoris* in gut microbiota of both *A. neomexicanus* populations is zero, while the median relative abundance of *Corynebacterium testudinoris* in gut microbiota of both progenitor species is non-zero. Thus, the relative abundance of *Corynebacterium testudinoris* in hybrid gut microbiota is transgressive.

#### Fodinibacter luteus

An ASV mapping to *Fodinibacter luteus* is the dominant loading (99.15%) on PC1 (8.78% of variation) for skin microbiota. The median relative abundance of *Fodinibacter luteus* is higher in the skin microbiota of both lizard populations from SNW than it is in the skin microbiota of both lizard populations from SBluG (see Fig. 4D). However, there are no statistically significant differences in the relative abundances of *Fodinibacter luteus* between either hybrid population and its syntopic progenitor. This suggests that location effects, rather than lizard species effects, explain why *Fodinibacter luteus* is responsible for variation in our system. Thus, the relative abundance of *Fodinibacter luteus* in hybrid gut microbiota is conserved.

Table 1 summarizes results from microbiota diversity, microbial abundance and microbiota composition traits for each hybrid population. While several traits, particularly those associated with overall microbiota composition, are conserved, the majority of traits are transgressive. This is particularly true for traits related to diversity and abundances of important microbial taxa.

**Table 1.** Summary of results for the various hybrid microbiota traits that we consider. Specifically, a trait can be transgressive (T), intermediate (I), or conserved (C). When a trait is conserved in the hybrid, it can be similar to *A. inornatus* (^I^), similar to *A. marmoratus* (^M^), or similar to both progenitors (^I+M^). When a trait is transgressive, it can be higher than both progenitors (^∧^) or lower than both progenitors (_∨_). The first letter in each column refers to the hybrid population at SBluG. The second letter in each column refers to the hybrid population at SNW.

|  | Gut | Skin |
| --- | --- | --- |
| **Diversity** |  |  |
| α-diversity | $C^{M+I}C^{M+I}$ | $T^{\wedge}C^{M+I}$ |
| β-diversity | $T_v T_v$ | $T_v T_v$ |
| γ-diversity | $T^{\wedge} T^{\wedge}$ | $T^{\wedge}I$ |
| core diversity (absolute) | $T^{\wedge} T^{\wedge}$ | $T^{\wedge} T^{\wedge}$ |
| core diversity (percentage) | $T^{\wedge} T^{\wedge}$ | $C^M C^M$ |
| unique core diversity (absolute) | $T^{\wedge} T^{\wedge}$ | $T^{\wedge} T^{\wedge}$ |
| unique core diversity (percentage) | $T^{\wedge} T^{\wedge}$ | $T^{\wedge} T^{\wedge}$ |
| **Microbial Abundance** |  |  |
| *Dietzia maris* | $T_v T_v$ | $T_v T_v$ |
| *Corynebacterium testudinoris* | $T_v T_v$ | --- |
| *Fodinibacter luteus* | --- | $C^I C^M$ |
| **Composition** |  |  |
| PCoA axis 1 | $C^{M+I}C^M$ | $T^{\wedge}C^{M+I}$ |
| PCoA axis 2 | $C^I C^{M+I}$ | $C^I C^M$ |
| 20D axis of parental variation | $I C^M$ | $II$ |
| 20D height of triangle | $T^{\wedge} T^{\wedge}$ | $T^{\wedge} T^{\wedge}$ |

### Microbiota Restructuring

Summarizing the HA microbiota with one- and two-dimensional metrics (e.g., diversity, microbial abundance, position in ordination space) is convenient for testing our hypotheses that hybrid HA microbiota traits should be distinct from and transgressive to progenitor HA microbiota traits. In reality, however, HA microbiota are highly multidimensional and thus, are often difficult to fully summarize with simple one- and two-dimensional metrics. As a final analysis, we consider some of the more nuanced aspects of microbiota restructuring in the hybrid. This more holistic approach does not permit the simple hypothesis testing that we applied to the traits previously described but provides a summary of some of the subtler relationships between hybrid microbiota and the microbiota of progenitor species. Specifically, we consider a novel metric that we previously introduced - the 4H index^119^ - to describe restructuring of the hybrid microbiota according to four different models for how microbial taxa can be shared amongst progenitors and their hybrid offspring. In Additional files 6 and 7 we also present a description of the general microbiota (see Fig. 5) of each lizard species and identify taxa that are statistically over- and/or under-represented in hybrids relative to their parent species.

**Figure 5.**
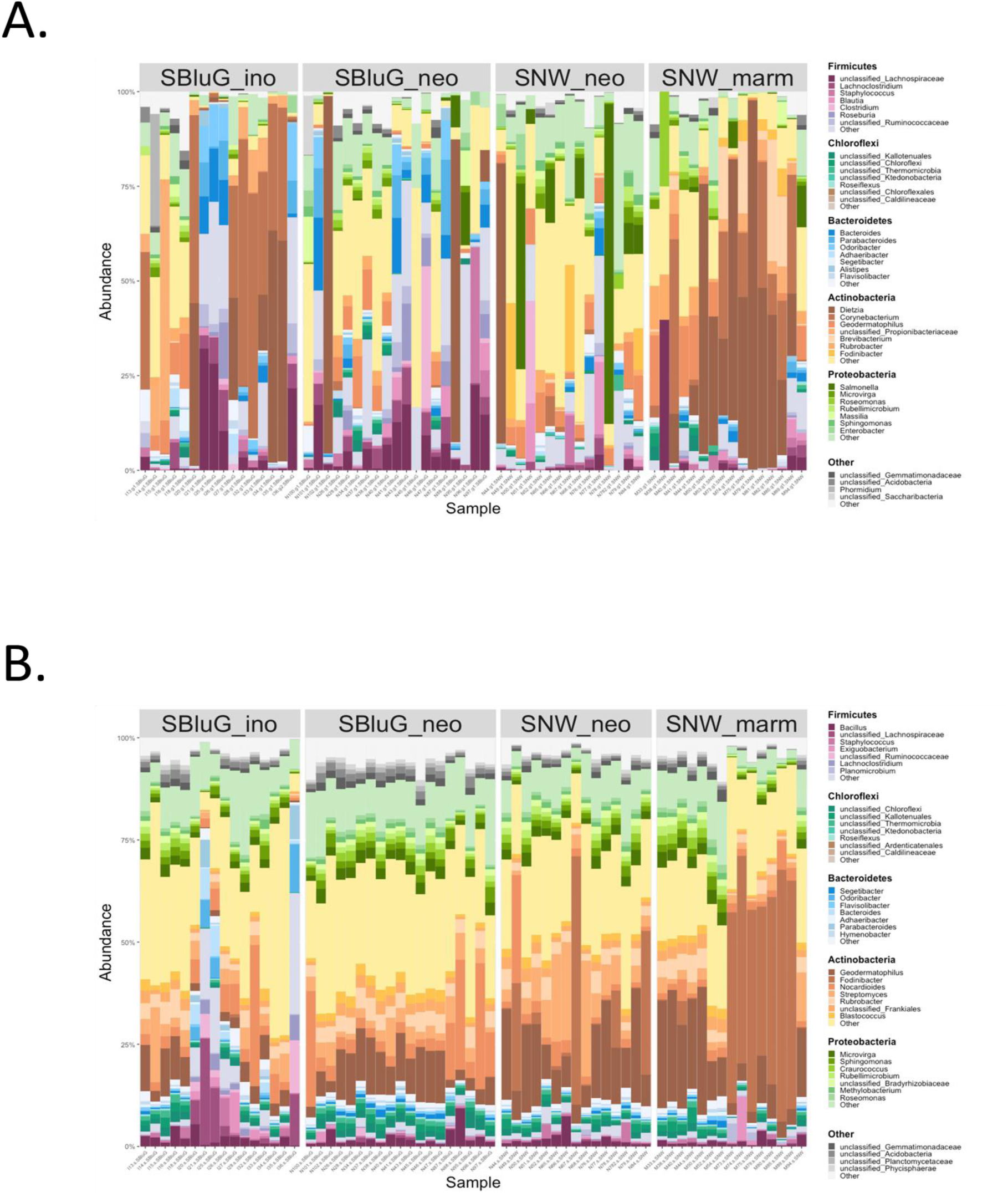
Bar graphs showing the genera composition of the gut (A) and skin (B) microbiota of each individual lizard in each population. See Additional files 6 and 7 for a description of dominant taxa and indicator species analysis.

#### 4H Index

Figure 6 shows our newly developed phylogenetic 4H index applied to the full microbiota (i.e., a core threshold of 0). The 4H index provides a measure of how hybrid microbiota restructuring occurs. In particular, the 4H index quantifies the extent to which hybrid microbiota (i) share microbial phylogenetic branches with the microbiota of both progenitors (Intersection Model), (ii) share microbial phylogenetic branches with the microbiota of one but not both progenitors (Union Model), (iii) are missing microbial phylogenetic branches found in the microbiota of one or both progenitors (Loss Model), and (iv) contain microbial phylogenetic branches not found in the microbiota of either progenitors (Gain Model). As compared to skin microbiota, gut microbiota of hybrid lizards include a higher proportion (see Additional file 8: Table 9.2) of microbial phylogenetic branches found on neither progenitor (Gain Model) and are missing a higher proportion of microbial phylogenetic branches found on one or both progenitors (Loss Model). By contrast, skin microbiota include a higher proportion of microbial phylogenetic branches found on both progenitors (Intersection Model). This results in gut microbiota showing approximately equal support for the ‘Transgressive Axis’ (54%) and the ‘Parental Axis’ (46%), whereas skin microbiota show a much higher support for the ‘Parental Axis’ (67%) relative to the ‘Transgressive Axis’ (33%). Interestingly, hybrid animals are more likely than expected by chance to retain gut and skin microbial phylogenetic branches shared by both progenitors as opposed to microbial phylogenetic branches present on only one progenitor (see Additional file 8 for analysis of other metrics and core thresholds).

**Figure 6.**
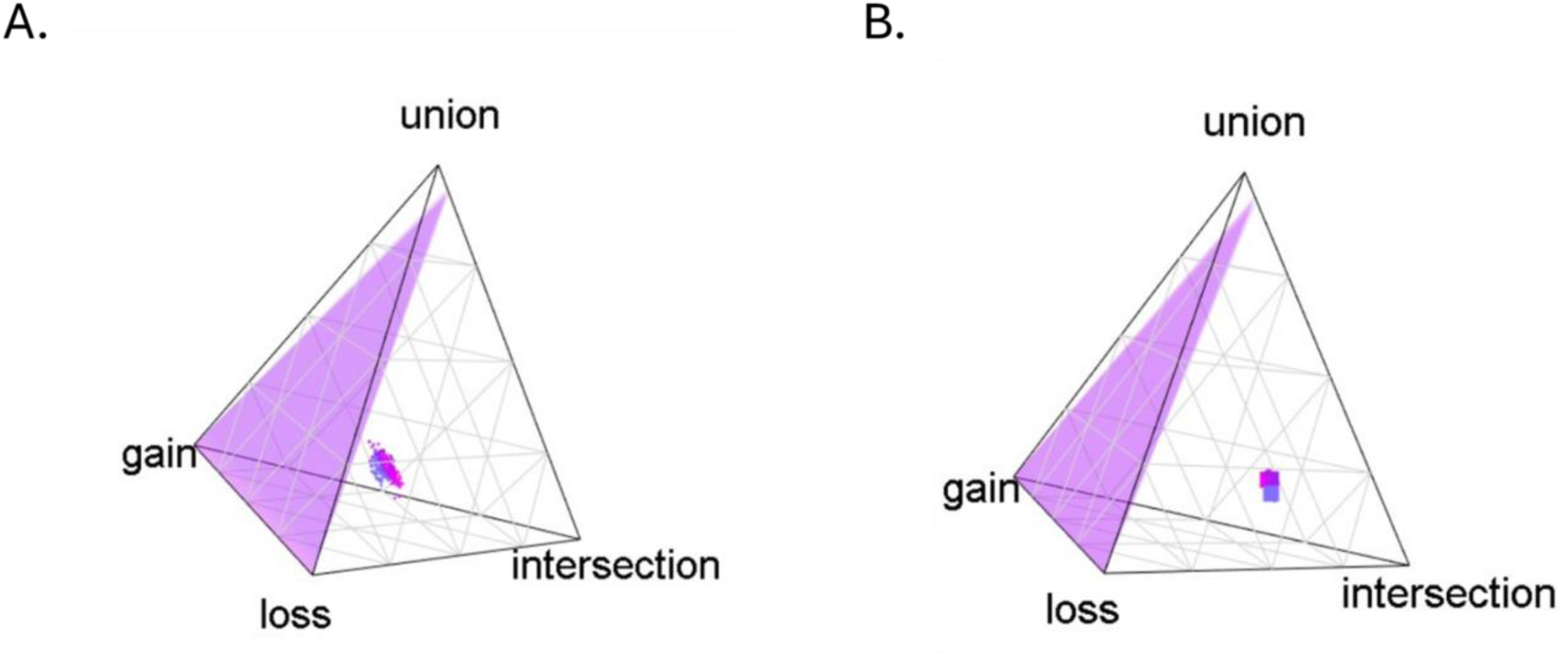
Quaternary plots of the 4H index for branch segments of the gut (A) and skin (B) microbiota of Aspidoscelis neomexicanus at SBluG (magenta), *A. neomexicanus* at SNW (blue-purple) and all *A. neomexicanus* at both locations (purple). The null plane is shown in purple in each plot. For 4H index values, see tables in Additional file 9.

## Discussion

Hybrid parthenogens represent model systems for understanding how hybridization contributes to ecological and evolutionary success. More specifically, because of their clonal reproductive mode, hybrid parthenogens preserve the adaptive variation generated during F_1_ hybridization - arguably the most dramatic stage of the hybridization continuum (i.e., F_1_ hybridization → adaptive introgression → lineage reticulation). As a result, hybrid parthenogens are ideal natural laboratories for exploring the emergence of adaptive variation^120^ (i.e., Goldschmidt’s macromutations^65^) during hybridization and how this impacts ecology and fitness. Indeed, hybrid parthenogens are essentially clones of the ‘hopeful monsters’ that emerged during the F_1_ hybridization event. Especially when utilizing ecologically successful hybrid parthenogens such as *Aspidoscelis neomexicanus*, these simplified systems can be used to explore how macromutations induced by F_1_ hybridization unlock novel ecological space (i.e., new adaptive zones).^121^ While the evolutionary literature certainly appreciates the role of adaptive introgression^122,123^ and lineage reticulation,^124^ the importance of F_1_ hybridization is generally obscured at these deeper scales. Studies on hybrid parthenogens can shed light on the impacts of F_1_ hybridization so that it can be better evaluated as a precursor for adaptive introgression. Ultimately, this will help characterize hybridization’s role in bridging microevolutionary process to macroevolutionary pattern.

By leveraging locations of syntopy between hybrid *Aspidoscelis neomexicanus* and its progenitor species, *A. marmoratus* and *A. inornatus*, we expose the effects of hybridization on HA microbiota in an ecologically successful vertebrate parthenogen. Broadly speaking, hybrid traits can be similar to one or both progenitors (conserved), intermediate to both progenitors, or transgressive. This is true for other types of traits controlled by host genes such as certain aspects of morphology^14–17^ and physiology.^18–22^ It is also true for HA microbiota which, though typically viewed as microbial communities living on or in a host, can also be thought of as multidimensional holobiont traits. Here, we provide evidence supporting our hypotheses that hybrid *A. neomexicanus* gut and skin microbiota are distinct from progenitor microbiota and, for many HA microbiota traits, are transgressive (see Table 1). Importantly, transgressive segregation in host traits is the basis for phenotypic novelty that host evolution can act upon. Our findings suggest that the same may be true for holobiont evolution more broadly.

### Full Microbiota Diversity

Broadly speaking, diversity measures support our hypotheses that hybrid microbiota are both distinct from progenitor microbiota and transgressive. In general, microbial diversity on a single host (*⍺*-diversity) shows the smallest differences between hybrids and their progenitors. These differences are rarely significant for comparisons of individual populations; however, if populations are pooled into a single hybrid class and a single progenitor class, hybrid animals have significantly higher *⍺*-diversity. This suggests that *⍺*-diversity is probably not conserved but rather is transgressive with small effect sizes. Although our study is not designed to determine why hybrid microbiota have higher *⍺*-diversity, our results are consistent with previous findings that have correlated high allelic heterozygosity to high microbiota diversity. In a mouse model, for instance, major histocompatibility complex (MHC) heterozygosity was positively correlated with HA fecal microbiota *⍺*-diversity and function.^125^ Meanwhile, in Seychelles warblers, genome-wide heterozygosity (but not MHC heterozygosity) was positively correlated with gut microbiota diversity.^126^ Notably, hybrid *Aspidoscelis* spp. maintain exceptionally high allelic heterozygosity, upwards of 68% in some species.^110^

The higher *⍺*-diversity of hybrid microbiota is interesting because it could offer alternative explanations for hybrid vigor.^127^ Typically, hybrid vigor is explained as the result of novel allelic combinations produced in newly heterozygous genomes (e.g., in hybrid progeny). Known as the heterosis hypothesis, the assumption is that these novel allelic combinations drive superior trait values such as faster growth rates or greater environmental tolerance through either dominance or overdominance mechanisms.^128^ In the former, hybrid vigor is the result of reducing homozygous alleles, thereby masking deleterious recessive traits. In the latter, hybrid vigor is the result of allelic complementation, wherein organisms benefit from having two different alleles that function in different ways^129,130^ (see Birchler et al.,^128^ Fu et al.,^130^ Kaushik et al.,^131^ and Kochetov et al.^132^ for definitions and discussion of the alternative models for heterosis including more complex mechanisms like intergenic interactions). With respect to holobionts, hybrid vigor could emerge not only through dominance or overdominance in the host genome, but also through dominance- or overdominance in the broader hologenome.^23^ There may, for example, be fitness benefits to inheriting two different microbial metagenomes because functions missing in one metagenome are present in the other (like masking of recessive alleles). Alternatively, there may be benefits to having microbial taxa that function in slightly different ways to perform similar tasks (complementation). Importantly, these ‘holobiont heterosis’ (or ‘hybrid holobiont vigor’) mechanisms would require higher *⍺*-diversity in the hybrid microbial metagenome, which is what we observe.

The higher microbial *⍺*-diversity on individual hybrid lizards is also interesting in light of the general-purpose genotype (GPG) hypothesis. Formulated to explain the paradoxical success of asexual lineages, the GPG hypothesis postulates that successful clones persist because they exhibit greater ecological niche breadth, a more generalist ecological strategy, and/or greater phenotypic flexibility across one or more traits.^133,134^ Typically, this greater flexibility is attributed to differences in epigenetic regulation.^135,136^ Our results, however, suggest an alternative ‘general purpose metagenome’ (GPMG) hypothesis. In particular, if functional diversity parallels taxonomic/phylogenetic diversity (an admittedly tenuous assumption), then the larger and more diverse microbial metagenomes of hybrid lizards may be at least partially responsible for conferring a more generalist ecological strategy and/or greater phenotypic flexibility to hybrid lizards.

Like *⍺*-diversity, *β*-diversity (turnover between lizards or interindividual variation) of the hybrid gut and skin microbiota is also transgressive. However, whereas *⍺*-diversity is higher for hybrid populations, *β*-diversity is lower. Several potential mechanisms could underlie the ‘extreme’ *β*-diversity of hybrids relative to their progenitors. First, lower interindividual variation among hybrids could be explained by Lerner’s Theory.^137^ In particular, the more diverse alleles of more heterozygous individuals might buffer against environmental perturbations, allowing for higher developmental stability. If Lerner’s hypothesis applies to microbial phenotype (e.g., through a more diverse set of host MHC genes that buffer host–microbe interactions), then hybrid lizards should have more ‘stable’ microbiota as compared to progenitors. Second, lower interindividual variation of hybrid lizards could be explained by the Diversity-Stability Theory.^138–141^ Again, this explanation leans on the fact that the genomes of hybrid organisms are, in general, more heterozygous. However, in this case, high host heterozygosity leads to high microbial *⍺*-diversity (see above for examples of how this could occur via MHC genes), and it is this higher *⍺*-diversity that then drives heightened microbiota stability (low microbiota *β*-diversity). Though similar in its underlying driver (i.e., heterozygosity of the host), the Diversity-Stability hypothesis differs from the Lerner’s hypothesis in that stability is not brought about directly by host–microbe interactions, but rather by microbe-microbe interactions^128^ (notice that both Lerner’s hypothesis and the Diversity-Stability hypothesis requires that we make a ‘space for time’ substitution and assume that interindividual variation mimics temporal variation in a single individual). A final explanation for the lower *β*-diversity of hybrid lizards is that they are clones, and thus exhibit relatively low interindividual genetic variation. In this case, it is not the high intraindividual host genetic diversity (i.e., high heterozygosity), but rather the low interindividual host genetic diversity that limits microbial variation among hybrids.^142^ Without further study it is impossible to confidently rule out any of the above three hypotheses for the observed low *β*-diversity of hybrid microbiota. In reality, the true cause could be a combination of these mechanisms or alternative mechanisms altogether. Regardless, the pattern of lower *β*-diversity among parthenogens is an intriguing finding that warrants future investigation.

Population-wide or *γ*-diversity is, in essence, the combination of *⍺*-diversity and *β*-diversity. Similar to *⍺*-diversity, *γ*-diversity of *A. neomexicanus* gut microbiota is transgressive and significantly higher than that of either progenitor species (see Fig. 1 and Additional file 1: Fig. 1.9 and Table 1.1). *γ*-diversity of the *A. neomexicanus* skin microbiota is also higher than that of its syntopic progenitor at both locations (see Fig. 1D) and is transgressive at the SBluG location. However, *γ*-diversity of the *A. neomexicanus* skin microbiota is intermediate to its progenitors at SNW, a finding that we attribute to location-level effects that mask the generally transgressive nature of *A. neomexicanus* skin *γ*-diversity at this location (see ‘Caveats and Future Directions’). Like *⍺*-diversity, the greater *γ*-diversity of hybrid populations has interesting implications for hybrid fitness and ecological success. In particular, it suggests that hybrids not only have greater taxonomic (and putatively functional) diversity at the scale of individual animals (i.e., see GPMG hypothesis above) but also at the scale of entire populations. This is important because it suggests that the greater diversity conferred by interindividual variation (i.e., *β*-diversity) in progenitor populations is outweighed by the greater per individual diversity (i.e., *⍺*-diversity) of hybrids.

### Core Microbiota Diversity

In most ways considered, the hybrid core supports our hypotheses that hybrid HA microbiota traits are both different from progenitors and transgressive. Overall, for both gut and skin microbiota, hybrid animals exhibit higher core diversity and higher diversity of the unique components of their core microbiota (see Additional file 2: Figs. 2.1-2.28). Further, the core gut microbiota of hybrid animals comprise a higher percentage of overall *γ*-diversity, while the unique components of the hybrid core gut and skin microbiota comprise a higher percentage of the hybrid core (see Fig. 2). Like diversity measures, the transgressive nature of the hybrid core microbiota has a number of interesting implications. Whereas higher *⍺*- and/or *γ*-diversity could reflect a weaker immune system (though high microbiome diversity is not strongly positively correlated with immunocompetence in other species^143^), higher core diversity reinforces that at least some of the excess diversity on hybrids is conserved. This makes hybrid diversity less likely to be due to an immune system that is more permissive to environmental exposures. Indeed, hybrids share proportionately more of their overall *γ*-diversity (see Fig. 2B, F), which would not be expected if all the excess diversity reflected transient environmental acquisitions. The higher diversity of the unique core microbiota and the higher percentage of the core that is unique in hybrid animals is also of interest because it suggests that hybrids may have novel holobiont functionality^144,145^ not readily, or at least routinely, observed in their progenitors. Further, it suggests that hybrids may be more distinct from progenitors than progenitors are distinct from each other.

### Microbiota Composition and Microbial Abundances

Many of the metrics that we use to examine microbiota composition in hybrid animals do not support our hypotheses that the hybrid microbiota is transgressive. Rather, depending on the analysis and the axis considered, they suggest that microbiota composition is either conserved or intermediate. The one compositional analysis where we do see transgressive segregation in the hybrids is in the perpendicular axis of our triangle plots. By definition, this axis captures variation that does not differentiate progenitors; thus, it is arguably more likely to detect variation that separates hybrids from both progenitors. Interestingly, even though much of our microbiota composition analysis suggests that composition is conserved or intermediate, the abundances of individual microbial taxa are often strongly transgressive, at least when focusing on the taxa responsible for variation in the system. This indicates a pattern wherein individually transgressive microbial abundances are combined in ways that lead to an overall conserved or intermediate microbiota composition. Although difficult to conceptualize in non-euclidean space (e.g., UniFrac), in a simple PCA this could occur when the abundances of some microbial taxa on hybrids are transgressive and higher than on progenitors while the abundances of other microbial taxa on hybrids are transgressive and lower than on progenitors. Depending on the ordination technique used and the number of axes considered, a scenario like this could result in projecting hybrids onto intermediate locations along ordination axes, even though the hybrids are independently transgressive along all original axes in the system.

The complexities that emerge when attempting to understand a highly multidimensional set of traits using one-dimensional metrics or ordination axes highlight the importance of supplementing hypothesis testing on individual traits with a more holistic approach that can better capture the multidimensional nature of microbiota restructuring. One way to do this is with our newly developed 4H index. Importantly, the 4H index suggests that both gut and skin microbiota exhibit some degree of transgressive segregation, with evidence for both the Gain Model (i.e., hybrids acquire novel microbes not found on either progenitor) and Loss Model (i.e., hybrids are missing microbes found on one or both progenitors). At the same time, we find that there are stark body site differences in how microbiota restructuring occurs. In particular, we see considerably more support for the Gain and Loss models in gut microbiota (see Fig. 6), whereas skin microbiota are overwhelming dominated by the Intersection (i.e., hybrids retain microbes found on both progenitors) model. As a consequence, the restructuring of gut microbiota falls more along the ‘Transgressive Axis’, whereas the restructuring of the skin microbiota falls more along the ‘Parental Axis’ (i.e., conserved or intermediate). Differences in the 4H index across body sites can be used to develop hypotheses for how hybrid HA microbiota might be important to and/or selected for based on host ecology. The two models characterizing the transgressive axis (Gain and Loss) are the foundation for rapid evolutionary change. The Gain Model, in particular, resembles saltation, or the generation of phenotypic novelty (‘hopeful monsters’). By contrast, the parental axis and the Intersection Model, in particular, implies a likeness of hybrid microbiota to progenitor microbiota. This may contribute to ecological success if certain constituents of the progenitors’ skin microbiota that are strongly beneficial for survival, or if the loss of particular microbes disrupts host-microbiome symbioses.^40,146,147^ In this case, retaining progenitor traits may be a requirement for lineage persistence.

### Hybrid Microbiota and the Hybrid Niche

Our findings on *A. neomexicanus* microbiota have a number of interesting implications for holobiont ecology and evolution. In particular, some of the correlations that we observe between the *A. neomexicanus* microbiome and the *A. neomexicanus* niche suggests that there may be a relationship between hybrid microbiome restructuring and hybrid niche restructuring (throughout the remainder of the discussion, we will use ‘microbiome’ even when referring to ‘microbiota’^148^ to indicate the generality of our conclusions, not only to the microbial taxa on/in hosts, but also to the entire microbial ecosystems housed on hosts). Very broadly, there are two reasons why hybrid microbiomes might co-vary with hybrid niche. First, genetic restructuring of the hybrid might directly impact the hybrid microbiome, resulting in altered microbial function that then enables an altered hybrid niche (microbial cause). Second, genetic restructuring of the hybrid might directly impact the hybrid niche, resulting in different exposures to environmental microbes which then alters community reassembly of hybrid microbiomes (microbial effect). In reality, interactions between host niche and the HA microbiome are likely bidirectional,^149,150^ with each affecting aspects of the other. Our study cannot be used to differentiate the directionality of restructuring (i.e., microbial cause = host genetics → host microbiome → host niche in contrast to microbial effect = host genetics → host niche → host microbiome). However, because of what we know about the ecology of *A. neomexicanus* relative to *A. inornatus* and *A. marmoratus*, our study *can* be used to discuss correlations between the hybrid niche and hybrid microbiomes, a first step in determining whether host niche-HA microbiome interactions are occurring. In the following, we relate *A. neomexicanus* niche expansion and niche shift to *A. neomexicanus* microbiome phenotypes.

In general, the higher *⍺*- and *γ*-diversity of hybrid microbiomes suggest a broader or expanded hybrid niche. This is true whether we consider a microbial cause or a microbial effect and is generally consistent with what is known about *A. neomexicanus* ecology, that *A. neomexicanus* is capable of colonizing a wider breadth of habitats than either progenitor (see Additional file 9).^106,114^ Under the microbial cause scenario, it is the greater hybrid microbial diversity that facilitates access to a wider range of environments and/or food sources (this argument assumes that functional diversity parallels taxonomic diversity). By contrast, under the microbial effect scenario, it is the use of a wider range of environments and/or food sources that alters microbiome assembly through exposure to a more diverse set of microbes. While the positive correlation between hybrid microbiome *⍺*/*γ*-diversity and hybrid niche breadth could imply either a microbial cause or a microbial effect, the greater hybrid core diversity preferentially supports a microbial cause. This is because high core diversity implies conservation of excess microbial taxa across entire hybrid populations—a finding that is less likely if excess microbial taxa are purely a result of more diverse environmental exposures (i.e., an expanded hybrid niche). Indeed, for exposures alone to explain our findings, not only would both hybrid populations (higher *γ*-diversity) and individual hybrid animals (higher *⍺*-diversity) have to range over greater environmental variation, but also, individual hybrid animals would have to range over this greater environmental variation in a conserved manner such that they all acquired the same set of identical environmental microbes (higher core diversity). This seems unlikely. That said, exposure to a broader range of environments could explain the portion of excess hybrid *⍺*- and *γ*-diversity that is variable from individual to individual. While excess core diversity generally supports a microbial cause, there is one scenario where environmental exposures (i.e., microbial effect) could explain our findings: if hybrids occur in locations with inherently more diverse (but conserved) environmental microbiomes.

Whereas the excess diversity of the hybrid microbiome suggests niche expansion, the uniqueness of the hybrid microbiome suggests a niche shift. Notably, we see the uniqueness of the hybrid microbiome both in our core microbiome analysis and in our triangle plots that show separation from the progenitors both through some degree of intermediacy along the parental axis and even more so through transgression along the perpendicular axis. PCoA analyses and PERMANOVA analyses (not shown, see Additional file 4: Tables 4.1-4.4) also lend support to the distinctness of the hybrid microbiome, though not for all hybrid–progenitor comparisons (note that PERMANOVA can only test whether hybrid and progenitor microbiota are different, and not whether hybrid microbiota are transgressive). Again, a niche shift is consistent with what we know about *A. neomexicanus* ecology - that *A. neomexicanus* exhibits different dietary^151,152^ and microhabitat preferences^115^ from its progenitors (diel activity pattern differences have also been suggested, though it is less obvious how this type of temporal niche shift might have microbial causes or effects).^103^ As with diversity, the correlation between hybrid microbiome composition and the hybrid niche could reflect either a microbial cause or a microbial effect. Under the microbial cause scenario, it is the compositional restructuring of the hybrid microbiome that allows hybrids to colonize new types of habitat (again, this assumes that altered taxonomic composition is associated with altered functional composition). Novel gut bacteria, for example, could drive differences in prey consumption or efficiency of use in hybrids relative to their progenitors. These changes could then facilitate shifts in diet, shifts in microhabitat preference, or colonization of novel habitat classes (like riparian zones and disturbed habitats for *A. neomexicanus*; see Additional file 9 for analysis and discussion of *A. neomexicanus* habitat niche transgression).^106,114,115,152^ By contrast, under the microbial effect scenario, it is altered habitat associations that drive compositional restructuring of the hybrid microbiome, likely due to altered environmental exposures. Changes in prey consumption or habitat preference, for instance, could result in the hybrid coming into contact with new microbial taxa or different relative abundances of microbial taxa, thereby altering the pool of microbes from which the hybrid microbiome is assembled.

### Implications for Host Ecology and Evolution

Whether ultimately due to a microbial cause or a microbial effect, the concurrent increase in hybrid microbiome diversity and hybrid niche expansion as well as the concurrent shift in hybrid microbiome composition and hybrid niche shift indicate possible relationships between phenotypic novelty in the hybrid microbiome and ecological novelty that has previously been associated with hybrid ecological success. A broader host niche, for example, suggests a competitive advantage for the hybrid.^153^ Meanwhile, a host niche shift indicates potential competitive release from progenitors.^154,155^ Observing potentially beneficial phenotypic novelty in the HA microbiome of our parthenogen model has important implications for how we should view the role of hybridization in holobiont evolution more broadly. More specifically, our findings suggest that transgressive HA microbiome traits (spanning single microbes to compositional phenotypes; see Table 1) may function similarly to transgressive host traits with respect to their impacts on host evolution. Thus, transgressive microbiome traits produced during hybridization could impact host niches in ways that allow hybrid hosts to reach new adaptive zones.^121^ This, in turn, could result in hybrids experiencing greater rates of evolution^156,157^ (potentially leading to adaptive radiations^158,159^) or greater capacity for evolution (increased total diversification, morphological disparity, or ecological disparity^160–162^). If a transgressive microbiome trait enables occupation of a new adaptive zone and an increase in evolutionary capacity, then this trait, despite being of microbial origin, could be considered a host evolutionary/key innovation.^163^ This would not be unprecedented. Indeed, some individual host–microbe symbioses have been previously identified as drivers of adaptive radiations^164,165^ and evolutionary innovations in hosts.^166–168^ Still, the role of entire HA microbiomes in adaptive radiations and key innovations of their host lineages has not been extensively discussed and should be explored as a potential mechanism for host evolution.

Beyond implications for host lineage evolution, hybrid microbiome restructuring could also impact evolution of HA microbes.^169^ More specifically, host hybridization causes the unification of divergent microbial metagenomes. Even in the F_1_ generation, the resulting ‘hybrid microbial metagenomes’ are not necessarily intermediate in genetic compositions (i.e., 50% paternal and 50% maternal) between the two progenitor microbial metagenomes from which they arose. Still, in most cases they likely represent combinations of microbial metagenomic components from both progenitor species. Such microbial metagenome restructuring has the potential to alter microbiome function by introducing novel microbe–microbe interactions, novel host–microbe interactions, or even novel microbes entirely (e.g., the colonization of niches left unoccupied or newly created). Over evolutionary timescales, this microbial metagenome hybridization could contribute to reticulate evolution of the microbial metagenome. Interestingly, because the ratio of progenitor microbial metagenomes does not always parallel the ratio of progenitor genomes in hybrid holobionts, reticulate evolution may occur asymmetrically between the host genome and microbial metagenome. Differences in patterns of gene flow between host genomes and their microbial metagenomes may have significant implications for how we understand holobiont/hologenome evolution, both within the context of host hybridization and more broadly.

Finally, host hybridization could even increase the capacity for evolution of individual microbial genomes. In particular, when microbiomes are suddenly and dramatically reorganized into communities containing novel combinations of microbial taxa, the capacity for microbial novelty within existing microbial taxa, both as a result of altered selection forces imposed by ecological opportunity and via new opportunities for horizontal gene transfer, likely increases. This is consistent with previous applications of the hopeful monster concept^65^ to the rapid evolution of antibiotic resistance due to horizontal gene transfer.^170^ Importantly, over longer timescales, new ecological opportunities and horizontal gene transfer could lead to microbial adaptive radiations,^171,172^ evolutionary/key innovations,^173,174^ and ultimately, reticulate evolution^124^ (i.e. horizontal gene transfer) within the phylogenies of individual microbial taxa.

The simultaneous impact of host hybridization across multiple scales within the hologenome (i.e.., host genome, microbial metagenome, and microbial genome) suggests extensive opportunity for phenotypic novelty, some of which may lead to fitness benefits or niche changes at their respective scale, across multiple scales, or even across the holobiont entirely. Thus, the multiscale nature of holobiont hybridization gives rise to a much wider range of genetic combinations and genetic interactions than are possible with host genomes alone. Despite this, whether and how scales of hybridization interact to generate hopeful holobionts and their implications for eco-evolutionary success remain open questions.

### Caveats and Future Directions

In this study, we explore the effects of hybridization on the HA microbiome in a parthenogenetic vertebrate. Compared to previous studies,^47^ our work has several advantages. First, and foremost, it considers HA microbiomes in an ecologically successful hybrid. Further, it does so in a wild population, allowing microbiomes to reflect both host genetics and, potentially, host environmental associations. As such, our study presents a full, *in situ* depiction of the hybrid microbiota. Finally, by using a vertebrate parthenogen, our study provides a unique lens on hybridization in that it allows us to isolate front-end effects in a frozen F_1_ genotype. Despite the many advantages of our hybrid parthenogen system, a range of extensions to our existing work could prove interesting and help to identify underlying mechanisms driving host–microbe interactions in hybrid organisms. Most obviously, during our field season, we were not able to locate triple syntopy – that is, hybrid *A. neomexicanus* co-occurring with both progenitors simultaneously. This does not impact the majority of our analyses. However, when the hybrid is transgressive at one location and intermediate or conserved at the other, interpretation is less clear. As an example, for skin *γ*-diversity, the hybrid was transgressive at SBluG and intermediate at SNW. Here, the large differences in *γ*-diversity between locations, and the fact that the hybrid was intermediate at the lower *γ*-diversity location and transgressive at the higher *γ*-diversity location suggest that the hybrid was, in fact, transgressive and that lower hybrid diversity at SNW is driven by location-level, rather than species-level effects. As another example, for the first skin UniFrac PCoA axis, the hybrid was transgressive at SBluG and conserved with both progenitors at SNW. Here again, both SBluG populations were higher than both SNW populations, and the centroid for the hybrid SNW population was higher than the centroid for either progenitor population. This suggests that the hybrid was likely transgressive, but the effect size was too small to detect given location-specific effects or even variation within certain locations. While our use of two different locations did not impact the majority of our conclusions, discovery of rare locations of triple syntopy could help to address interpretation challenges. Alternatively, insight could be gained from broader sampling of hybrid and progenitor microbiomes across the full spectrum of habitats where they co-occur.

Another limitation of our study is that, because it involves a wild population, it is observational in nature. Captive studies that house hybrid and progenitor species under identical conditions could help to separate cause from effect as the underlying drivers of microbiome differences between species. Another caveat in our study is that we focus on broad-scale differences in microbiome taxonomy. Without knowing the functional consequences, we are unable to relate the hybrid holobiont traits that we observe with potential ecological impacts on the host. Now that we have demonstrated significant and transgressive changes in the hybrid HA microbiome, including changes that broadly parallel the hybrid niche, it would be interesting to address potential functional changes. Therefore, shotgun metagenomic sequencing or even culture experiments on some of the dominant strains that we observe driving variation across species (e.g., *D. maris*; see Additional files 6 and 7) are important future directions.

A further drawback of our study was that we are unable to differentiate between interindividual and temporal variation in HA microbiomes. Due to low recapture rates, we focus on single time snapshots of individual host HA microbiomes. In Figure 5 (B, D), however, lizards are ordered by date of capture. Clearly, some of the ‘blooms’ of *Dietzia* and *Fodinibacter* in progenitor lizards appear to occur over contiguous time periods. While hybrids were also caught during these time periods, the fact they do not or rarely show this same signature suggests that differences between hybrids and progenitors may be a result of temporal stability and/or resilience of the hybrid microbiome rather than permanent differences in interindividual variation. This could be due to a more rigid hybrid ecological niche that does not respond as much to environmental conditions or due to mechanisms like Lerner’s theory^137^ or Diversity-Stability theory,^138,139^ both of which imply a more temporally robust microbiome. Additional studies targeting recaptures of hybrids and their progenitors could address the differences between interindividual variation and temporal stability within a single individual. In doing so, these studies could lend further support for some of the theories proposed as an explanation for the lower *β*-diversity of hybrid animals.

Another complication of our study is that we assume *A. neomexicanus* represents a frozen F_1_ hybrid and that we are, thus, isolating the front-end effects of hybridization. However, *A. neomexicanus* originated in the Pleistocene, allowing time for the accumulation of genetic mutations.^175^ This could result in the sampled *A. neomexicanus* population exhibiting differentiation from the true F_1_ offspring of the original *A. marmoratus* **x** *A. inornatus* cross. Consequently, we cannot rule out the possibility that additional evolutionary processes beyond hybridization have contributed to the microbiota differences that we observe between our hybrid parthenogens and their progenitors. Further, because parthenogenetic *Aspidoscelis* evolution is constrained by their clonal reproductive mode, these systems are not ideal for investigating every question related to the hybridization continuum and how it connects microevolutionary process to macroevolutionary pattern. For example, hybrid parthenogens, as front-end systems, have limited utility for understanding the role of hybridization at deeper evolutionary scales such as ongoing adaptive introgression^122,123^ and reticulate phylogenetics.^124^ ‘Back-end’ hybrid systems (i.e., those exhibiting considerable backcrossing) by contrast, are much better models for investigating these scales. Indeed, pairing ecologically successful front-end and back-end systems will ultimately be key for unraveling hybridization’s role as a mode of evolution. Fortunately, there are numerous sexually reproducing *Aspidoscelis* spp. that hybridize without forming parthenogens,^176,177^ allowing for the possibility of extending our study to consider front-end and both back-end systems simultaneously.

## Conclusion

We found considerable evidence of transgressive (i.e., novel) phenotypes in both the gut and skin microbiomes of an ecologically successful hybrid, *A. neomexicanus* (see Table 1). Our findings raise important questions about the nature of host–microbiome interactions in hybrid animals and their relevance to the ecological and evolutionary success of the holobiont. Exploring the bidirectional effects of hybridization on host niche and the HA microbiome, as well as teasing apart the microbiome reassembly mechanisms responsible for these relationships, mark exciting future prospects for hybrid microbiome research. Goldschmidt’s lasting legacy was his realization that hopeful monsters have disproportionately positive impacts on evolutionary trajectories. Since Goldschmidt, transgressive segregation has become a widely accepted driver of macrobial adaptive radiation, macrobial evolutionary innovation, and evolutionary diversification more broadly. Holobiont research, however, has largely neglected hybridization as an important driver of evolutionary success. This is because existing studies have found largely negative consequences for host–microbe and microbe–microbe interactions.^47,55,56^ Aligning holobiont research with broader evolutionary paradigms requires understanding that the numerous hybrid holobiont systems with mismatched microbiomes, host–microbe incompatibilities, and/or immune system dysregulation do not negate the importance of hybridization to positive evolutionary outcomes. Rather, host hybridization can still be important because it is the rare instances of ecological success—the hopeful holobionts—that have outsized benefits on holobiont evolution.

## Materials and Methods

### Species Description, Location Description, and Animal Capture

We used a hybrid lizard (*Aspidoscelis* spp., *Teiidae*) system broadly distributed throughout central New Mexico, USA. More specifically, we sampled adults representing the hybrid, obligately parthenogenetic species *Aspidoscelis neomexicanus* where it occurred syntopically with its two progenitor species: *Aspidoscelis inornatus* (♂, also considered *Aspidoscelis arizonae*^94,175^) and *Aspidoselis marmoratus* (♀).^93^ We chose *A. neomexicanus* as our focal hybrid system because it is broadly distributed across diverse habitats, is known to occur syntopically with both progenitor species,^106,114,115^ is of recent hybrid origin, and, to the best of our knowledge, is representative of a single hybrid origin.^175^ Note that we follow taxonomic recommendations by Tucker^178^ and Walker et al.^96^ and use the masculine species epithet for *Aspidoscelis* spp.

All *Aspidoscelis* lizards were captured from two locations within Sevilleta National Wildlife Refuge in Socorro County, New Mexico, USA between late May and early August, 2022. *A. marmoratus* and *A. neomexicanus* occurred syntopically at the ‘Sevilleta Northwest’ (referred herein as ‘SNW’) location (34.397882, -106.867637, Decimal Degrees WGS84), while *A. inornatus* and *A. neomexicanus* occurred syntopically at the ‘Sevilleta Blue Grama’ (referred herein as ‘SBluG’) location (34.331178, - 106.636717, Decimal Degrees WGS84). Lizards were either captured by hand or using drift fences with box funnel traps. All animal bycatch was immediately released from traps upon capture (see Camper et al.^179^ for further location details [where locations SNW=S-NW and SBluG=S-NE] and notes on our trapping methodology). Locations were sampled until we caught at least 15 female lizards of each whiptail species.

### Animal Sampling

Following animal capture, lizards were washed with sterile millipore water for ca. 30 seconds. Lizards were then swabbed (sterile Puritan HydraFlock #253406H; tip style Large; tip dimensions 1.6 cm [0.64 in.] length × 0.55 cm [0.21 in.] diameter) along each major surface 15–20 times (dorsal surface of the cranium, dorsum, laterally in between limbs, dorsal and ventral surfaces of all extremities including tail, venter, and throat) avoiding the mouth and cloaca. Next, the lizards’ cloaca were wiped with a 70% isopropyl ethanol pad for ca. 30 seconds, and a swab (sterile Puritan HydraFlock #253318H; tip style Micro ultrafine; tip dimensions 0.7 cm [0.31 in.] length × 0.3556 cm [0.14 in.] diameter) was inserted into the cloaca and gently twisted and pulsed back-and-forth along the anterior–posterior axis for ca. 30 seconds more. Each swab was deposited in a separate 2 mL DNA/RNA shield tube (Zymo; #R1102) immediately after collection. All lizards were handled using nitrile gloves wiped down with 70% isopropyl alcohol throughout the entire sampling process. We only used female lizards in our data analysis and confirmed their sex by attempting to evert the hemipenes using gentle pressure applied ventrolaterally ca. 0.5 cm posterior to the cloaca. Before releasing sampled lizards 50–100 m away from their capture location (if captured using traps), we marked each lizard with a unique identifier using a battery-powered miniature medical cautery unit (see Additional file 10: Fig. 10.1 regarding our marking scheme). This was done to prevent duplicate sampling of the same lizard.

### 16S rRNA Gene Sequencing

All samples were sent to ZymoBIOMICS Targeted Sequencing Service for Microbiome Analysis (Zymo Research) using 16S rRNA gene sequencing. Below we outline the basic methods that were used. Additional details are available from https://www.zymoresearch.com/pages/16s-its-amplicon-sequencing.

#### DNA Extraction

Gut and skin microbiota samples were extracted using the ZymoBIOMICS®-96 MagBead DNA Kit (Zymo Research, Irvine, CA) using an automated platform.

#### Targeted Library Preparation

The 16S ribosomal RNA gene was targeted using custom primers (*Quick*-16STM Primer Set) to amplify the V3–V4 region. The *Quick*-16S™ NGS Library Prep Kit was used for library preparation and libraries were amplified using real-time PCR. Final PCR products were quantified with qPCR fluorescence readings and pooled based on equal molarity. PCR libraries were cleaned with the Select-a-Size DNA Clean & Concentrator™ and then quantified using TapeStation® (Agilent Technologies, Santa Clara, CA) and Qubit® (Thermo Fisher Scientific, Waltham, WA).

#### Control Samples

A positive control (ZymoBIOMICS® Microbial Community Standard) was used for each DNA extraction and targeted library preparation. Blank or negative controls were used for each extraction and library preparation. In addition to these standard controls, in the field, we collected a series of environmental controls including air (N=2), sterile millipore water (N=1), soil (N=2), box funnel traps (N=2), and a Chevrolet Express 1500 dashboard (N=1). Air samples were collected using the small swab (aforementioned sterile Puritan HydraFlock #253318H), and all other control samples were collected using the large swab (aforementioned sterile Puritan HydraFlock #253406H).

#### Sequencing

An Illumina® MiSeq™ was used to sequence the final library with a V3 reagent kit over 600 cycles. Sequencing was performed with a 10% PhiX spike-in.

#### Bioinformatics Analysis

We used the metadata and biom files provided to us by the ZymoBIOMICS Targeted Sequencing Service (available on our GitHub page: https://github.com/bewicklab/HopefulHybridHolobionts/) for all subsequent analyses. Briefly, Zymo

Research generates these files by inferring unique ASVs from raw reads, removing potential sequencing errors, and removing chimeric sequences using the DADA2 pipeline.^180^ They then assign taxonomy to the dataset using Uclust (Qiime v1.9.1) with reference to the Zymo Research Database—a 16S database curated by Zymo Research. Finally, we used Qiime2 to generate a phylogenetic tree for all microbial ASVs based on the 16S rRNA sequences provided by Zymo Research. The sequence files and script for tree generation are available on our GitHub page. Control samples were not used in analysis but associated FASTQ files will be uploaded to the National Center for Biotechnology Information (NCBI) database upon publication.

### Statistical Analyses

For all analyses, we applied statistical tests to both amplicon sequence variants (ASVs) and 16S rRNA gene sequences pooled to microbial genus. Running analyses at two different taxonomic scales allows for assessment of the effects of taxonomic scale. For ASV analyses, we considered both phylogenetically-aware and phylogenetically-agnostic methods. For genus-level analyses, we only considered phylogenetically agnostic methods. For both ASV and genus-level analyses, we considered both incidence-based and abundance-based approaches. In the main text we present results for analysis of ASVs using phylogenetically aware incidence-based measures. This is because abundance-based methods cannot be leveraged for metrics that calculate overall diversity across multiple animals (e.g., *γ*-diversity, the core microbiota). Further, the phylogenetically aware methods applied to ASVs represent an intermediate scenario that interpolates between focusing on strain-level variation versus focusing on large-scale phylogenetic differences amongst microbiota. Other analyses, however, are presented in the Additional files because these provide different insights into the types of changes that emerge in the microbiota of hybrid animals.

For all analyses, we first removed 16S rRNA gene sequences that could not be identified as Bacteria or Archaea based on the proprietary Zymo database. These sequences were often associated with long branch lengths in the microbial phylogenetic tree and likely represented sequencing artifacts and non-microbial contamination. Additionally, we removed all samples with a read depth lower than 10,000 reads. This resulted in the loss of three gut samples but no skin samples, leaving 19 *A. neomexicanus* at SBluG, 15 *A. neomexicanus* at SNW, 16 *A. marmoratus* at SNW, and 16 *A. inornatus* at SBluG. Finally, we rarefied all samples to the read depth of the lowest sample from each particular body site (13126 reads for gut samples, 116931 reads for skin samples) using the rarefy_even_depth function from the phyloseq package.^181^ All analyses were performed using the R programming language (version 4.2.1)^182^ available on our Github page (see ‘Bioinformatics Analysis’).

#### *⍺*-diversity

For ASVs, we considered richness (i.e., a count of ASVs), Shannon diversity, Simpson diversity, Faith’s phylogenetic diversity (PD), and Rao’s quadratic entropy. For microbial genera, we considered richness, Shannon diversity, and Simpson diversity. Richness was calculated by counting microbial presences in the ASV and genus tables for each sample. Shannon and Simpson diversities were calculated using the diversity function in the vegan package (version 2.6.4).^183^ Faith’s PD was calculated using the pd function in the picante package (version 1.8.2).^184^ Rao’s quadratic entropy was calculated using the raoD function in the picante package. For Faith’s PD, we used a rooted, non-ultrametric microbial phylogeny. However, because the raoD function requires an ultrametric tree, for Rao’s quadratic entropy, we converted the microbial phylogeny into an ultrametric tree by first removing multichotomies using the multi2di function and then applying molecular dating with mean path lengths using the chonoMPL function, both from the ape package (version 5.7.1).^185^ To identify significant differences in *⍺*-diversity between groups, we first used a Kruskal-Wallis test applied to the four distinct populations (*A. inornatus* at SBluG, *A. neomexicanus* at SBluG, *A. neomexicanus* at SNW, and *A. marmoratus* at SNW). This was done using the kruskal.test function from the stats package (version 4.2.1).^182^ When the Kruskal-Wallis test was significant, we did pairwise post-hoc testing by applying pairwise Wilcoxon rank sum tests with a Benjamini-Hochberg correction using the pairwise.wilcox.test function, also from the stats package. In addition to testing for diversity differences between the four populations separately, we also tested for diversity differences between asexual (both hybrid populations) and bisexual (both progenitor populations) lizards. For asexual/bisexual comparisons, all statistical tests were performed as described above, except that we used a Mann-Whitney U test in place of a Kruskal-Wallis test. This was done with the wilcox.test function from the stats package.

#### *β*-diversity

For all estimates of *β*-diversity, we used multisite dissimilarity indices.^186,187^ We chose this approach over pairwise or averaged pairwise dissimilarity indices because multisite dissimilarity indices avoid pseudoreplication (a problem with pairwise metrics) and account for higher order co-occurrences (a problem with averaged pairwise metrics). Thus, multisite dissimilarity indices better represent dissimilarity across larger (>3) sets of communities.^186^ For ASVs we considered Jaccard, Bray-Curtis, and unweighted UniFrac dissimilarity indices. We did not consider any abundance-based phylogenetically aware metrics because these are not currently available as part of the betapart package (version 1.6).^188^ For microbial genera, we considered Jaccard and Bray-Curtis dissimilarity indices. To test for significant differences between groups, we used a resampling approach. Specifically, for each of our four distinct populations (*A. inornatus* at SBluG, *A. neomexicanus* at SBluG, *A. neomexicanus* at SNW, and *A. marmoratus* at SNW) we calculated the multisite *β*-diversity of 15 different subsamples, with each subsample consisting of 12 randomly selected animals from the focal population. *β*-diversity was calculated using the beta.multi, the beta.multi.abund, and the phylo.beta.multi functions from the betapart package for Jaccard, Bray-Curtis, and unweighted UniFrac dissimilarities respectively. To assess the significance of differences between populations, we applied Kruskal-Wallis tests followed by pairwise post-hoc testing with a Benjamini-Hochberg correction (see *⍺*-diversity section for details).

#### *γ*-diversity

For *γ*-diversity, we again used a resampling approach, selecting 15 subsamples from each population, with each subsample consisting of 12 animals (see *β*-diversity for details). For ASVs, we calculated the overall *γ*-diversity based on richness (i.e., a count of all of the microbial taxa present across all 12 animals) and Faith’s PD. For microbial genera, we only considered richness. Similar to *⍺*-diversity, we used the pd function in the picante package to calculate Faith’s PD. The significance of differences between populations was assessed using Kruskal-Wallis tests followed by pairwise post-hoc testing with a Benjamini-Hochberg correction (see *⍺*-diversity section for details).

#### Core Diversity

Similar to *β*- and *γ*-diversity, we used a resampling approach to examine the properties of the core microbiota for each lizard population. As before, each subsample consisted of 12 randomly selected animals, and 15 subsamples were used for each population. In the main text, we focused on the core microbiota based on a 50% threshold, meaning that a microbial taxon had to be present on 50% of hosts (6 hosts) in a population to be considered core. However, we also evaluated the effect of changing this threshold from 8%–100% (1–12 hosts). For ASVs, we calculated the richness (i.e., a count of all the microbial taxa present) and the Faith’s PD of the core microbiota, and for genera we calculated the richness (i.e., a count of all the microbial genera present). We also determined what percentage of the overall *γ*-diversity was represented in the core by dividing core diversity by *γ*-diversity at each core threshold considered. Also, we calculated the ASV richness, Faith’s PD, and genus richness of the unique component of the core microbiota of each population. For the two progenitor populations, ‘unique’ was defined as any microbial taxon found only on that population. For the hybrid populations, ‘unique’ was defined as any microbial taxon not found on either progenitor population, regardless of whether or not it was present on the other hybrid population. In addition to taxa that were unique to the core of each species and unique to the core of hybrids versus progenitors, we also considered taxa that were part of the core of one species (or both progenitors) that were exclusive to that species (or exclusive to progenitors). These are species that were not found at all on other species (or on hybrids). For this analysis we only considered a core threshold of 50%. Finally, we calculated the percentage of the core microbiota that was unique to any given population by dividing the richness/diversity of the unique component of the core microbiota by the richness/diversity of the entire core microbiota of that population. The significance of differences between groups was again assessed using Kruskal-Wallis tests followed by pairwise post-hoc testing with a Benjamini-Hochberg correction (see *⍺*-diversity section for details). Venn diagrams were similarly made by defining the core microbiota as any microbial taxon that was part of the microbiota of at least 50% of hosts of any given population. Venn diagrams were generated based on the entire dataset (i.e., not using resampling) using the venn.diagram function in the VennDiagram package (version 1.7.3).^189,190^

#### Triangle Plots & Ordination

Triangle plots amd Principal Coordinate Analysis (PCoA) were performed on Euclidean, Jaccard, Bray-Curtis, unweighted UniFrac, and weighted Unifrac distances for ASVs and Euclidean, Jaccard, and Bray-Curtis distances for microbial genera. Triangle plots and associated null models were generated using the TriangleHbootstrap, and TriangleHnull functions from the HybridMicrobiomes package. Briefly, triangle plots are generated by locating the progenitor centroids and the centroid of the hybrid population(s), finding the projection of the hybrid centroid onto the axis connecting the parental centroids (i.e., ‘position along the main axis of parental variation’), and then finding the perpendicular distance between the parental axis and the hybrid centroid (i.e., ‘triangle height’). For abundance-based metrics (i.e., Euclidean, Bray-Curtis, and weighted UniFrac), we generated a hybrid null model by averaging microbial abundances from pairs of randomly selected *A. marmoratus* and *A. inornatus* individuals. For incidence-based metrics (i.e., Jaccard and unweighted UniFrac), we generated a null model by selecting a random pair of *A. marmoratus* and *A. inornatus,* recording the presence of microbial taxa on both progenitors and randomly recording half of the presences of microbial taxa found on either progenitor but not both. Progenitor models were generated by randomly sampling individuals from the *A. inornatus* and *A. marmoratus* populations. For more details, see function descriptions in the HybridMicrobiomes package.^119^ PCoA was performed using the pcoa function from the ape package.^185^ For triangle plots, differences between hybrid and progenitor microbiota were assessed using the TriangleHcompare function individually on the x-axis (position) and y-axis (height). For PCoA analyses, differences between hybrid and progenitor microbiota were assessed by applying Mann-Whitney U tests to the hybrid and progenitor populations separately based on position along the first and second principal coordinate axes.

#### 4H Index

Quaternary plots were generated by defining the core microbiota as any microbial taxon or (for the UniFrac-inspired 4H metric) microbial phylogenetic branch segment that was part of the microbiota of a fixed number of hosts of any given population. For the main paper, we consider a core threshold of 0% (i.e., the full microbiome). However, we also present analysis using a 50% core threshold in Additional file 8. For Jaccard-inspired 4H indices, we used the FourHbootstrap, FourHquaternary, and FourHnullplane functions from the HybridMicrobiomes package (version 0.1.2).^119^ Because the HybridMicrobiomes package does not include a UniFrac-inspired 4H index, we developed this for the current study. Briefly, using the phylogenetic tree for the entire system (i.e., the microbiota of both progenitors and the hybrid) a branch segment in the phylogenetic tree was determined to be part of the core of a particular population or species if it was present on at least the threshold number of animals of that population/species. Once core branch segments had been determined for all populations/species, the populations were compared. Intersection was taken as the fraction of the total branch length shared between the hybrid and both progenitors. Union was taken as the fraction of the total branch length shared between the hybrid and one but not both progenitors. Gain was taken as the fraction of the total branch length only found on the hybrid, and Loss was taken as the fraction of the total branch length not found on the hybrid but found on at least one progenitor. This is analogous to our previous definition of the 4H index but uses branch lengths instead of taxon counts. To fully characterize the range of possible outcomes for the 4H index, we used 100 subsamples of 12 animals from each population, with the hybrid populations either including only the animals from SBluG, only the animals from SNW, or both. In order to ensure comparability between gut and skin microbiota, we rarefied all samples to the lowest read depth present in our gut dataset.^191^

## Declarations

### Ethics approval and consent to participate

All research was approved by Clemson University under IACUC protocol numbers #2020-015 and #2021-047. We completed this work under the Sevilleta National Wildlife Refuge Special Use Permit #SEV_Bewick_Camper_2022_59 and the New Mexico Department of Game and Fish permit authorization #3772.

### Consent for publication

not applicable

### Availability of data and materials

All sequence data will be deposited in the National Center for Biotechnology Information (NCBI) database upon acceptance. BIOM and metadata files are included in Additional file 11. Additionally, BIOM files, metadata files, and all code necessary for the analyses presented in this manuscript are available at https://github.com/bewicklab/HopefulHybridHolobionts/.

### Authors’ contributions

BTC and SAB conceived the idea, performed the data analysis, and wrote the first draft of the manuscript. BTC, ASK, ZTL, and RTM performed all related fieldwork. All authors contributed substantially to the content of the manuscript.

## Acknowledgements

We thank Thomas Dempster, August Spencer, and Eva Purcell for their field assistance in New Mexico as well as Lily Margeson, Simon Dunn, Georgianna Bellinger, Henry Egloff, Kaila Hodges, Camryn Lachica, and Savannah Utz for their assistance assembling drift fence trapping arrays.

