## Additional file 8 for "Transgressive Hybrids as Hopeful Holobionts"

**Additional File 8: 4H Index Analyses**

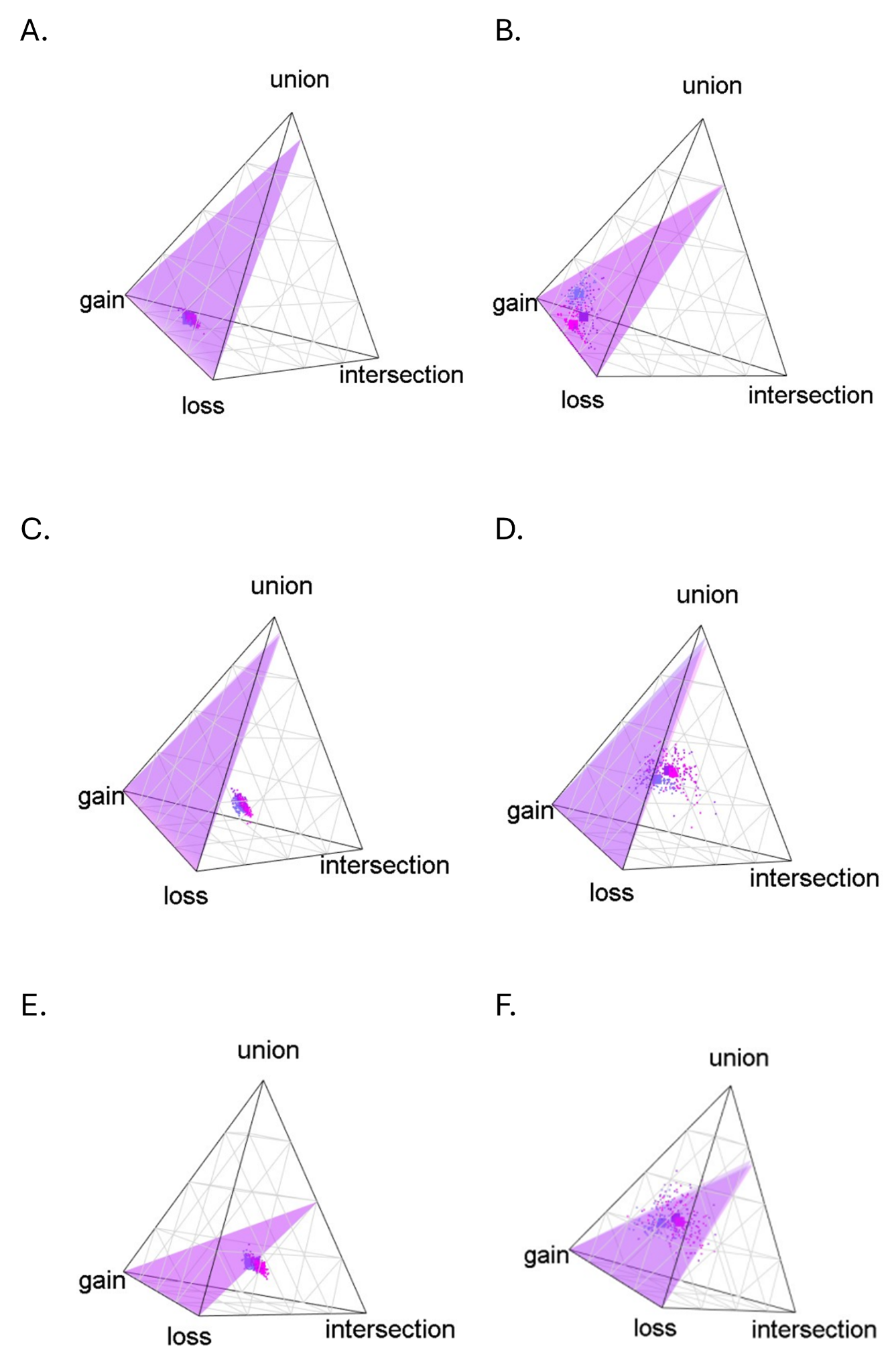

**Figure 8.1** Quaternary plots for skin microbiota assuming a 0% core threshold (full, microbiota, A,C and E) and a 50% core threshold (B,D and F). Here we use a Jaccard-inspired metric on ASVs (A,B), a UniFrac-inspired metric on ASVs (C,D) and a Jaccard-inspired metric on genera (E,F).

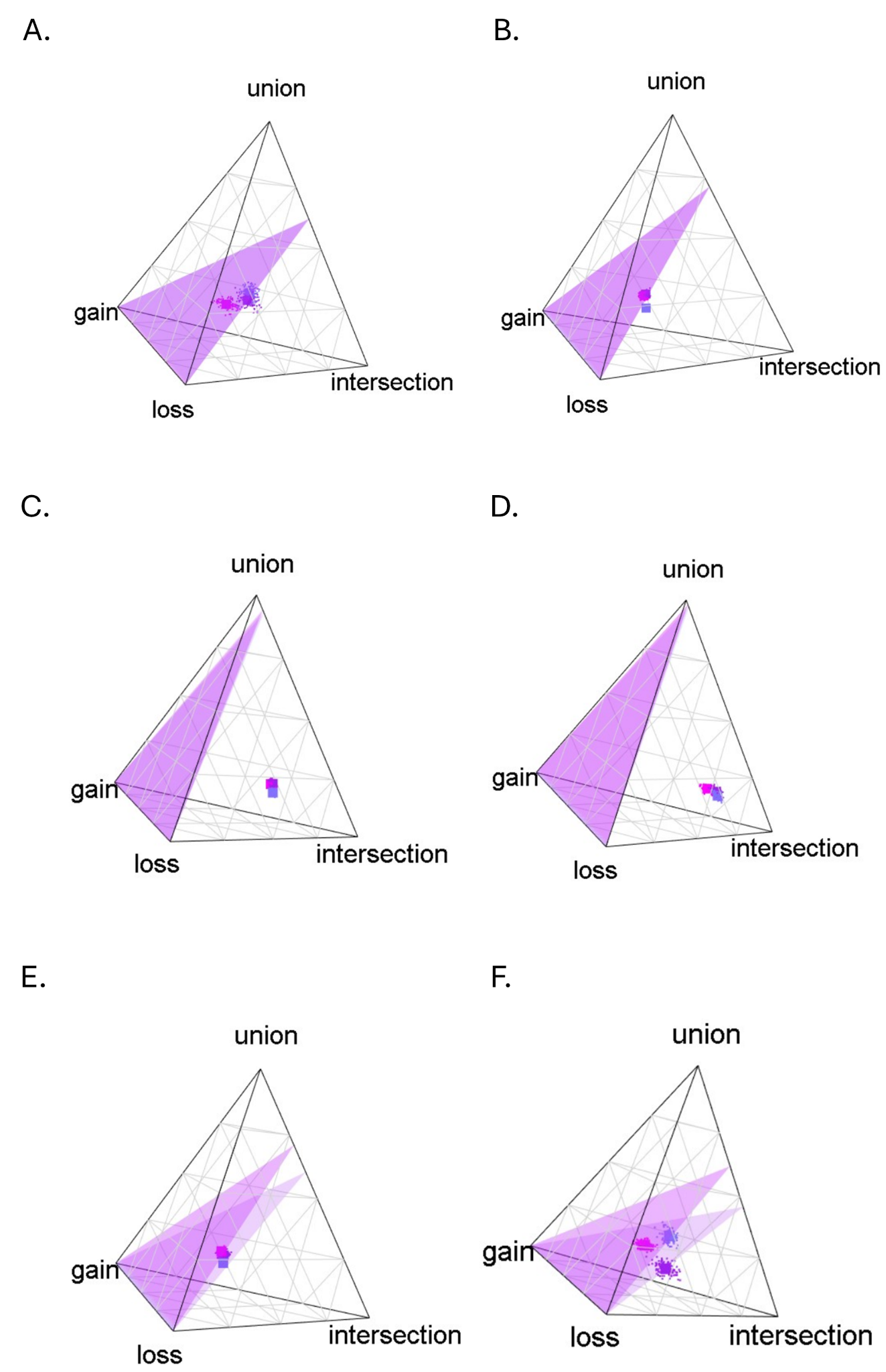

**Figure 8.2** Quaternary plots for gut microbiota assuming a 0% core threshold (full, microbiota, A,C and E) and a 50% core threshold (B,D and F). Here we use a Jaccard-inspired metric on ASVs (A,B), a UniFrac-inspired metric on ASVs (C,D) and a Jaccard-inspired metric on genera (E,F).

**Table 8.1.** The four dimensions (intersection, union, gain, and loss) of the 4H metric for gut and skin microbiota of *Aspidoscelis neomexicanus* at SBluG, *A. neomexicanus* at SNW, and *A. neomexicanus* overall using a Jaccard-inspired 4H metric on ASVs and a core threshold of 0%.

|  | intersection | union | Parental axis | gain | loss | Transgressive axis |
| --- | --- | --- | --- | --- | --- | --- |
| Gut ASV (both) | 0.291976 | 0.167519 | 0.459495 | 0.2355 | 0.305005 | 0.540505 |
| Gut ASV (SNW) | 0.288655 | 0.14596 | 0.434615 | 0.242212 | 0.323173 | 0.565385 |
| Gut ASV (SBluG) | 0.301845 | 0.156991 | 0.458836 | 0.206907 | 0.334257 | 0.541164 |
| Skin ASV (both) | 0.191526 | 0.24807 | 0.439596 | 0.182442 | 0.377961 | 0.560404 |
| Skin ASV (SNW) | 0.193422 | 0.202381 | 0.395803 | 0.144459 | 0.459738 | 0.604197 |
| Skin ASV (SBluG) | 0.187028 | 0.240872 | 0.4279 | 0.187098 | 0.385002 | 0.5721 |
| Gut Genus (both) | 0.352039 | 0.18224 | 0.534279 | 0.206532 | 0.259189 | 0.465721 |
| Gut Genus (SNW) | 0.339219 | 0.177576 | 0.516795 | 0.240319 | 0.242886 | 0.483205 |
| Gut Genus (SBluG) | 0.362021 | 0.169939 | 0.53196 | 0.164889 | 0.303151 | 0.46804 |
| Skin Genus (both) | 0.227088 | 0.212734 | 0.439822 | 0.232683 | 0.327495 | 0.560178 |
| Skin Genus (SNW) | 0.193313 | 0.203594 | 0.396907 | 0.143356 | 0.459737 | 0.603093 |
| Skin Genus (SBluG) | 0.186878 | 0.23976 | 0.426638 | 0.18652 | 0.386842 | 0.573362 |

**Table 8.2.** The four dimensions (intersection, union, gain, and loss) of the 4H metric for gut and skin microbiota of *Aspidoscelis neomexicanus* at SBluG, *A. neomexicanus* at SNW, and *A. neomexicanus* overall using a UniFrac-inspired 4H metric on ASVs and a core threshold of 0%.

|  | intersection | union | Parental axis | gain | loss | Transgressive axis |
| --- | --- | --- | --- | --- | --- | --- |
| Gut ASV (both) | 0.291976 | 0.167519 | 0.459495 | 0.2355 | 0.305005 | 0.540505 |
| Gut ASV (SNW) | 0.288655 | 0.14596 | 0.434615 | 0.242212 | 0.323173 | 0.565385 |
| Gut ASV (SBluG) | 0.301845 | 0.156991 | 0.458836 | 0.206907 | 0.334257 | 0.541164 |
| Skin ASV (both) | 0.466235 | 0.202241 | 0.668476 | 0.102447 | 0.229077 | 0.331524 |
| Skin ASV (SNW) | 0.467406 | 0.170353 | 0.637759 | 0.078168 | 0.284073 | 0.362241 |
| Skin ASV (SBluG) | 0.459918 | 0.199827 | 0.659746 | 0.109123 | 0.231132 | 0.340254 |

**Table 8.3.** The four dimensions (intersection, union, gain, and loss) of the 4H metric for gut and skin microbiota of *Aspidoscelis neomexicanus* at SBluG, *A. neomexicanus* at SNW, and *A. neomexicanus* overall using a Jaccard-inspired 4H metric on ASVs and a core threshold of 50%.

|  | intersection | union | Parental axis | gain | loss | Transgressive axis |
| --- | --- | --- | --- | --- | --- | --- |
| Gut ASV (both) | 0 | 0.107366 | 0.107366 | 0.3525 | 0.540134 | 0.892634 |
| Gut ASV (SNW) | 0 | 0.157988 | 0.157988 | 0.50857 | 0.333442 | 0.842012 |
| Gut ASV (SBluG) | 0 | 0.05508 | 0.05508 | 0.423589 | 0.521331 | 0.94492 |
| Skin ASV (both) | 0.288093 | 0.243767 | 0.531859 | 0.185853 | 0.282288 | 0.468141 |
| Skin ASV (SNW) | 0.286947 | 0.273862 | 0.560809 | 0.18218 | 0.257011 | 0.439191 |
| Skin ASV (SBluG) | 0.244858 | 0.199172 | 0.44403 | 0.294612 | 0.261357 | 0.55597 |
| Gut Genus (both) | 0.166464 | 0.32791 | 0.494374 | 0.316415 | 0.189211 | 0.505626 |
| Gut Genus (SNW) | 0.148701 | 0.283391 | 0.432092 | 0.402685 | 0.165223 | 0.567908 |
| Gut Genus (SBluG) | 0.178326 | 0.315567 | 0.493893 | 0.283788 | 0.222319 | 0.506107 |
| Skin Genus (both) | 0.357385 | 0.127176 | 0.484562 | 0.204795 | 0.310643 | 0.515438 |
| Skin Genus (SNW) | 0.293998 | 0.268983 | 0.562981 | 0.178201 | 0.258818 | 0.437019 |
| Skin Genus (SBluG) | 0.247156 | 0.199134 | 0.446289 | 0.28939 | 0.264321 | 0.553711 |

**Table 8.4.** The four dimensions (intersection, union, gain, and loss) of the 4H metric for gut and skin microbiota of *Aspidoscelis neomexicanus* at SBluG, *A. neomexicanus* at SNW, and *A. neomexicanus* overall using a UniFrac-inspired 4H metric on ASVs and a core threshold of 50%.

|  | intersection | union | Parental axis | gain | loss | Transgressive axis |
| --- | --- | --- | --- | --- | --- | --- |
| Gut ASV (both) | 0.262693 | 0.323627 | 0.58632 | 0.309406 | 0.104274 | 0.41368 |
| Gut ASV (SNW) | 0.246477 | 0.269075 | 0.515552 | 0.37496 | 0.109488 | 0.484448 |
| Gut ASV (SBluG) | 0.277674 | 0.322314 | 0.599988 | 0.28124 | 0.118772 | 0.400012 |
| Skin ASV (both) | 0.612187 | 0.162919 | 0.775106 | 0.098164 | 0.12673 | 0.224894 |
| Skin ASV (SNW) | 0.615627 | 0.148881 | 0.764508 | 0.082369 | 0.153123 | 0.235492 |
| Skin ASV (SBluG) | 0.58287 | 0.162787 | 0.745657 | 0.144062 | 0.110281 | 0.254343 |
