## Additional file 2 for "Transgressive Hybrids as Hopeful Holobionts"

**Additional File 2: Core Microbiota Analyses**

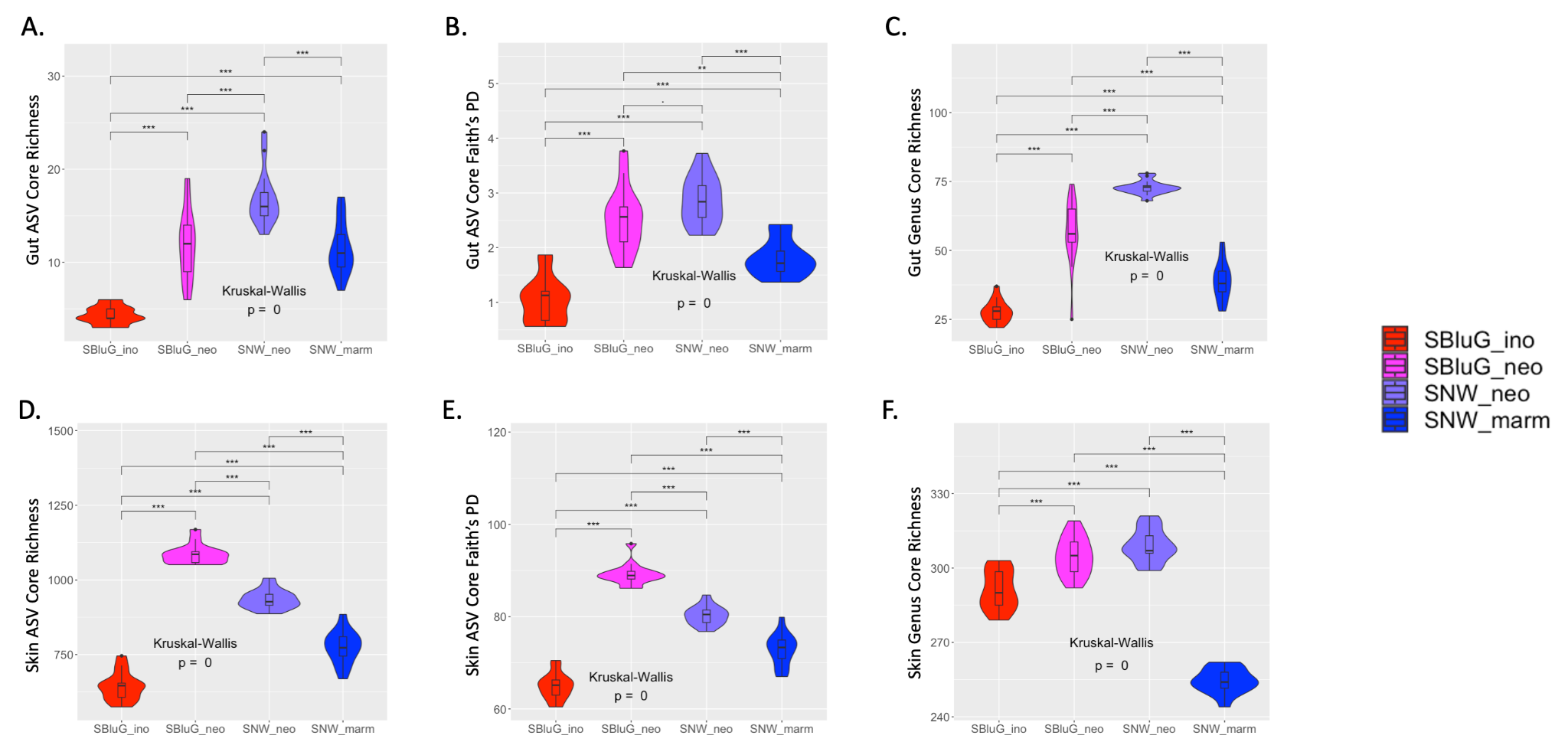

**Figure 2.1.** Comparison of the diversity of the core gut (A-C) and skin (D-F) microbiota between populations of *Aspidoscelis inornatus* from SBluG (red), *A. neomexicanus* from SBluG (magenta), *A. neomexicanus* from SNW (purple), and *A. marmoratus* from SNW (blue) as measured using (A, D) richness of amplicon sequence variants (ASV count), (B, E) Faith’s phylogenetic diversity (PD) of ASVs, and (C, F) richness of microbial genera (genera count). Significant differences in diversity between groups, as determined by a Kruskal-Wallis test followed by post hoc pairwise Wilcox tests using a Benjamini-Hochberg correction, are indicated as follows: p-value ≤ 0.001 (***), p-value ≤ 0.01 (**), p-value ≤ 0.05 (*), p-value ≤ 0.1 (.). In all panels, core microbiota diversity estimates were generated using 15 subsamples of 12 individuals from each population and core microbiota were determined at the 50% threshold (i.e., must be present on 6 hosts to be classified as part of the core).

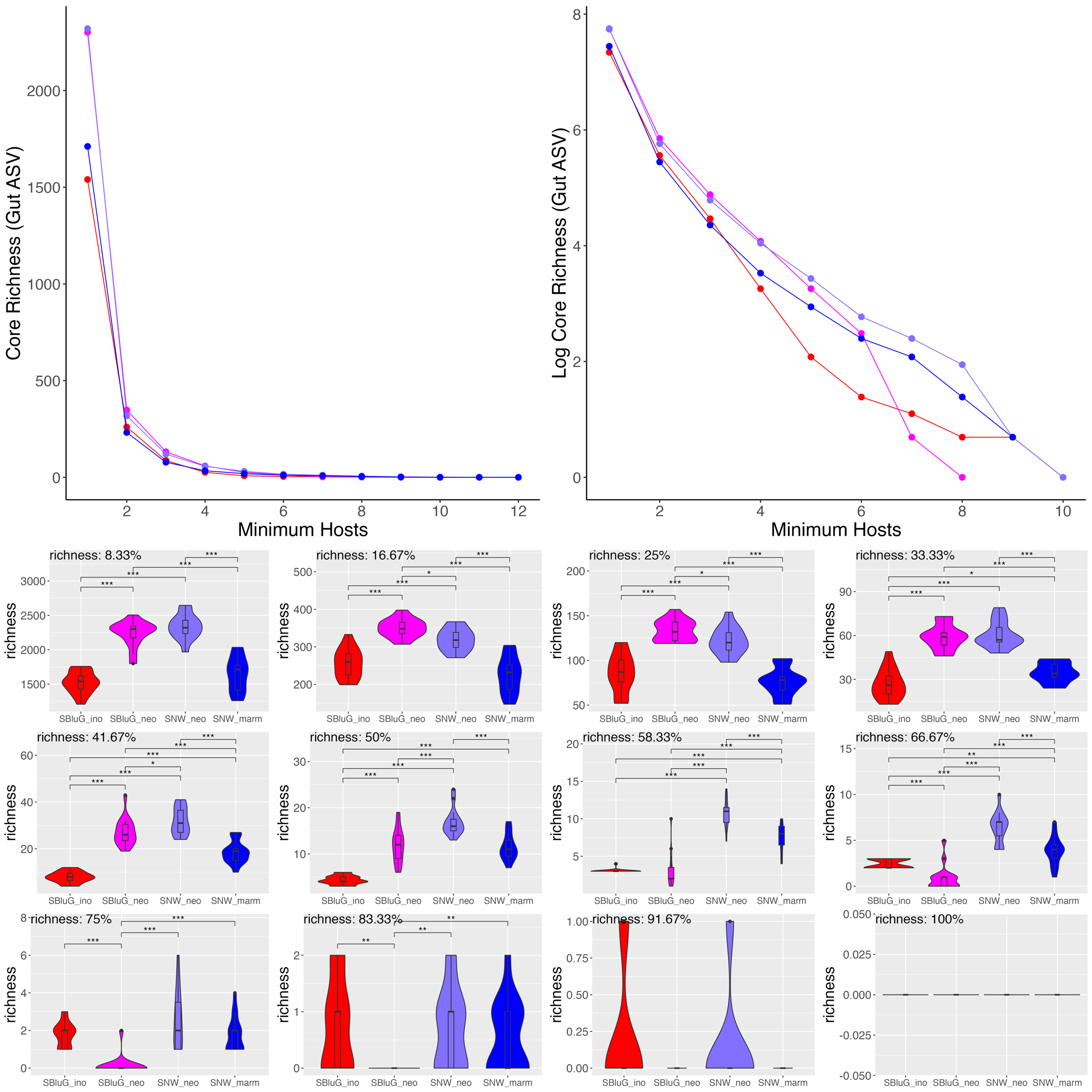

**Figure 2.2.** Richness of amplicon sequence variants (ASV count) for core gut microbiota as a function of the core threshold (i.e., the number of hosts an ASV must be present on in order to be classified as part of the core). In all panels, core microbiota richness estimates were generated using 15 subsamples of 12 individuals from each population and colors, statistical tests and symbols are as in Fig. 2.1. The top two panels show core richness as a function of core threshold. The bottom twelve panels show violin plots for each particular core threshold.

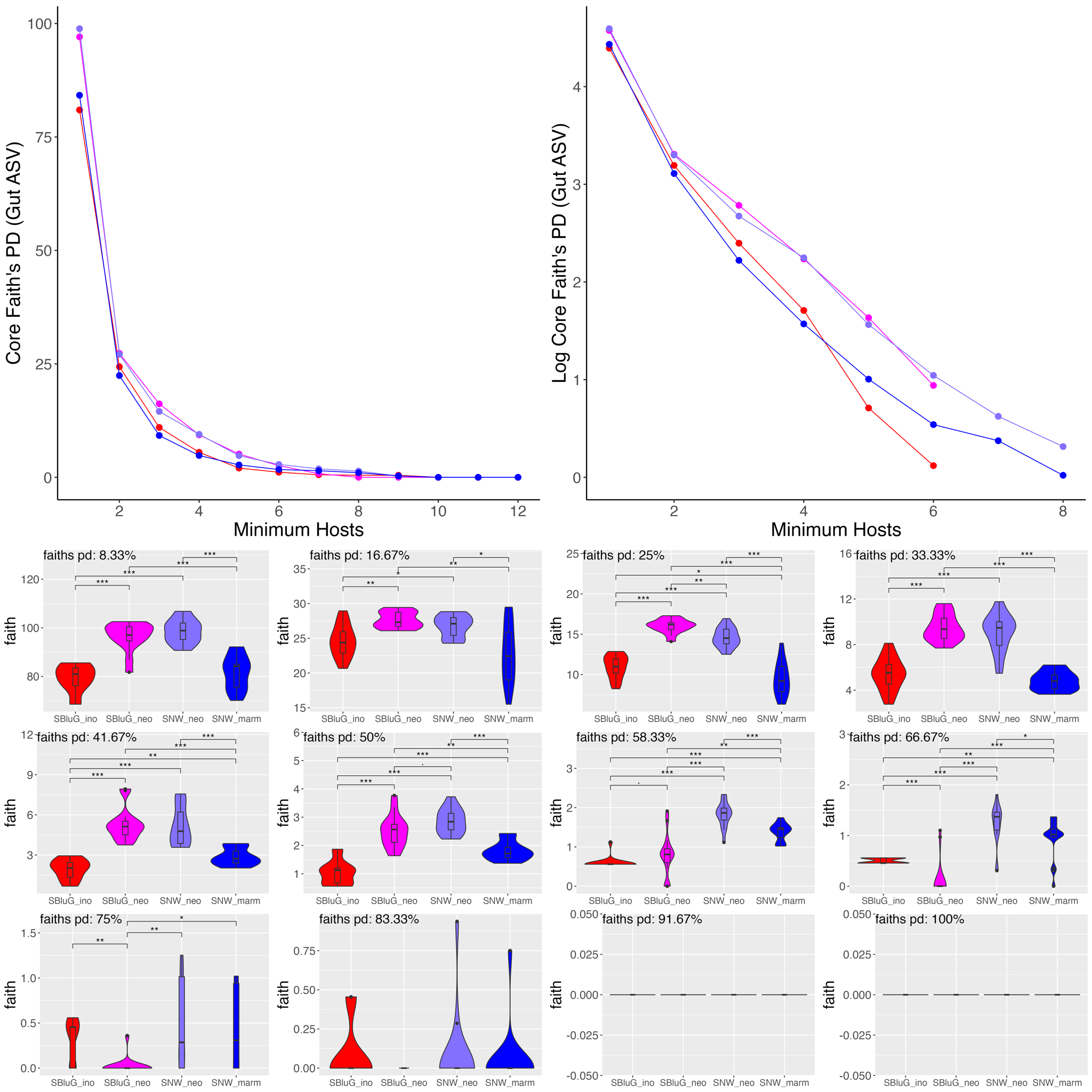

**Figure 2.3.** Faith’s PD of amplicon sequence variants for core gut microbiota as a function of the core threshold (i.e., the number of hosts an ASV must be present on in order to be classified as part of the core). In all panels, core microbiota Faith’s PD estimates were generated using 15 subsamples of 12 individuals from each population and colors, statistical tests and symbols are as in Fig. 2.1. The top two panels show core Faith’s PD as a function of core threshold. The bottom twelve panels show violin plots for each particular core threshold.

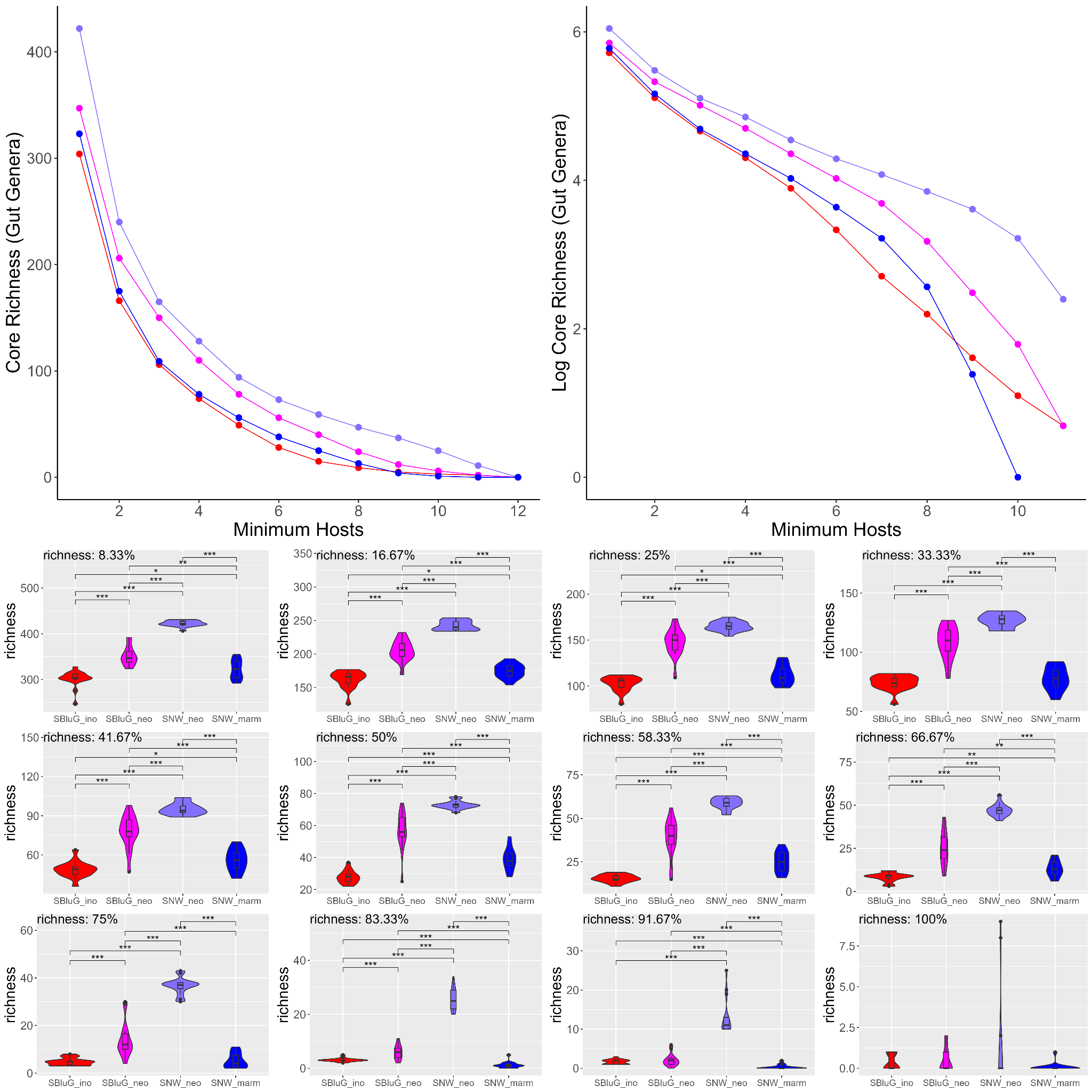

**Figure 2.4.** Richness of genera (genus count) for core gut microbiota as a function of the core threshold (i.e., the number of hosts a genus must be present on in order to be classified as part of the core). In all panels, core microbiota richness estimates were generated using 15 subsamples of 12 individuals from each population and colors, statistical tests and symbols are as in Fig. 2.1. The top two panels show core richness as a function of core threshold. The bottom twelve panels show violin plots for each particular core threshold.

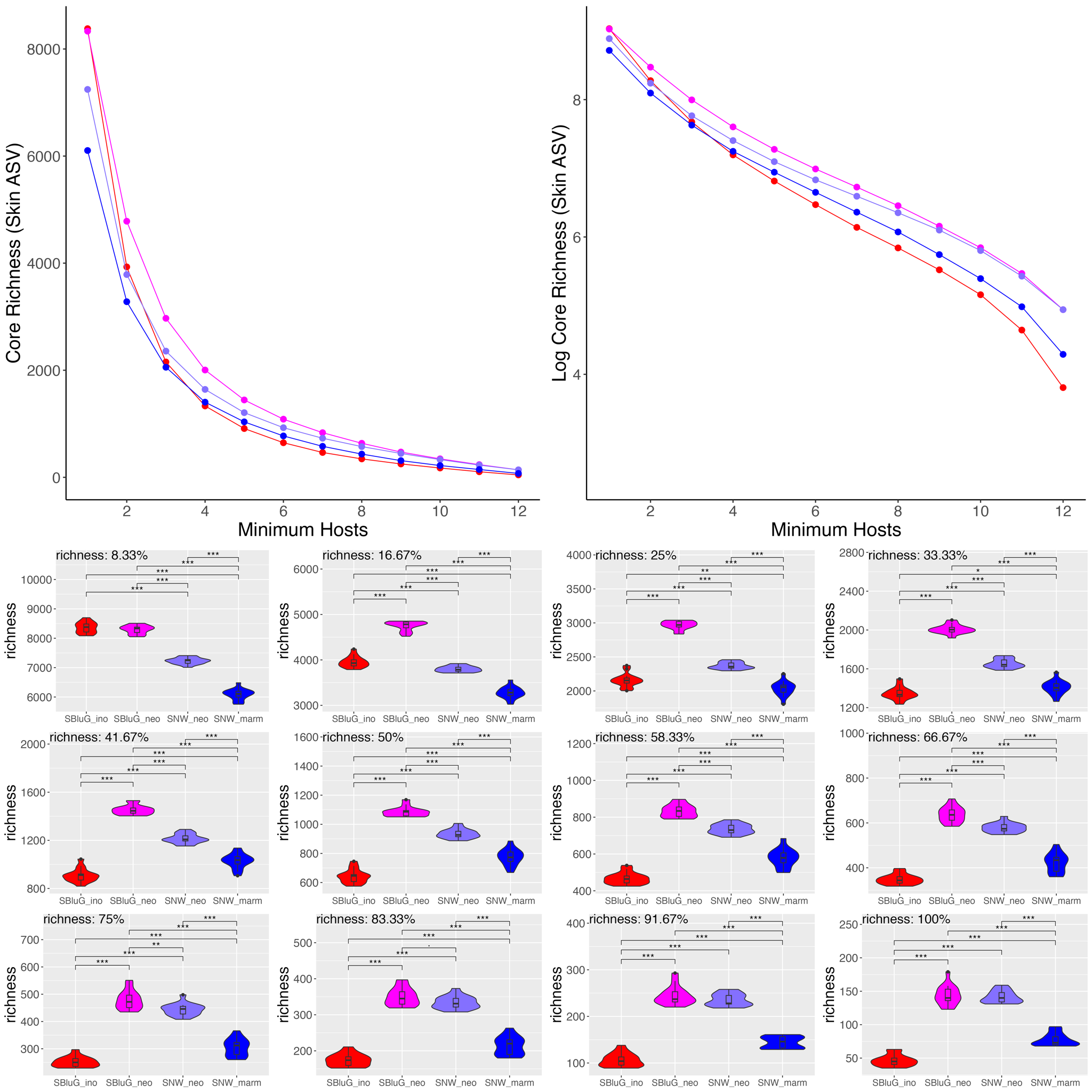

**Figure 2.5.** Richness of amplicon sequence variants (ASV count) for core skin microbiota as a function of the core threshold (i.e., the number of hosts an ASV must be present on in order to be classified as part of the core). In all panels, core microbiota richness estimates were generated using 15 subsamples of 12 individuals from each population and colors, statistical tests and symbols are as in Fig. 2.1. The top two panels show core richness as a function of core threshold. The bottom twelve panels show violin plots for each particular core threshold.

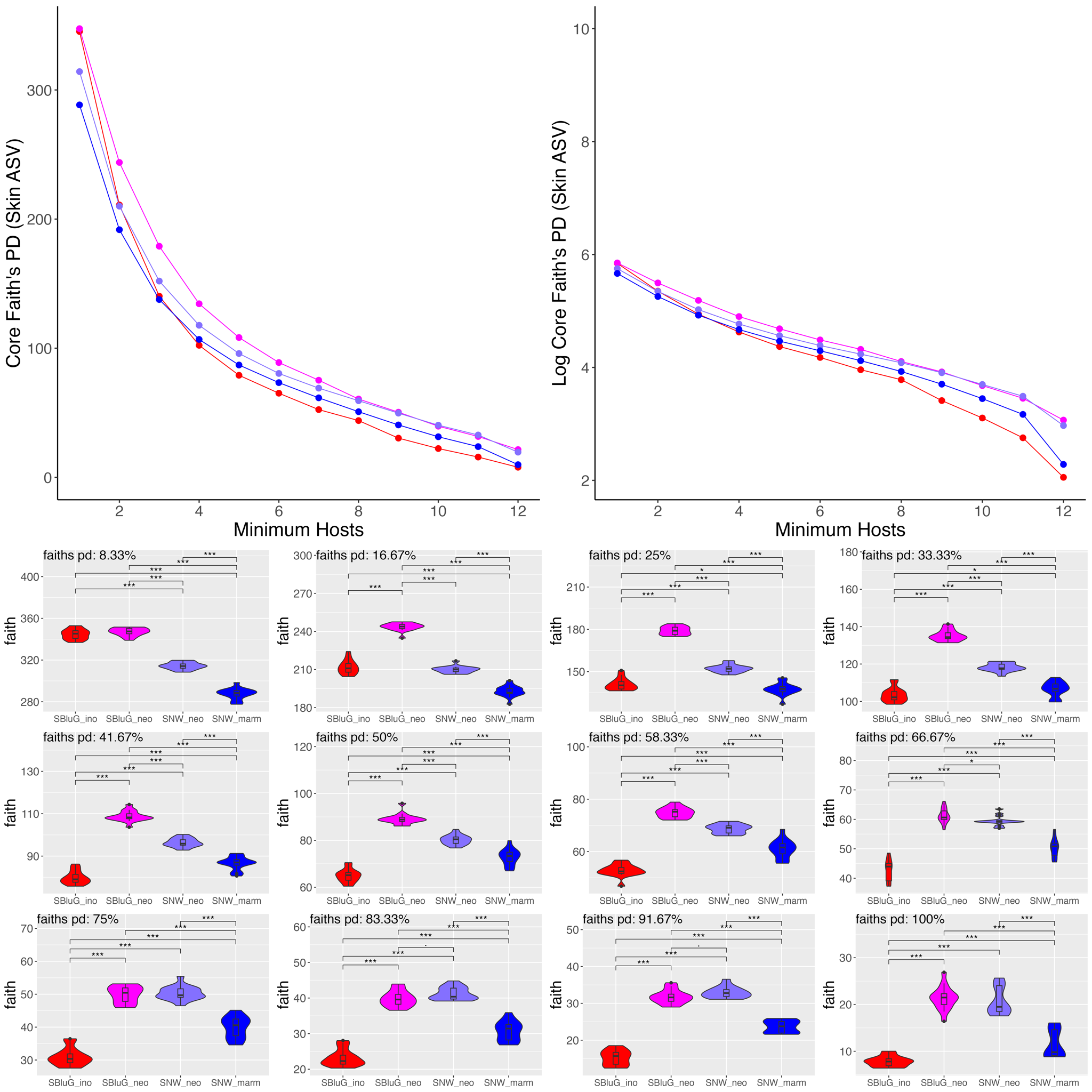

**Figure 2.6.** Faith’s PD of amplicon sequence variants for core skin microbiota as a function of the core threshold (i.e., the number of hosts an ASV must be present on in order to be classified as part of the core). In all panels, core microbiota Faith’s PD estimates were generated using 15 subsamples of 12 individuals from each population and colors, statistical tests and symbols are as in Fig. 2.1. The top two panels show core Faith’s PD as a function of core threshold. The bottom twelve panels show violin plots for each particular core threshold.

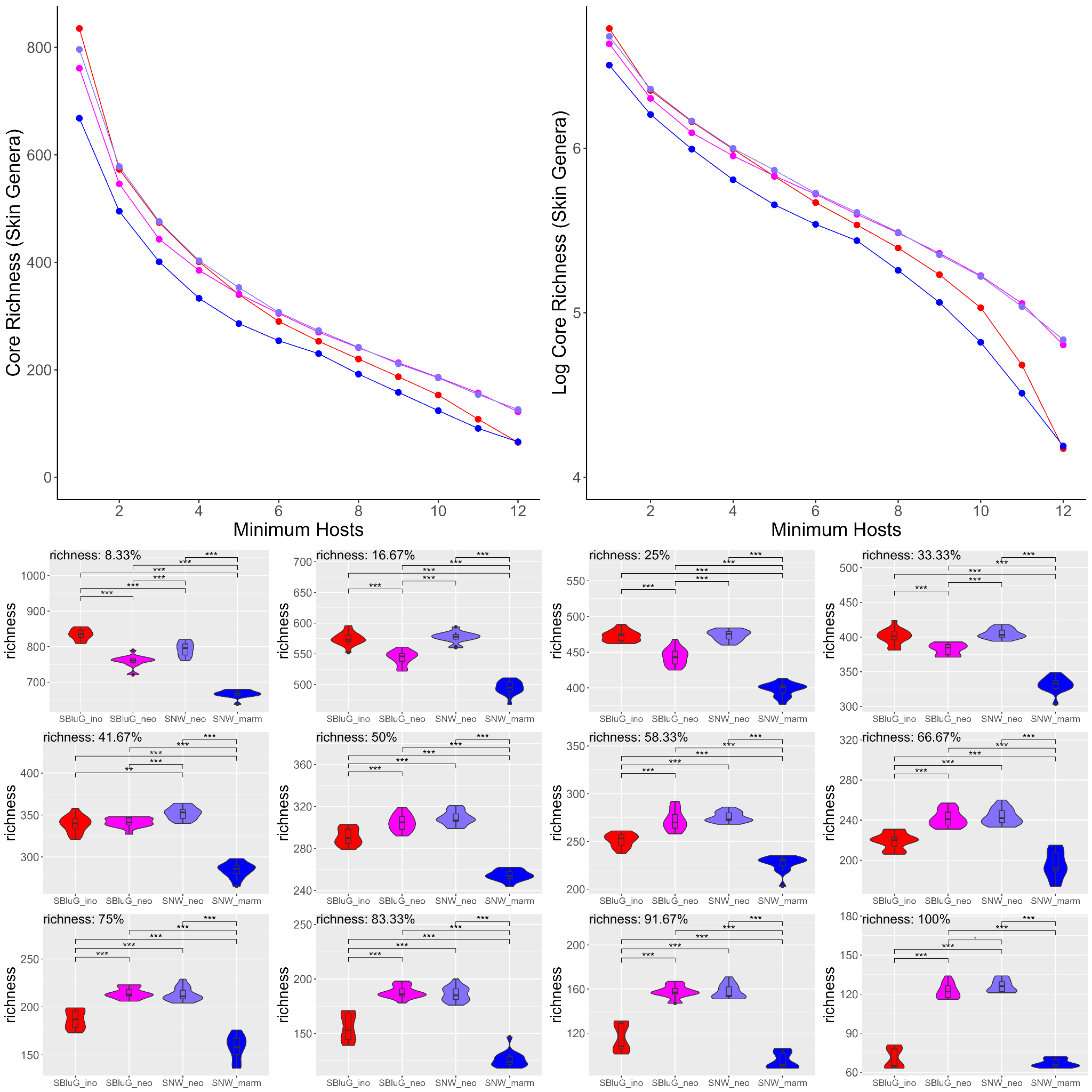

**Figure 2.7.** Richness of genera (genus count) for core skin microbiota as a function of the core threshold (i.e., the number of hosts a genus must be present on in order to be classified as part of the core). In all panels, core microbiota richness estimates were generated using 15 subsamples of 12 individuals from each population and colors, statistical tests and symbols are as in Fig. 2.1. The top two panels show core richness as a function of core threshold. The bottom twelve panels show violin plots for each particular core threshold.

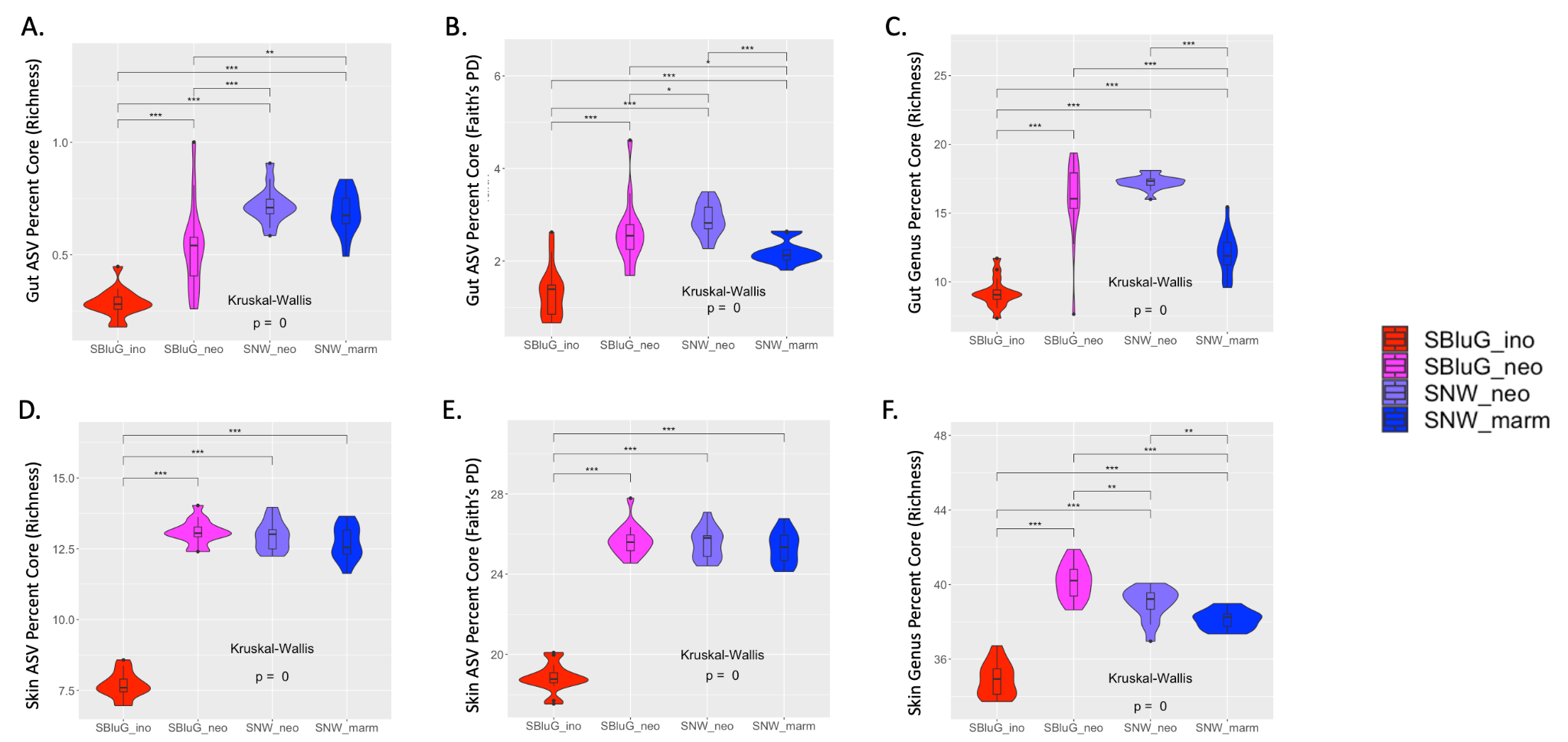

**Figure 2.8.** Comparison of the percentage of the microbiota that is part of the core for gut (A-C) and skin (D-F) microbiota between populations of *Aspidoscelis inornatus* from SBluG (red), *A. neomexicanus* from SBluG (magenta), *A. neomexicanus* from SNW (purple), and *A. marmoratus* from SNW (blue) as measured using (A, D) richness of amplicon sequence variants (ASV count), (B, E) Faith’s phylogenetic diversity (PD) of ASVs, and (C, F) richness of microbial genera (genera count). Significant differences in diversity between groups, as determined by a Kruskal-Wallis test followed by post hoc pairwise Wilcox tests using a Benjamini-Hochberg correction, are indicated as follows: p-value ≤ 0.001 (***), p-value ≤ 0.01 (**), p-value ≤ 0.05 (*), p-value ≤ 0.1 (.). In all panels, core microbiota diversity estimates were generated using 15 subsamples of 12 individuals from each population and core microbiota were determined at the 50% threshold (i.e., must be present on 6 hosts to be classified as part of the core).

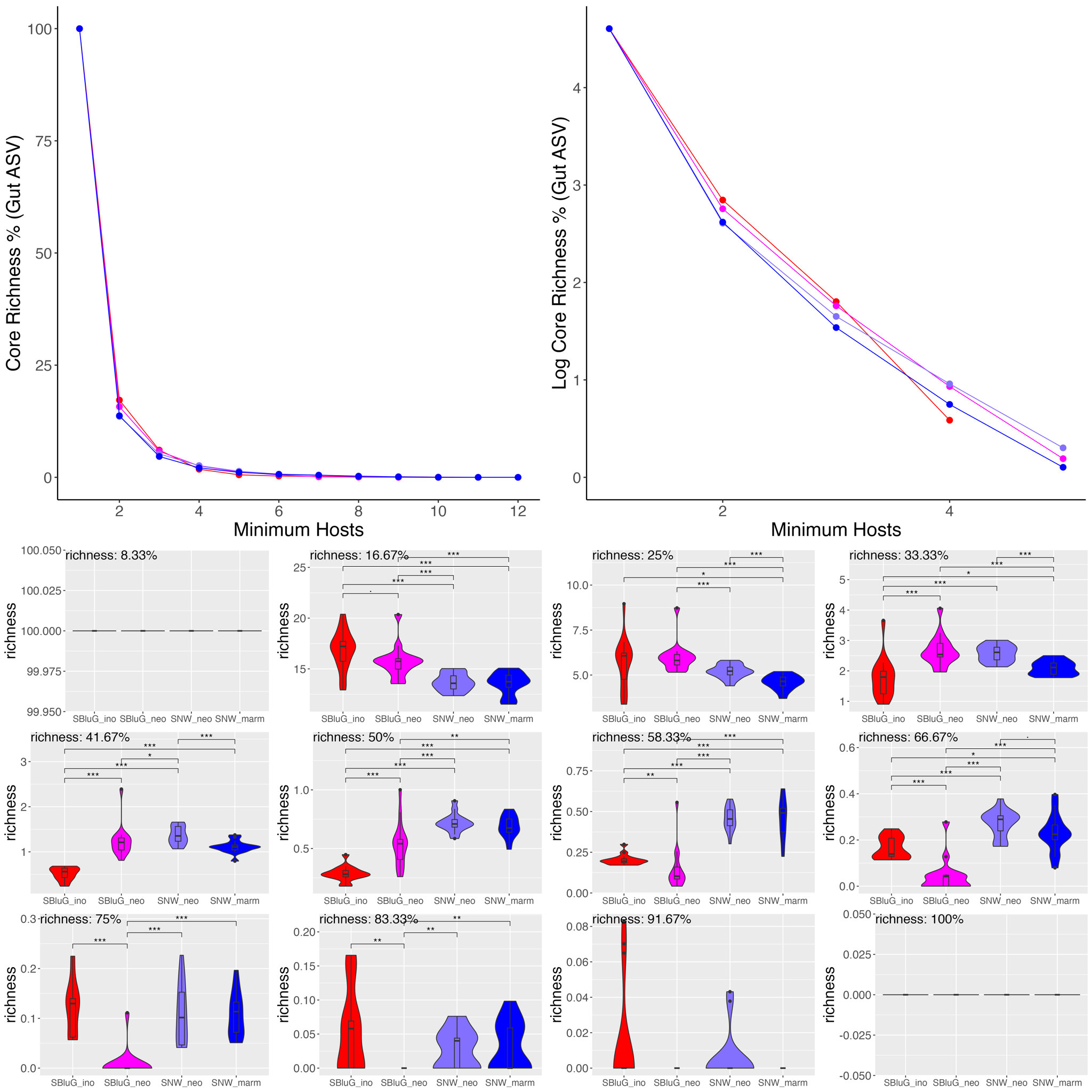

**Figure 2.9.** Percentage of amplicon sequence variant (ASV) richness that is part of the core gut microbiota as a function of the core threshold (i.e., the number of hosts an ASV must be present on in order to be classified as part of the core). In all panels, core microbiota richness estimates were generated using 15 subsamples of 12 individuals from each population and colors, statistical tests and symbols are as in Fig. 2.1. The top two panels show percentage of core richness as a function of core threshold. The bottom twelve panels show violin plots for each particular core threshold.

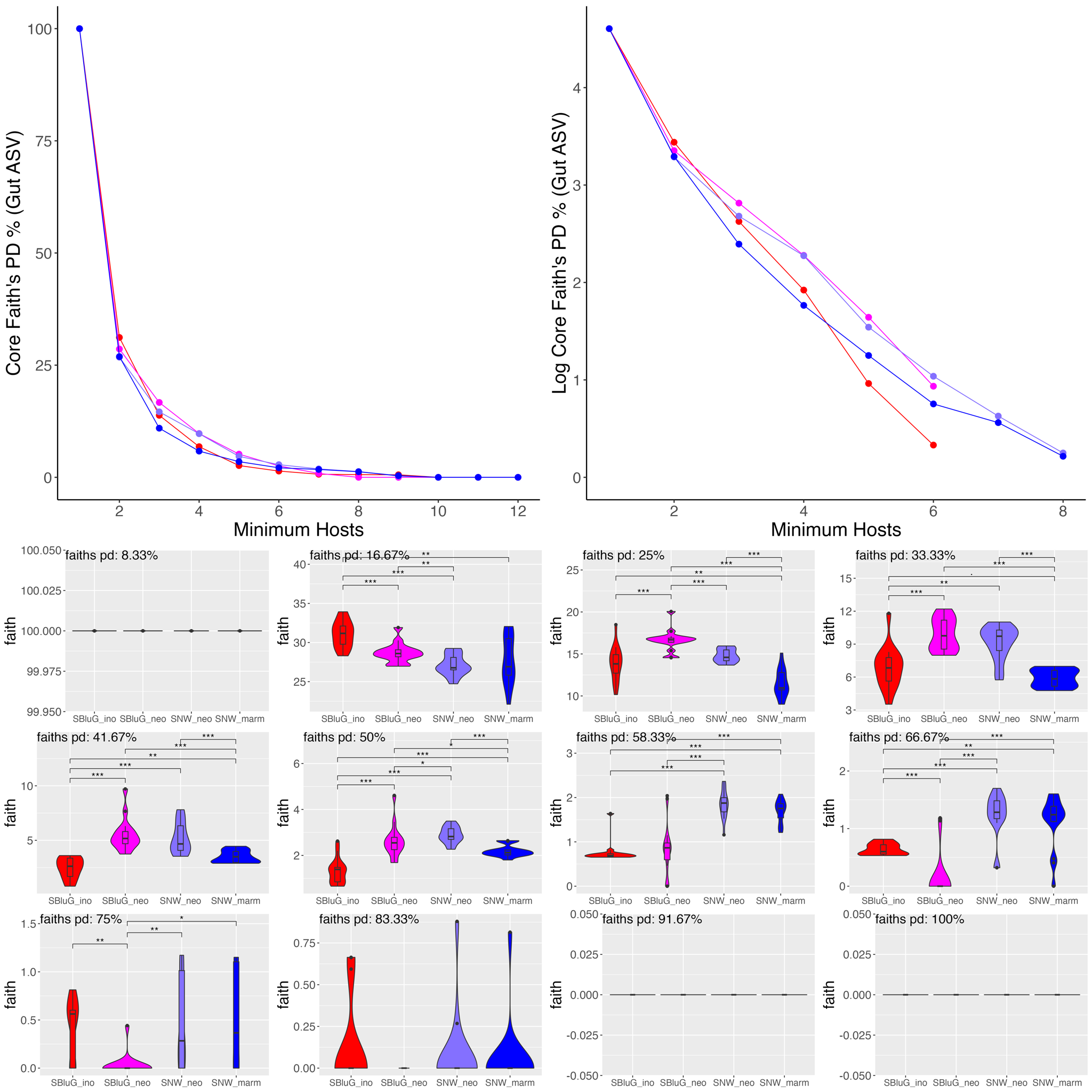

**Figure 2.10.** Percentage of Faith’s PD for amplicon sequence variants (ASV) that is part of the core gut microbiota as a function of the core threshold (i.e., the number of hosts an ASV must be present on in order to be classified as part of the core). In all panels, core microbiota Faith’s PD estimates were generated using 15 subsamples of 12 individuals from each population and colors, statistical tests and symbols are as in Fig. 2.1. The top two panels show percentage of core Faith’s PD as a function of core threshold. The bottom twelve panels show violin plots for each particular core threshold.

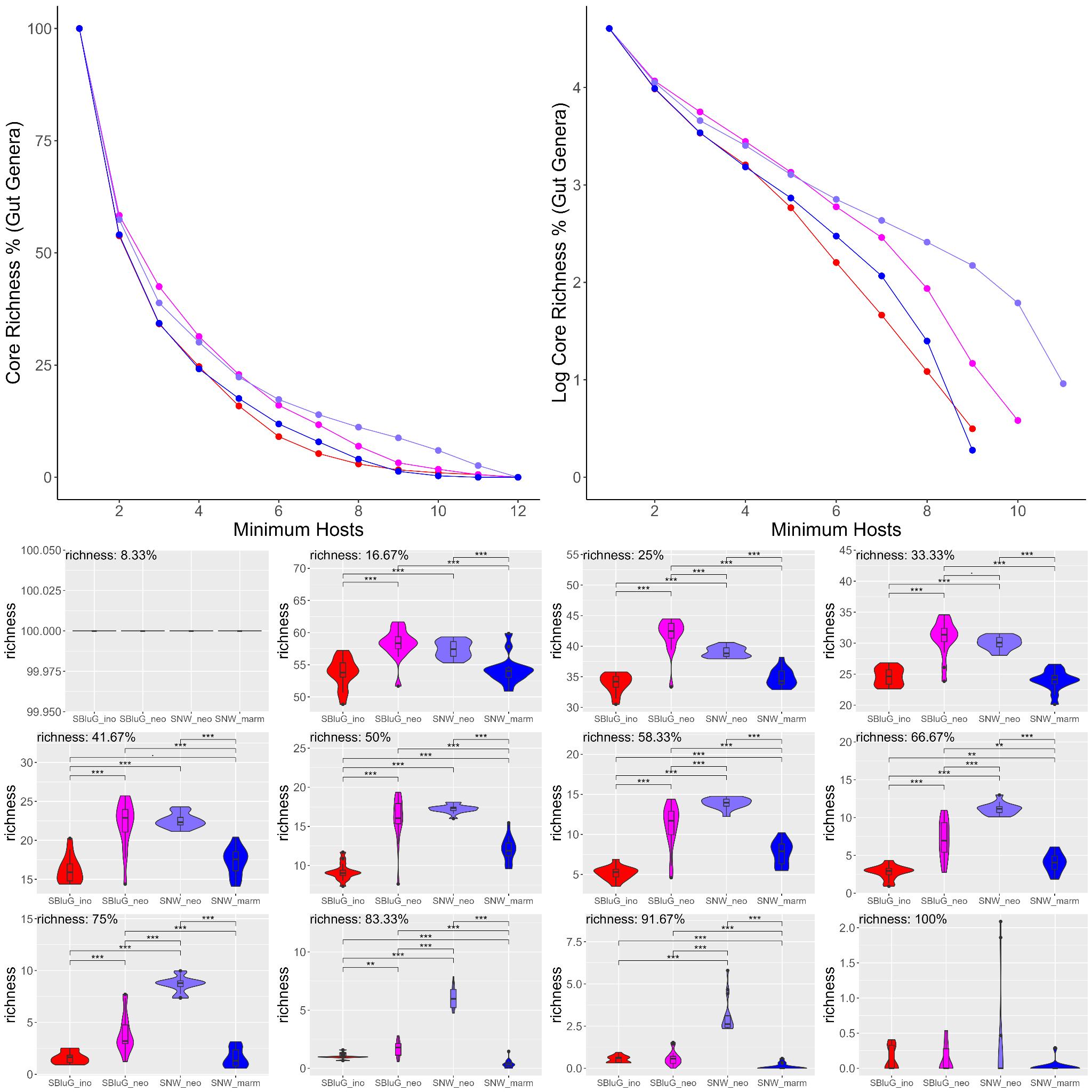

**Figure 2.11.** Percentage of genus richness that is part of the core gut microbiota as a function of the core threshold (i.e., the number of hosts a genus must be present on in order to be classified as part of the core). In all panels, core microbiota richness estimates were generated using 15 subsamples of 12 individuals from each population and colors, statistical tests and symbols are as in Fig. 2.1. The top two panels show percentage of core richness as a function of core threshold. The bottom twelve panels show violin plots for each particular core threshold.

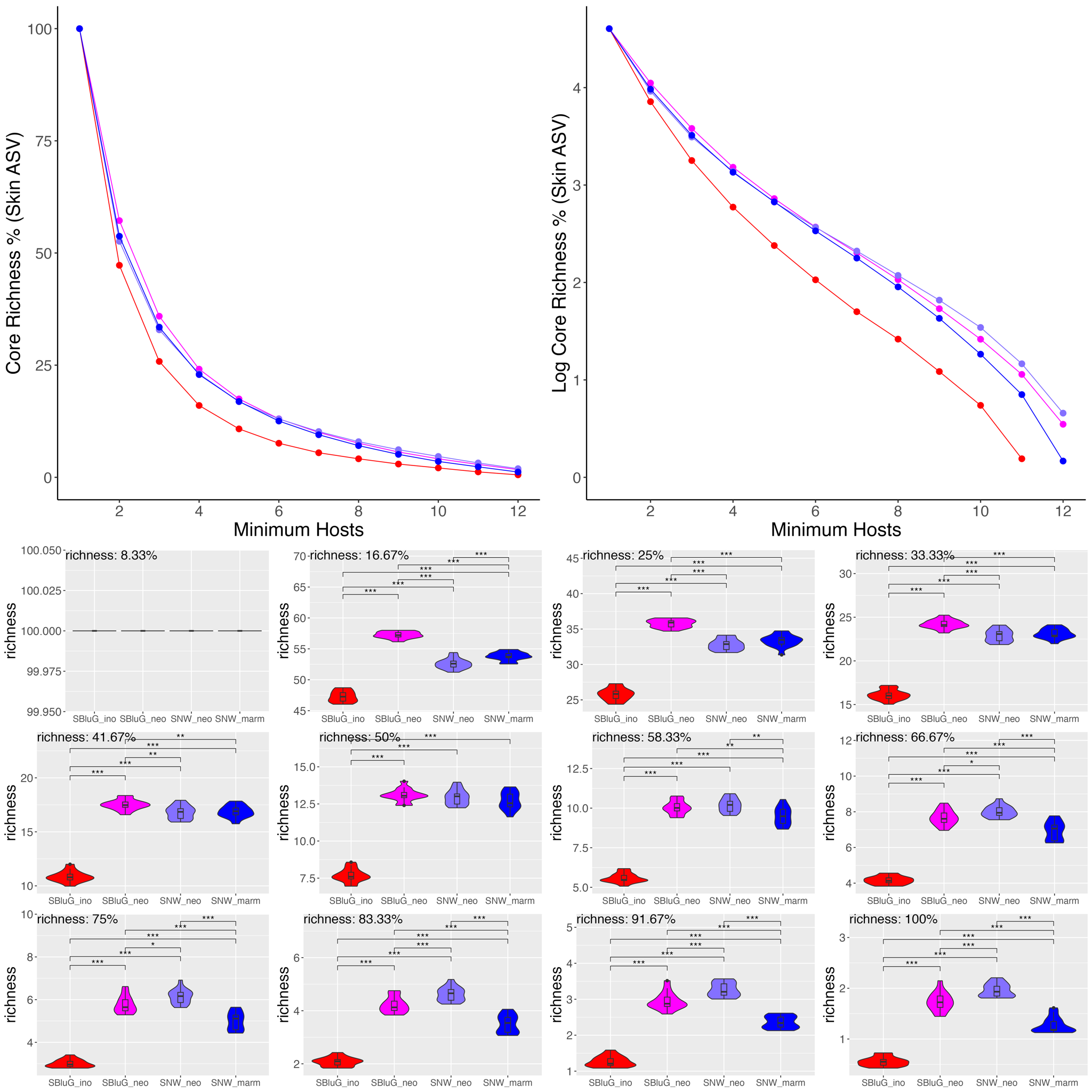

**Figure 2.12.** Percentage of amplicon sequence variant (ASV) richness that is part of the core skin microbiota as a function of the core threshold (i.e., the number of hosts an ASV must be present on in order to be classified as part of the core). In all panels, core microbiota diversity estimates were generated using 15 subsamples of 12 individuals from each population and colors, statistical tests and symbols are as in Fig. 2.1. The top two panels show percentage of core richness as a function of core threshold. The bottom twelve panels show violin plots for each particular core threshold.

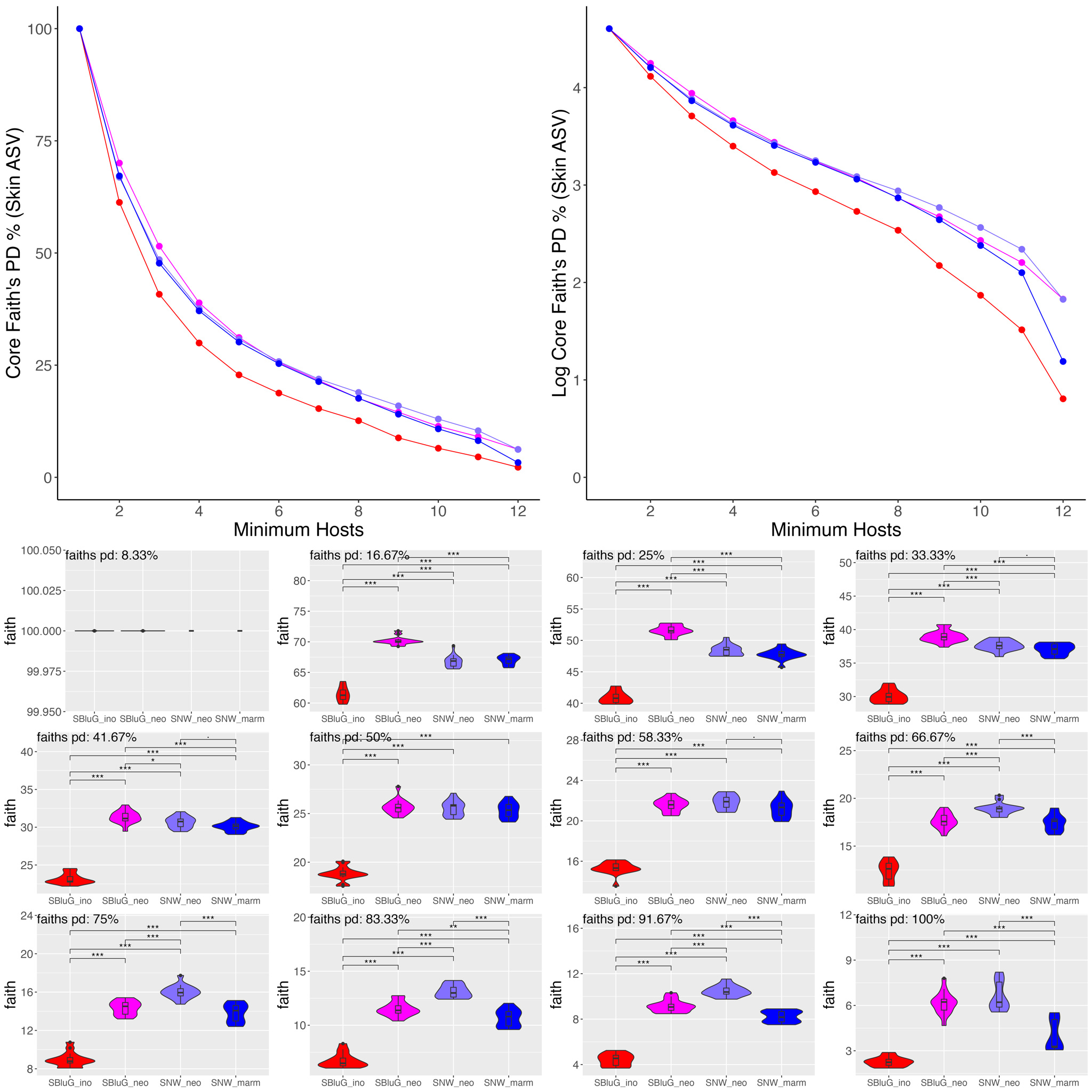

**Figure 2.13.** Percentage of Faith’s PD for amplicon sequence variants (ASV) that is part of the core skin microbiota as a function of the core threshold (i.e., the number of hosts an ASV must be present on in order to be classified as part of the core). In all panels, core microbiota Faith’s PD estimates were generated using 15 subsamples of 12 individuals from each population and colors, statistical tests and symbols are as in Fig. 2.1. The top two panels show percentage of core Faith’s PD as a function of core threshold. The bottom twelve panels show violin plots for each particular core threshold.

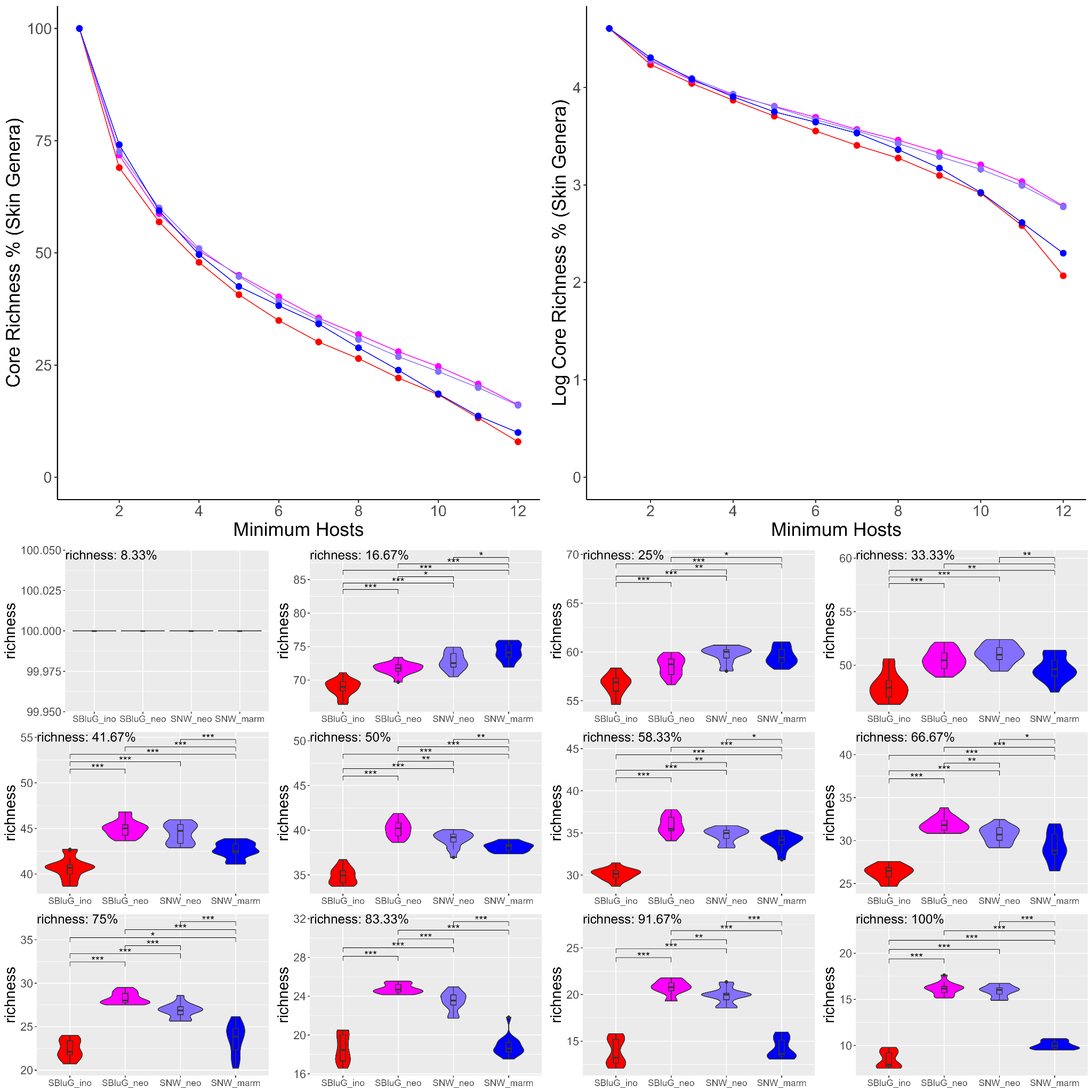

**Figure 2.14.** Percentage of genus richness that is part of the core skin microbiota as a function of the core threshold (i.e., the number of hosts a genus must be present on in order to be classified as part of the core). In all panels, core microbiota richness estimates were generated using 15 subsamples of 12 individuals from each population and colors, statistical tests and symbols are as in Fig. 2.1. The top two panels show percentage of core richness as a function of core threshold. The bottom twelve panels show violin plots for each particular core threshold.

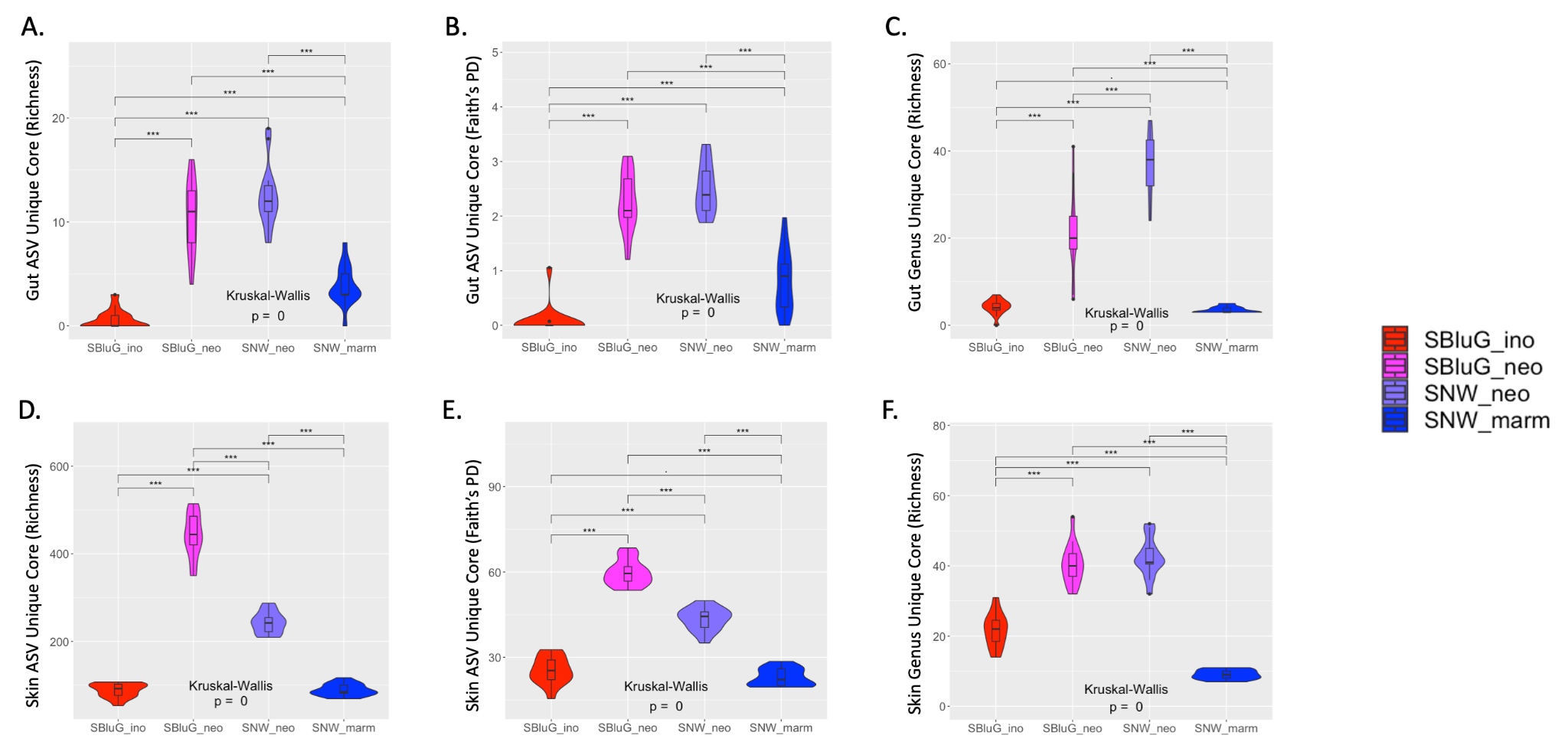

**Figure 2.15.** Comparison of the diversity of the unique component of the core gut (A-C) and skin (D-F) microbiota of populations of *Aspidoscelis inornatus* from SBluG (red), *A. neomexicanus* from SBluG (magenta), *A. neomexicanus* from SNW (purple), and *A. marmoratus* from SNW (blue) as measured using (A, D) richness of amplicon sequence variants (ASV count), (B, E) Faith’s phylogenetic diversity (PD) of ASVs, and (C, F) richness of microbial genera (genera count). Significant differences in diversity between groups, as determined by a Kruskal-Wallis test followed by post hoc pairwise Wilcox tests using a Benjamini-Hochberg correction, are indicated as follows: p-value ≤ 0.001 (***), p-value ≤ 0.01 (**), p-value ≤ 0.05 (*), p-value ≤ 0.1 (.). In all panels, core microbiota diversity estimates were generated using 15 subsamples of 12 individuals from each population and core microbiota were determined at the 50% threshold (i.e., must be present on 6 hosts to be classified as part of the core).

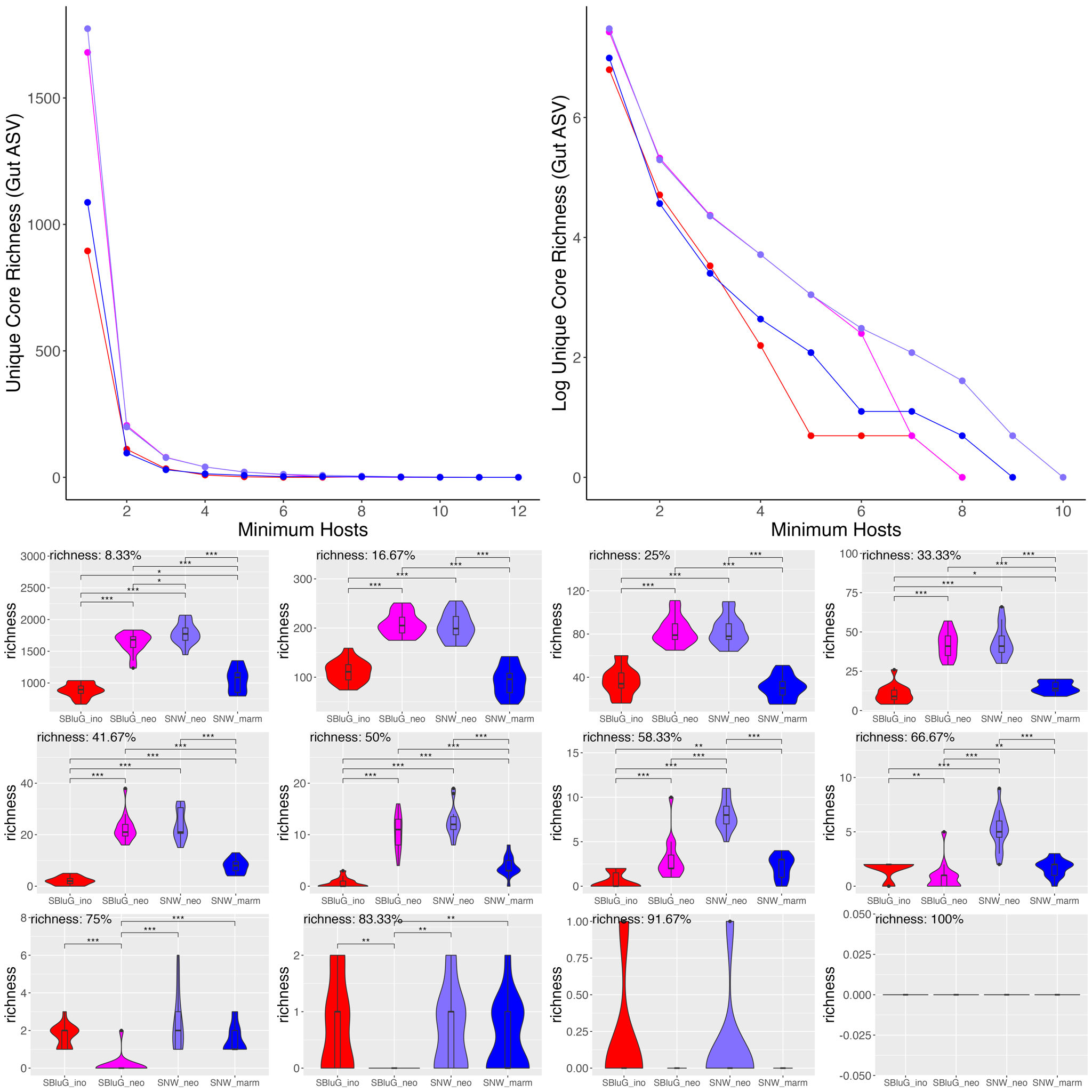

**Figure 2.16.** Richness of amplicon sequence variants (ASV count) for the core gut microbiota that is unique to each population as a function of the core threshold (i.e., the number of hosts an ASV must be present on in order to be classified as part of the core). In all panels, unique core microbiota richness estimates were generated using 15 subsamples of 12 individuals from each population and colors, statistical tests and symbols are as in Fig. 2.1. The top two panels show unique core richness as a function of core threshold. The bottom twelve panels show violin plots for each particular core threshold.

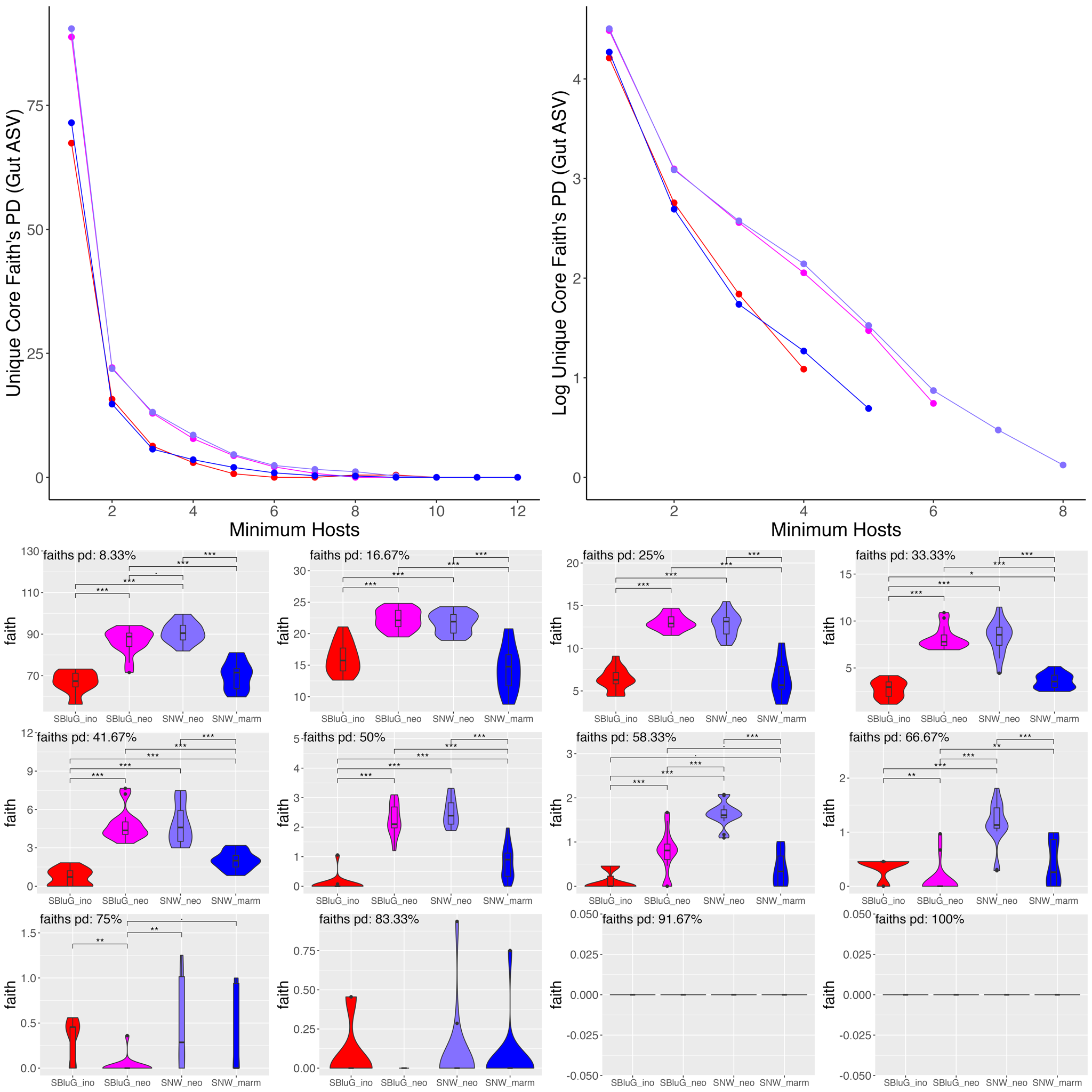

**Figure 2.17.** Faith’s PD of unique amplicon sequence variants for core gut microbiota as a function of the core threshold (i.e., the number of hosts an ASV must be present on in order to be classified as part of the core). In all panels, unique core microbiota Faith’s PD estimates were generated using 15 subsamples of 12 individuals from each population and colors, statistical tests and symbols are as in Fig. 2.1. The top two panels show unique core Faith’s PD as a function of core threshold. The bottom twelve panels show violin plots for each particular core threshold.

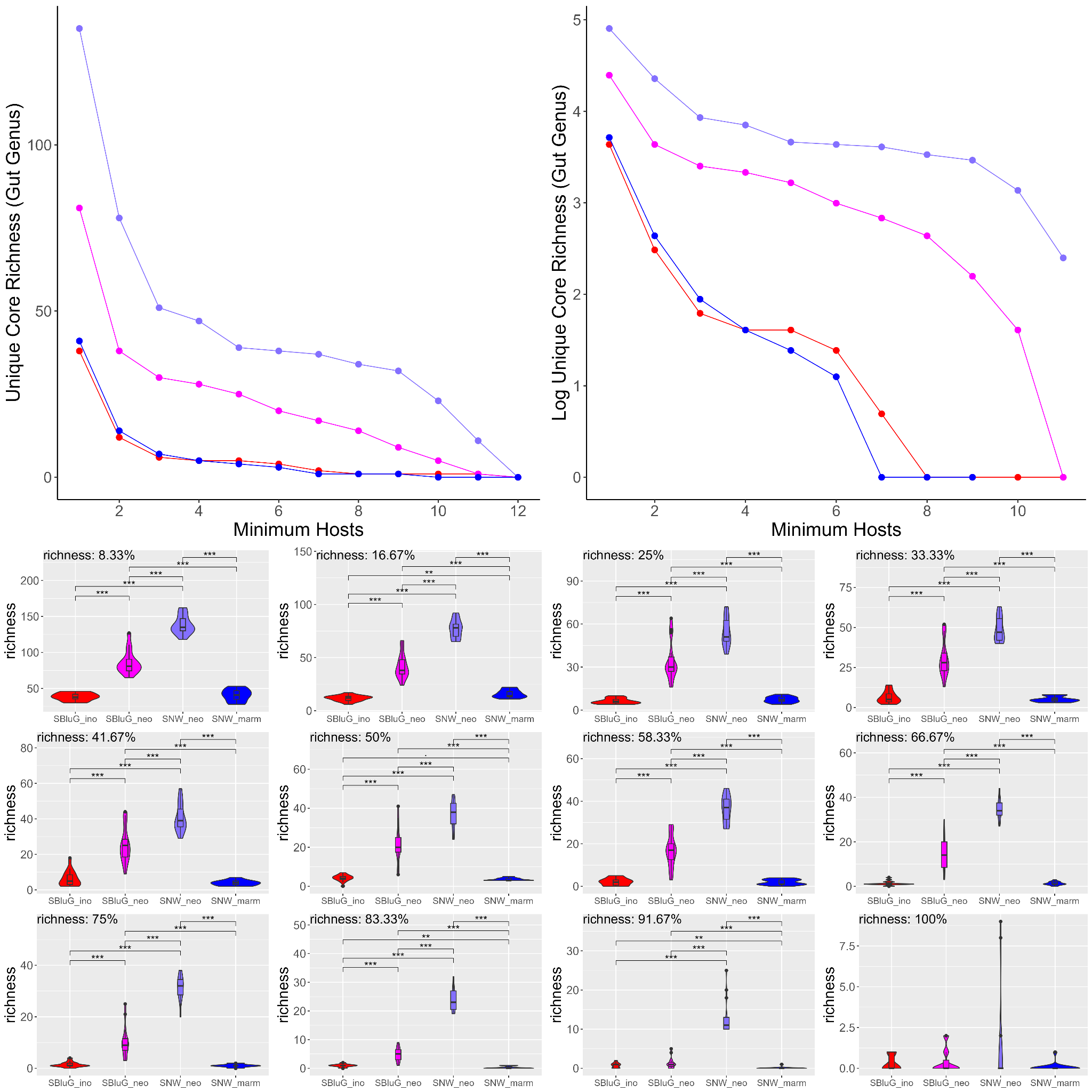

**Figure 2.18.** Richness of genera (genus count) for core gut microbiota that is unique to each population as a function of the core threshold (i.e., the number of hosts a genus must be present on in order to be classified as part of the core). In all panels, unique core microbiota richness estimates were generated using 15 subsamples of 12 individuals from each population and colors, statistical tests and symbols are as in Fig. 2.1. The top two panels show unique core richness as a function of core threshold. The bottom twelve panels show violin plots for each particular core threshold.

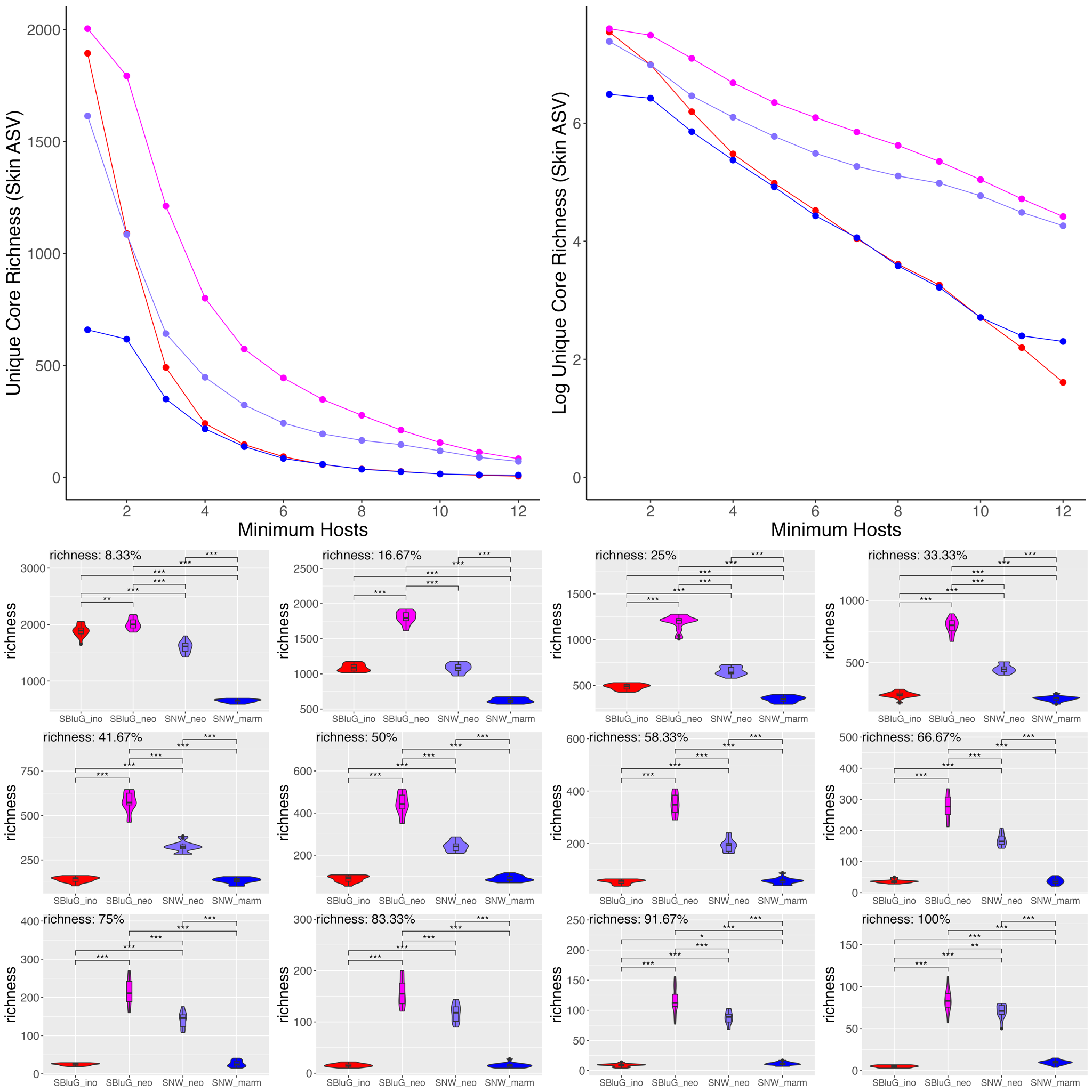

**Figure 2.19.** Richness of amplicon sequence variants (ASV count) for the core skin microbiota that is unique to each population as a function of the core threshold (i.e., the number of hosts an ASV must be present on in order to be classified as part of the core). In all panels, unique core microbiota richness estimates were generated using 15 subsamples of 12 individuals from each population and colors, statistical tests and symbols are as in Fig. 2.1. The top two panels show unique core richness as a function of core threshold. The bottom twelve panels show violin plots for each particular core threshold.

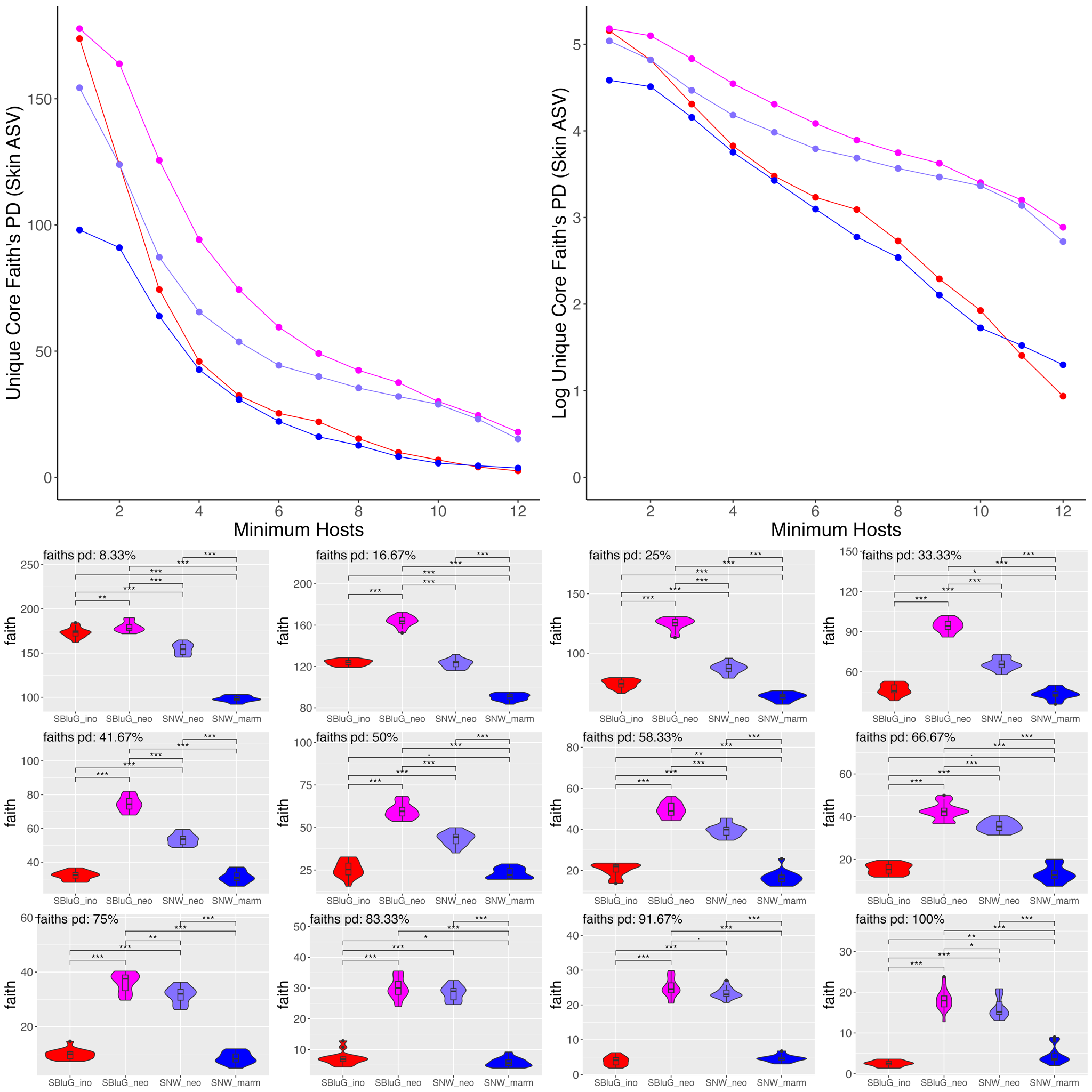

**Figure 2.20.** Faith’s PD of unique amplicon sequence variants for core skin microbiota as a function of the core threshold (i.e., the number of hosts an ASV must be present on in order to be classified as part of the core). In all panels, unique core microbiota Faith’s PD estimates were generated using 15 subsamples of 12 individuals from each population and colors, statistical tests and symbols are as in Fig. 2.1. The top two panels show unique core Faith’s PD as a function of core threshold. The bottom twelve panels show violin plots for each particular core threshold.

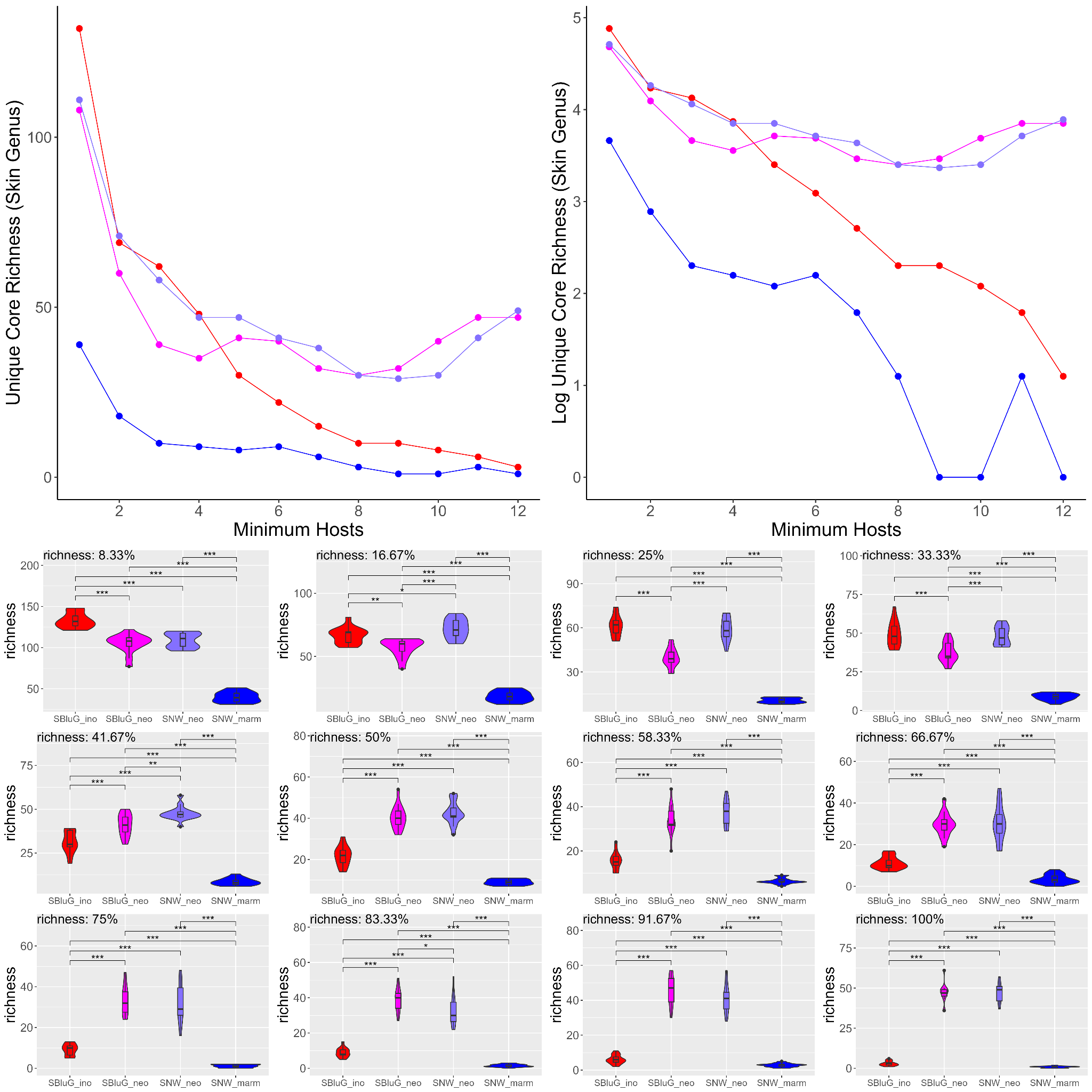

**Figure 2.21.** Richness of genera (genus count) for core skin microbiota that is unique to each population as a function of the core threshold (i.e., the number of hosts a genus must be present on in order to be classified as part of the core). In all panels, unique core microbiota richness estimates were generated using 15 subsamples of 12 individuals from each population and colors, statistical tests and symbols are as in Fig. 2.1. The top two panels show unique core richness as a function of core threshold. The bottom twelve panels show violin plots for each particular core threshold.

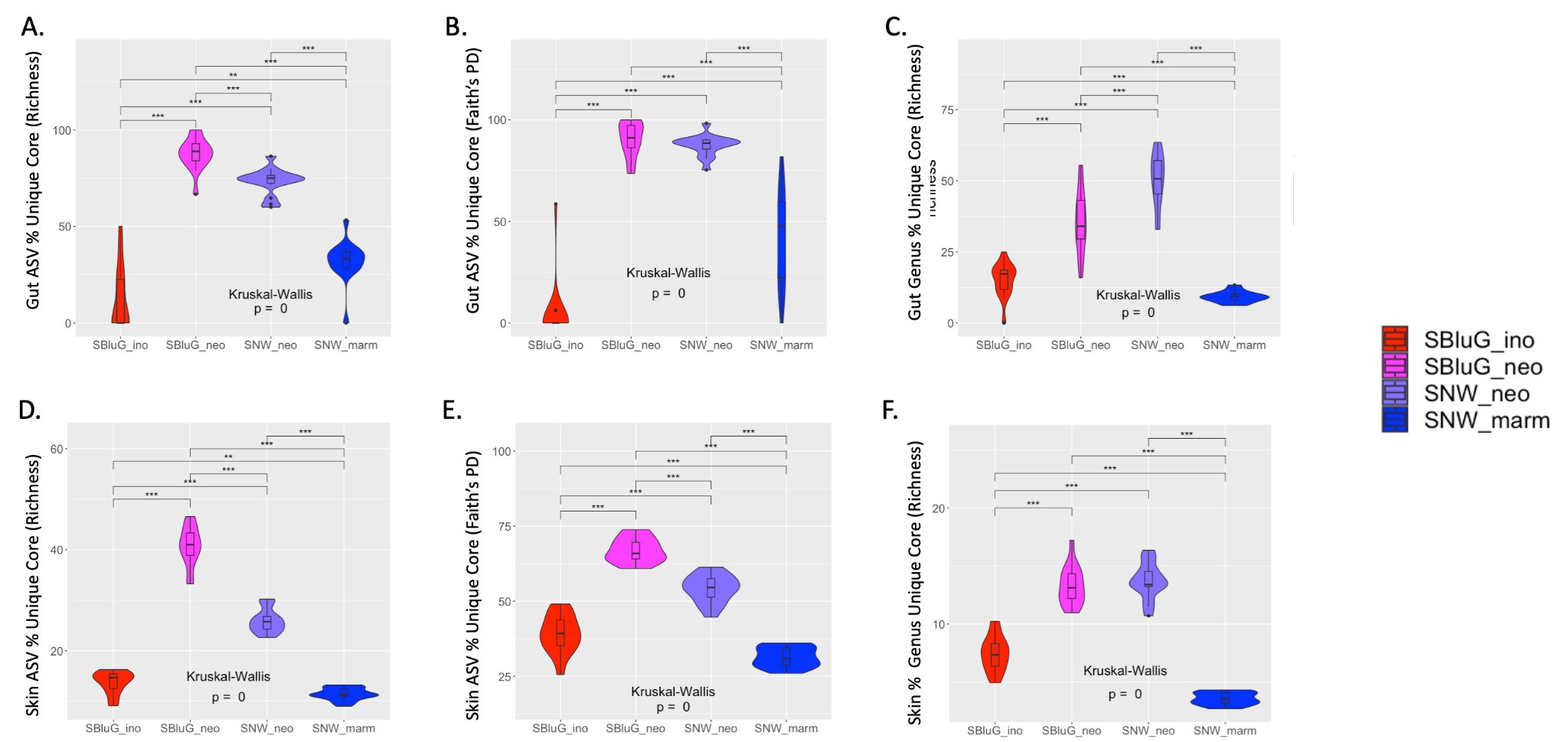

**Figure 2.22.** Comparison of the percentage of the core microbiota that is unique to a particular population for gut (A-C) and skin (D-F) microbiota between populations of *Aspidoscelis inornatus* from SBluG (red), *A. neomexicanus* from SBluG (magenta), *A. neomexicanus* from SNW (purple), and *A. marmoratus* from SNW (blue) as measured using (A, D) richness of amplicon sequence variants (ASV count), (B, E) Faith’s phylogenetic diversity (PD) of ASVs, and (C, F) richness of microbial genera (genera count). Significant differences in diversity between groups, as determined by a Kruskal-Wallis test followed by post hoc pairwise Wilcox tests using a Benjamini-Hochberg correction, are indicated as follows: p-value ≤ 0.001 (***), p-value ≤ 0.01 (**), p-value ≤ 0.05 (*), p-value ≤ 0.1 (.). In all panels, core microbiota diversity estimates were generated using 15 subsamples of 12 individuals from each population and core microbiota were determined at the 50% threshold (i.e., must be present on 6 hosts to be classified as part of the core).

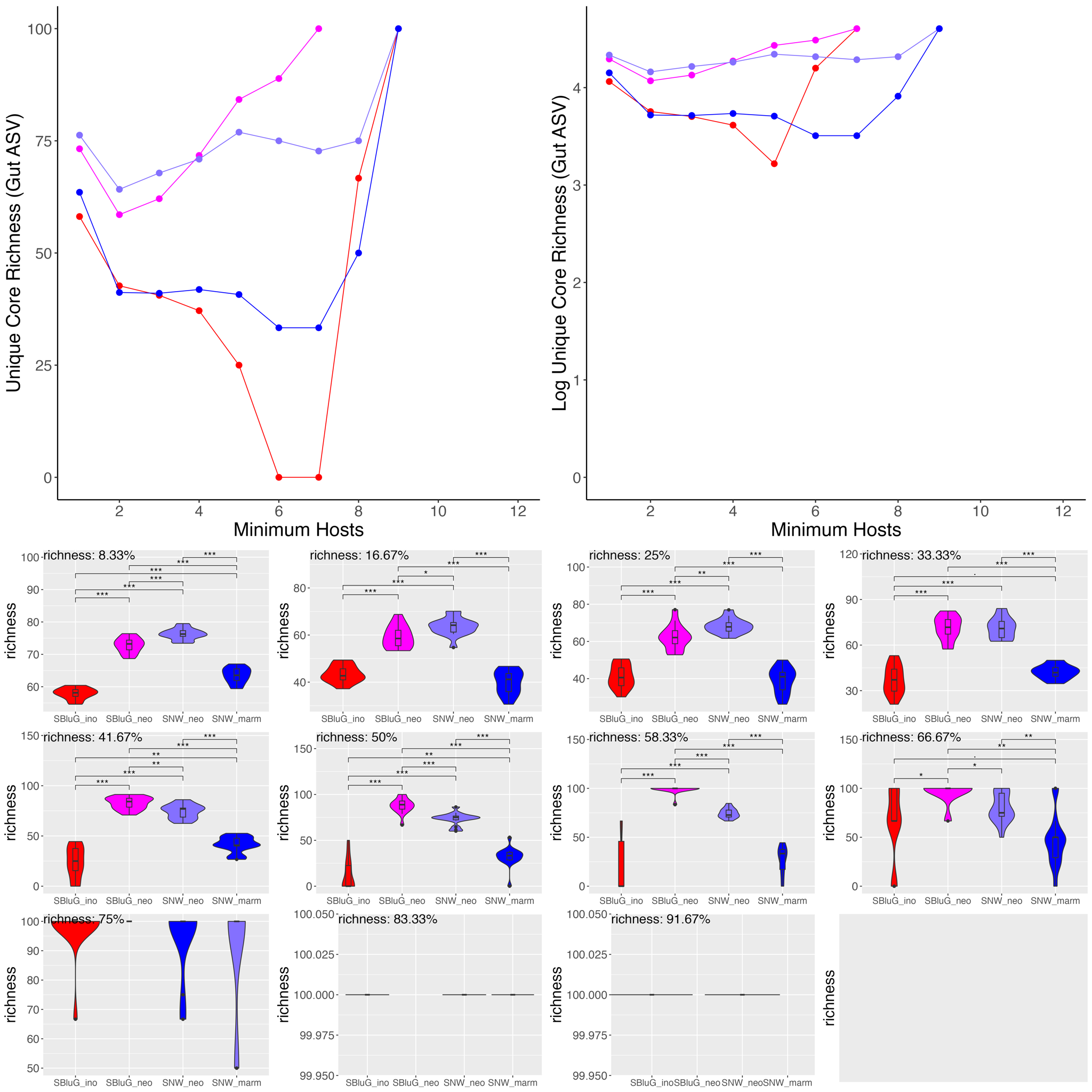

**Figure 2.23.** Percentage of gut core amplicon sequence variant (ASV) richness that is unique to each population as a function of the core threshold (i.e., the number of hosts an ASV must be present on in order to be classified as part of the core). In all panels, unique core microbiota richness estimates were generated using 15 subsamples of 12 individuals from each population and colors, statistical tests and symbols are as in Fig. 2.1. The top two panels show percentage of core richness that is unique as a function of core threshold. The bottom twelve panels show violin plots for each particular core threshold.

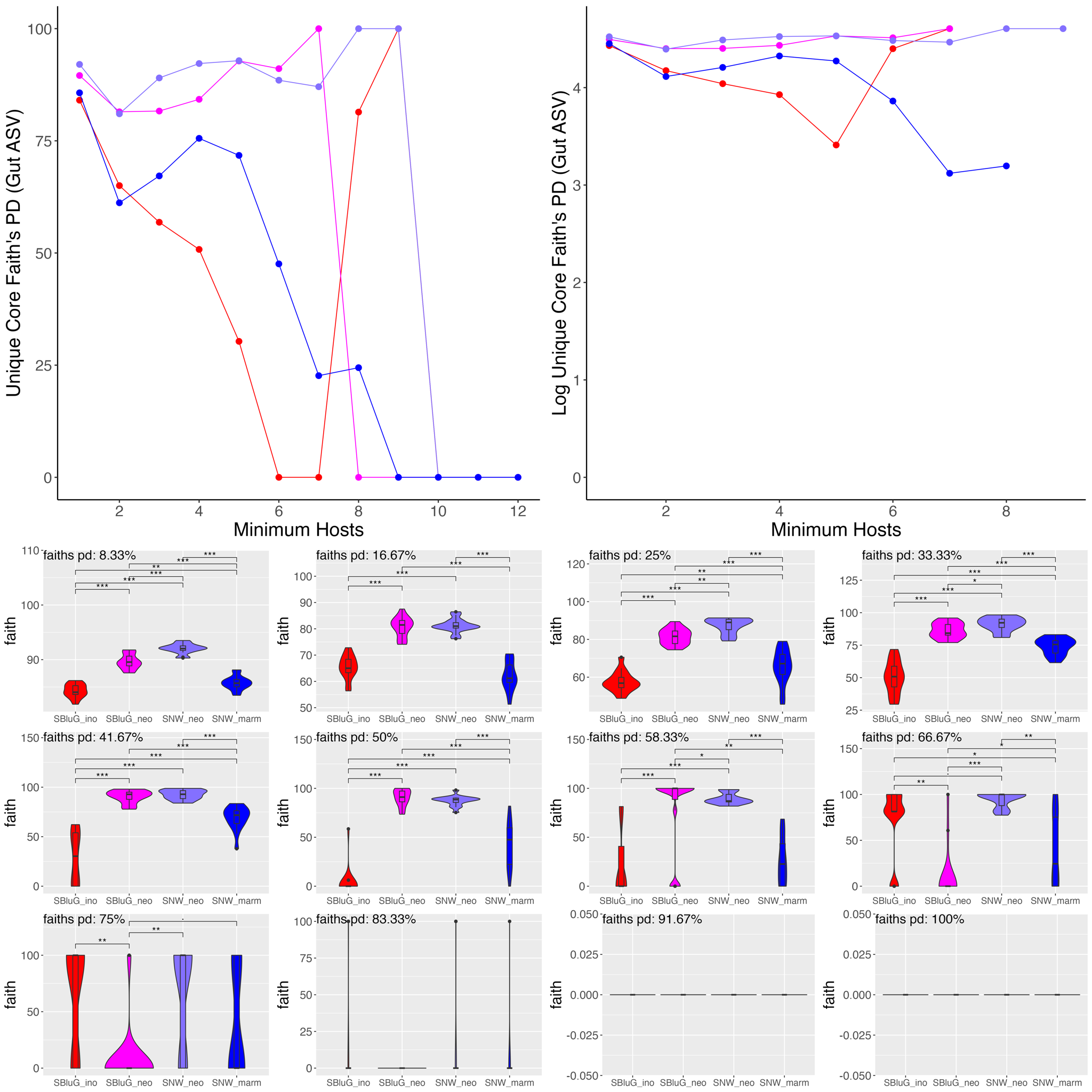

**Figure 2.24.** Percentage of gut core Faith’s PD for amplicon sequence variants (ASVs) that is unique to each population as a function of the core threshold (i.e., the number of hosts an ASV must be present on in order to be classified as part of the core). In all panels, unique core microbiota Faith’s PD estimates were generated using 15 subsamples of 12 individuals from each population and colors, statistical tests and symbols are as in Fig. 2.1. The top two panels show percentage of core Faith’s PD that is unique as a function of core threshold. The bottom twelve panels show violin plots for each particular core threshold.

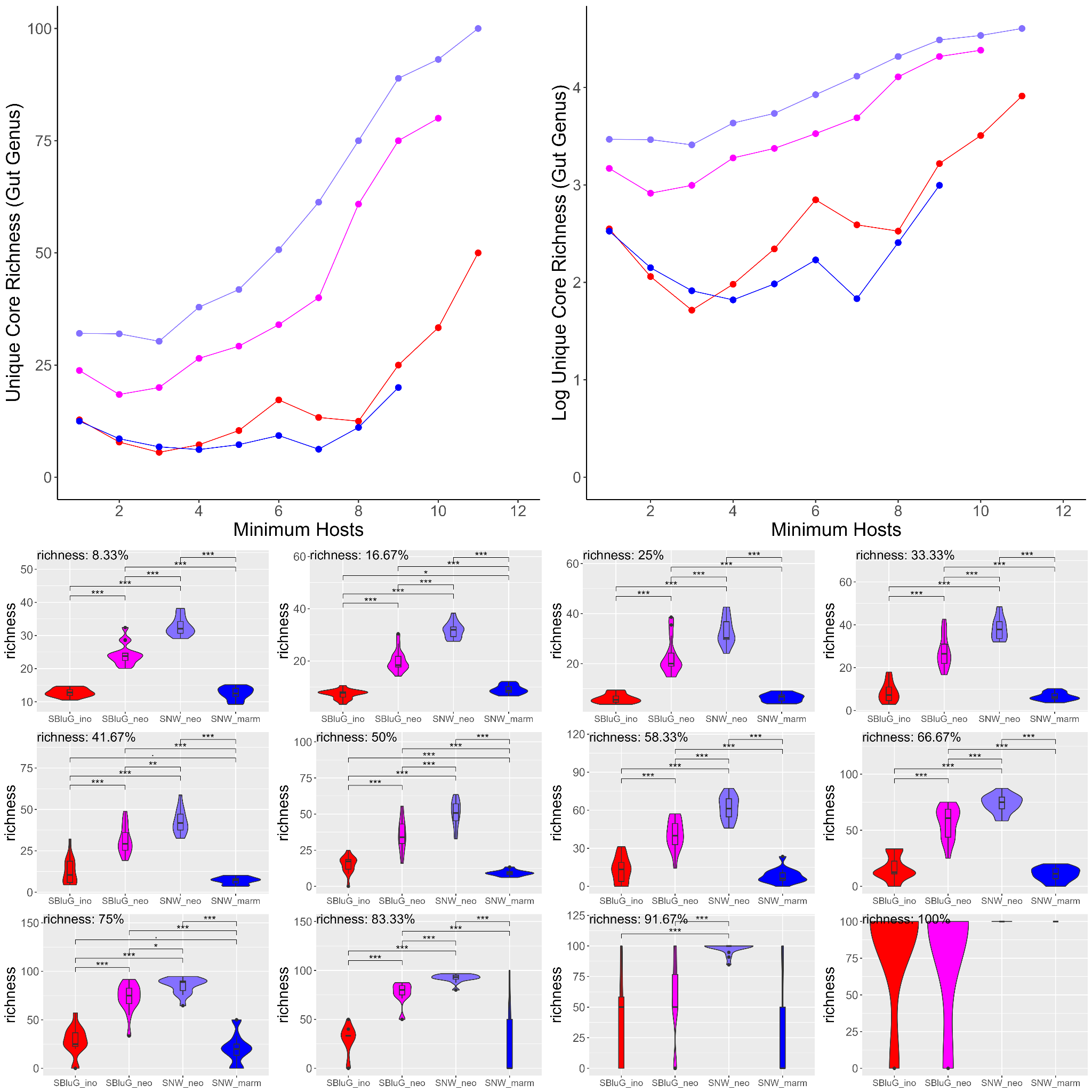

**Figure 2.25.** Percentage of gut genus richness that is unique to each population as a function of the core threshold (i.e., the number of hosts a genus must be present on in order to be classified as part of the core). In all panels, unique core microbiota richness estimates were generated using 15 subsamples of 12 individuals from each population and colors, statistical tests and symbols are as in Fig. 2.1. The top two panels show percentage of core richness that is unique as a function of core threshold. The bottom twelve panels show violin plots for each particular core threshold.

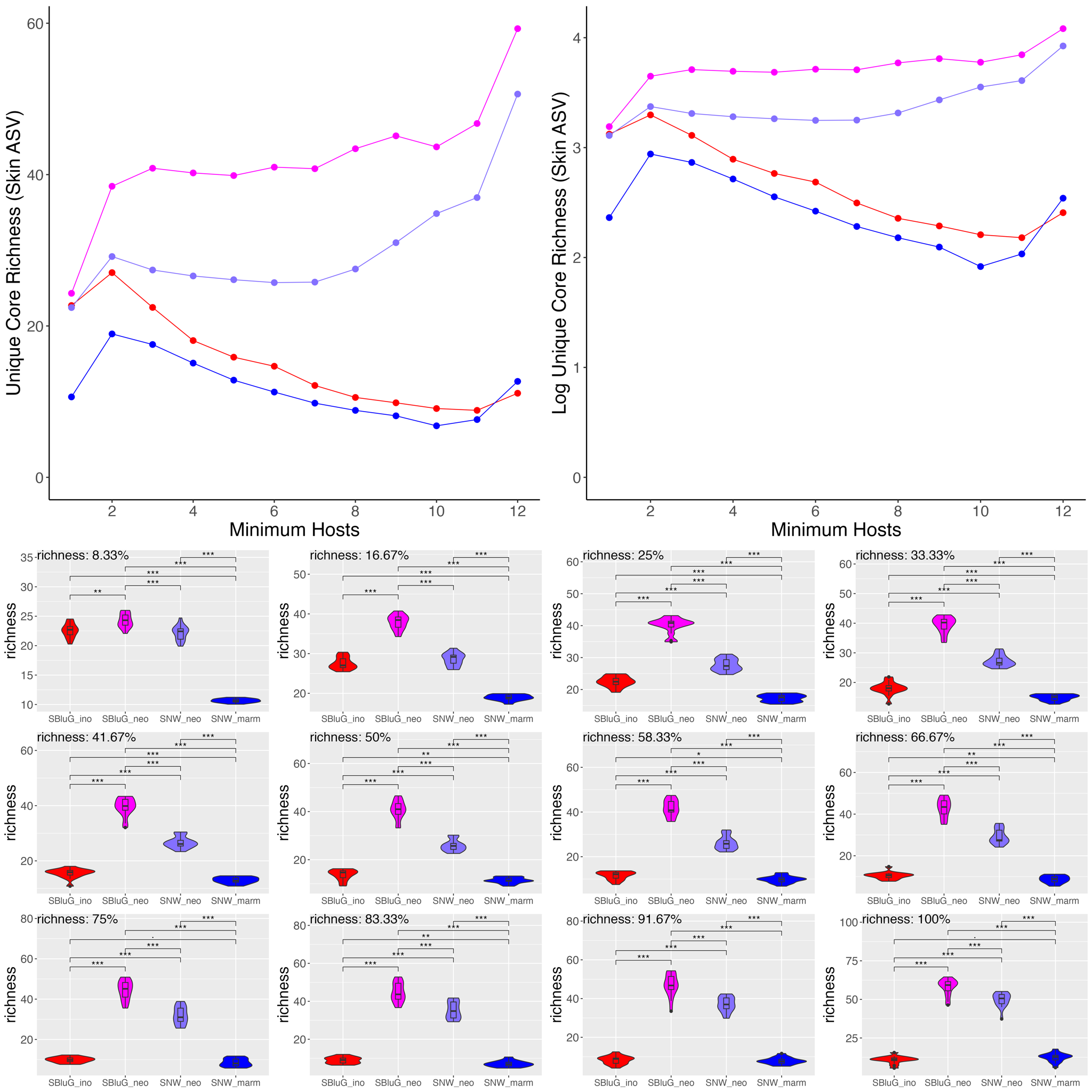

**Figure 2.26.** Percentage of skin core amplicon sequence variant (ASV) richness that is unique to each population as a function of the core threshold (i.e., the number of hosts an ASV must be present on in order to be classified as part of the core). In all panels, unique core microbiota richness estimates were generated using 15 subsamples of 12 individuals from each population and colors, statistical tests and symbols are as in Fig. 2.1. The top two panels show percentage of core richness that is unique as a function of core threshold. The bottom twelve panels show violin plots for each particular core threshold.

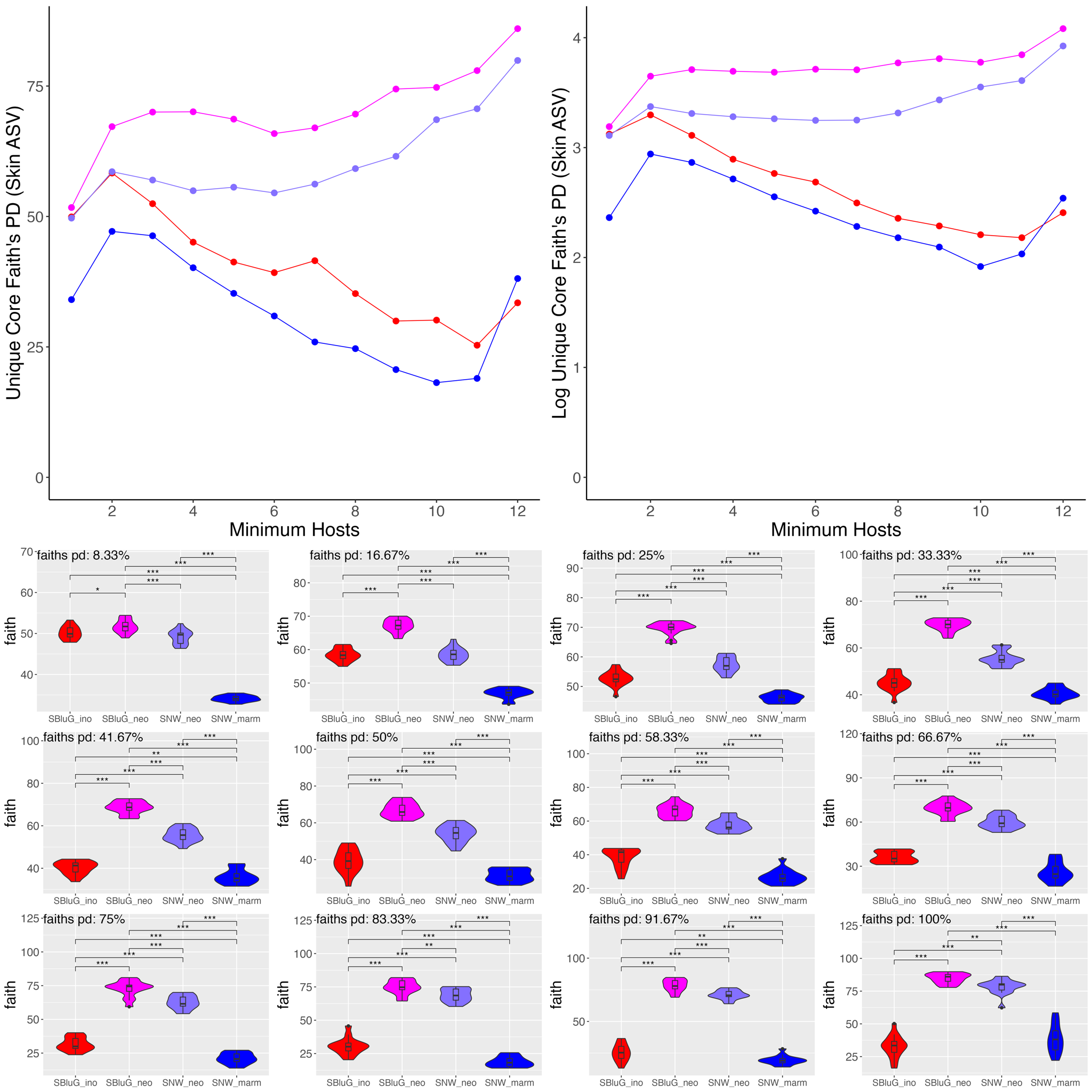

**Figure 2.27.** Percentage of skin core Faith’s PD for amplicon sequence variants (ASV) that is unique to each population as a function of the core threshold (i.e., the number of hosts an ASV must be present on in order to be classified as part of the core). In all panels, unique core microbiota Faith’s PD estimates were generated using 15 subsamples of 12 individuals from each population and colors, statistical tests and symbols are as in Fig. 2.1. The top two panels show percentage of core Faith’s PD that is unique as a function of core threshold. The bottom twelve panels show violin plots for each particular core threshold.

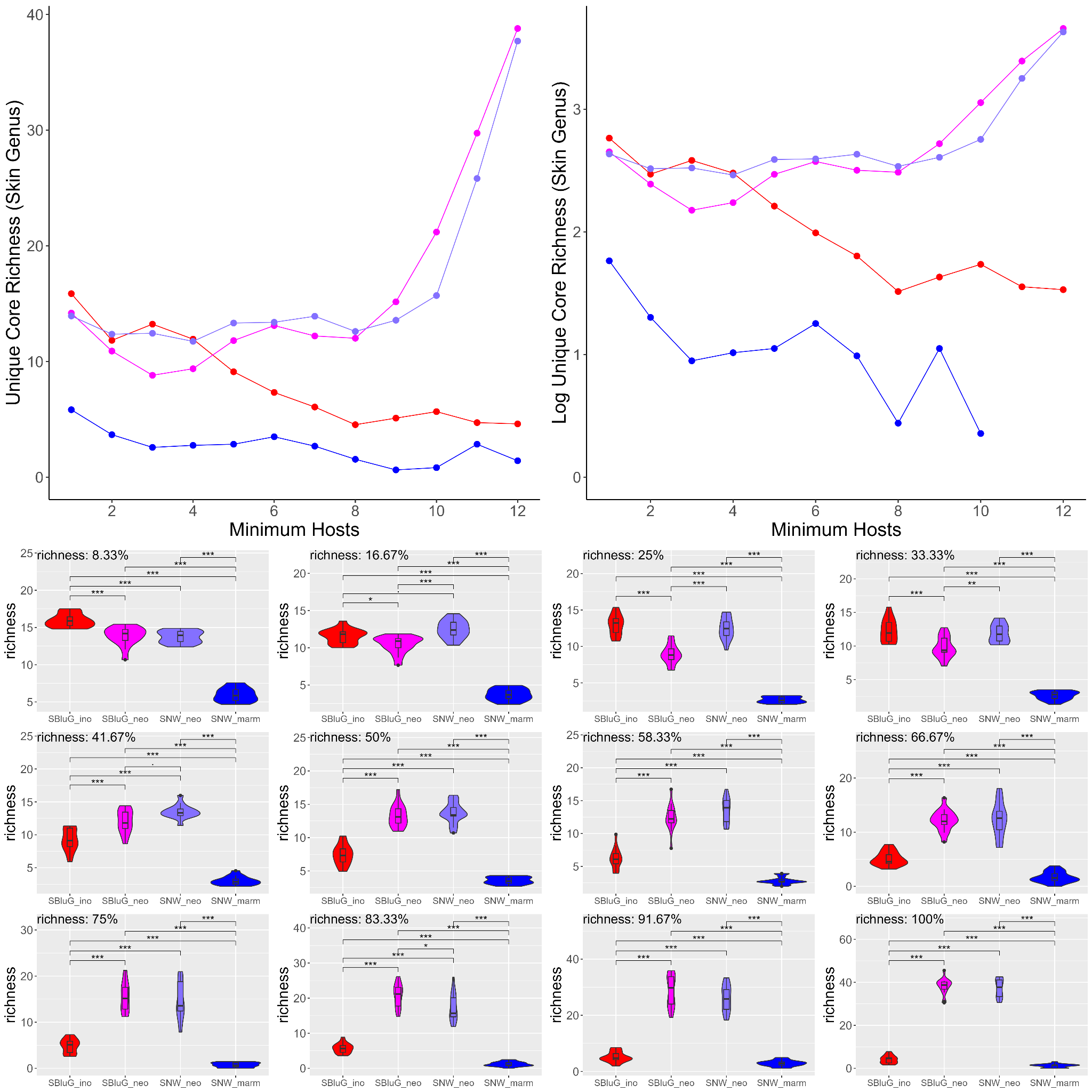

**Figure 2.28.** Percentage of skin genus richness that is unique to each population as a function of the core threshold (i.e., the number of hosts a genus must be present on in order to be classified as part of the core). In all panels, unique core microbiota richness estimates were generated using 15 subsamples of 12 individuals from each population and colors, statistical tests and symbols are as in Fig. 2.1. The top two panels show percentage of core richness that is unique as a function of core threshold. The bottom twelve panels show violin plots for each particular core threshold.

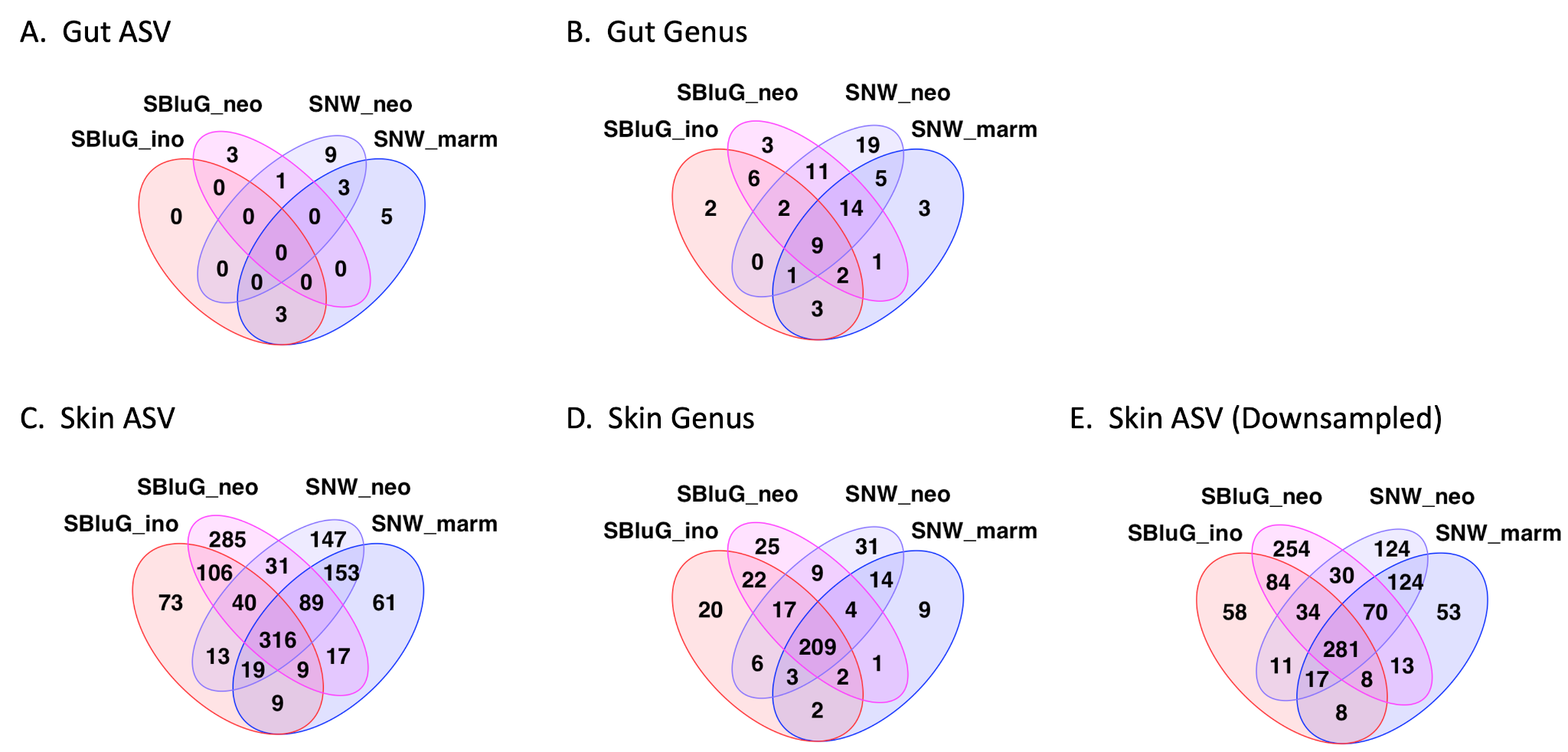

**Figure 2.29.** Venn diagrams showing the number of amplicon sequence variants (ASVs; A, C, E) and genera (B, D) shared among gut (A, B) and skin (C-E) microbiota of *Aspidoscelis inornatus* at SBluG (red), *A. marmoratus* at SNW (blue), *A. neomexicanus* at SBluG (magenta), and *A. neomexicanus* at SNW (purple). For all panels, a microbial ASV or genus is considered part of the core microbiota if it is found on at least 50% of individual lizards from a population. For panels A-D, microbiota were rarefied to the minimum reads present from the lizard microbiota with the lowest read count (gut: 11412 reads, skin: 116931). For panel E, we rarefied the skin microbiota to 11,412 reads, consistent with the depth available from our gut microbiota samples.
