## Additional file 6 for "Transgressive Hybrids as Hopeful Holobionts"

**Additional file 6: Dominant & Indicator Taxa**

**Dominant, indicator, and unique taxa**: The gut microbiota of all lizard populations are dominated by Actinobacteria, followed, in various orders for various lizard populations, by Protobacteria, Firmicutes and Bacteroidetes (see Fig. 5A and Table 6.1). However, the percent abundance of the four dominant orders varies considerably across lizard populations and species. Actinobacteria, for example, comprise on average 73.2% of the *Aspidoscelis marmoratus* gut microbiota and 53.2% of the *A. inornatus* gut microbiota, but only 39.3% of the *A. neomexicanus* gut microbiota. Inspection of individual lizard microbiota (see Fig. 5A) suggests that this difference is a result of a large fraction of *A. inornatus* and an even larger fraction of *A. marmoratus* individuals harboring microbiota dominated by *Dietzia*, *Corynebacterium* and, to a lesser extent, an unclassified Propionibacteriaceae. This finding is further supported by Indicator Species Analysis, which identifies *Dietzia*, *Corynebacterium*, an unclassified Propionibacteriaceae genus, and *Marinilutecoccus* as genus-level markers of parental lizards (see Additional file 7: Indicator_Species_Analysis.xlsx) and an unclassified Propionibacteriaceae strain, *Corynebacterium testudinoris*, and *Dietzia maris* as ASV markers of progenitors. Interestingly, despite being a strong indicator of progenitors, *Dietzia* is still the most abundant microbial genus across hybrid gut samples (see Table 6.2). This is largely a result of a much smaller fraction of hybrid lizard gut samples being dominated by *Dietzia* (see Fig. 5A).

The skin microbiota of all three lizard species are dominated, in order, by Actinobacteria, Proteobacteria and Firmicutes, followed by either Chloroflexi or Bacteriodetes (see Fig. 5C, Table 6.3). Geodermatophilus is the most abundant genus on *A. neomexicanus* skin and is the second and third most abundant genus on *A. marmoratus* and *A. inornatus* skin respectively. Streptomyces is also an abundant genus on all three lizard species. Other dominant genera include Nocardoides on *A. inornatus* and *A. neomexicanus* as well as Fodinibacter on *A. marmoratus* and *A. neomexicanus* (see Fig. 5C, Additional file 6: Table 6.4). Similar to gut microbiota, Indicator Species Analysis identifies *Dietzia* as a strong genus-level marker of parental skin microbiota and *Dietzia maris* and *Corynebacterium testudinoris* as ASV markers. An unclassified Rhodospirillales emerges as a strong genus-level marker of hybrid skin microbiota, while an ASV that maps to *Geodermatophilus siccatus* emerges as an ASV marker of hybrids; although the latter is only marginally significant after correction for multiple comparisons. Interestingly, the next two most significant ASV markers of hybrids, neither of which reach significance after correction for multiple comparisons, map to a different strain of *Geodermatophilus siccatus* and a strain of *Geodermatophilus ruber* (see Additional file 7: Indicator_Species_Analysis.xlsx). Note that *Dietzia* and *Corynebacterium* are two of the same markers of the progenitor population in gut microbiota as well. Several additional genera and ASVs emerge as markers of individual lizard species, but not of hybrid lizards relative to progenitor lizards (see Additional file 7: Indicator_Species_Analysis.xlsx).

**Indicator Species Analysis**: Indicator species analysis was performed on both ASVs and microbial genera using the multipatt function from the *indicspecies* package (version 1.7.12)^1^ and using 50,000 permutations for the permutational test of significance. Indicator species were identified based on having a Benjamini-Hochberg corrected *p* <0.05 for the permutational test. However, to avoid loss of statistical power due to correction for multiple comparisons for the ASV analysis, we only considered and performed Benjamini-Hochberg corrections based on the p-values of microbial ASVs that constituted at least 1% of the microbiota on at least one animal (this reduced to total number of ASVs from 7842 to 532 for gut microbiota and from 15046 to 190 for skin microbiota). Because there were fewer microbial genera, for analysis of microbial genera, we considered all taxa, regardless of their relative abundances on individual animals or across the whole population.

**Table 6.1.** The five most abundant microbial phyla in lizard gut microbiota across (i) all lizards, (ii) *Aspidoscelis neomexicanus* lizards from both locations, (iii) *A. neomexicanus* from SNW, (iv) *A. neomexicanus* from SBluG, (v) *A. marmoratus* from SNW, and (vi) *A. inornatus* from SBluG. The average percentage of the gut microbiota made up by each of the most abundant phyla is shown underneath the phylum name.

|  | 1 | 2 | 3 | 4 | 5 |
| --- | --- | --- | --- | --- | --- |
| ALL | Actinobacteria  50.900480 | Firmicutes 20.623075 | Proteobacteria 15.077177 | Bacteroidetes 7.208224 | Chloroflexi 1.297333 |
| NEO (ALL) | Actinobacteria 39.307705 | Firmicutes 25.772602 | Proteobacteria 20.736571 | Bacteroidetes 7.552366 | Chloroflexi 1.497925 |
| NEO (SNW) | Actinobacteria 44.069277 | Proteobacteria 30.646046 | Firmicutes 15.963228 | Bacteroidetes 3.728478 | Acidobacteria 1.239779 |
| NEO (SBluG) | Actinobacteria 35.548570 | Firmicutes 33.516845 | Proteobacteria 12.913302 | Bacteroidetes 10.571225 | Chloroflexi 1.811992 |
| MARM (SNW) | Actinobacteria 73.210613 | Proteobacteria 11.039635 | Firmicutes 8.413645 | Bacteroidetes 2.116029 | Chloroflexi 1.609877 |
| INO (SBluG) | Actinobacteria 53.224992 | Firmicutes 21.889761 | Bacteroidetes 11.569119 | Proteobacteria 7.088508 | Cyanobacteria 2.920730 |

**Table 6.2.** The five most abundant microbial genera in lizard gut microbiota across (i) all lizards, (ii) *Aspidoscelis neomexicanus* lizards from both locations, (iii) *A. neomexicanus* from SNW, (iv) *A. neomexicanus* from SBluG, (v) *A. marmoratus* from SNW, and (vi) *A. inornatus* from SBluG. The average percentage of the gut microbiota made up by each of the most abundant genera is shown underneath the genus name.

|  | 1 | 2 | 3 | 4 | 5 |
| --- | --- | --- | --- | --- | --- |
| ALL | Dietzia 14.342919 | Corynebacterium 8.483048 | unclassified_  Lachnospiraceae 4.517174 | Geodermatophilus 4.192466 | Salmonella 3.425886 |
| NEO (ALL) | Dietzia 7.572756 | Salmonella 6.017917 | Geodermatophilus 4.565254 | unclassified_  Lachnospiraceae 4.129209 | Bacteroides 2.335508 |
| NEO (SNW) | Salmonella 11.963533 | Geodermatophilus 6.124232 | Dietzia 4.940830 | Fodinibacter 4.727005 | Blautia 2.616690 |
| NEO (SBluG) | Dietzia 9.650593 | unclassified_  Lachnospiraceae 6.826147 | Bacteroides 3.422296 | Geodermatophilus 3.334483 | Parabacteroides 3.211785 |
| MARM (SNW) | Dietzia 30.105801 | Corynebacterium 12.072890 | Brevibacterium 6.020018 | Geodermatophilus 5.325309 | unclassified_  Lachnospiraceae 3.267370 |
| INO (SBluG) | Corynebacterium 21.878333 | Dietzia 12.966631 | unclassified_  Propionibacteriaceae 7.553234 | unclassified_  Lachnospiraceae 6.591403 | Odoribacter 3.197375 |

**Table 6.3.** The five most abundant microbial phyla on lizard skin microbiota across (i) all lizards, (ii) *A. neomexicanus* lizards from both locations, (iii) *A. neomexicanus* from SNW, (iv) *A. neomexicanus* from SBluG, (v) *A. marmoratus* from SNW, and (vi) *A. inornatus* from SBluG. The average percentage of the skin microbiota made up by each of the most abundant phyla is shown underneath the phylum name.

|  | 1 | 2 | 3 | 4 | 5 |
| --- | --- | --- | --- | --- | --- |
| ALL | Actinobacteria 58.683958 | Proteobacteria 18.040830 | Firmicutes 7.380724 | Bacteroidetes 4.024846 | Chloroflexi 3.610163 |
| NEO (ALL) | Actinobacteria 59.528671 | Proteobacteria 19.642202 | Firmicutes 4.377946 | Chloroflexi 4.055408 | Bacteroidetes 3.032910 |
| NEO (SNW) | Actinobacteria 64.857680 | Proteobacteria 17.555481 | Firmicutes 4.680139 | Chloroflexi 3.220874 | Bacteroidetes 2.689050 |
| NEO (SBluG) | Actinobacteria 55.321559 | Proteobacteria 21.289613 | Chloroflexi 4.714251 | Firmicutes 4.139373 | Bacteroidetes 3.304378 |
| MARM (SNW) | Actinobacteria 68.539224 | Proteobacteria 15.189072 | Firmicutes 4.085600 | Chloroflexi 2.813226 | Bacteroidetes 2.401644 |
| INO (SBluG) | Actinobacteria 47.649629 | Proteobacteria 17.311438 | Firmicutes 16.850803 | Bacteroidetes 7.654461 | Chloroflexi 3.411146 |

**Table 6.4.** The five most abundant microbial genera on lizard skin microbiota across (i) all lizards, (ii) *A. neomexicanus* lizards from both locations, (iii) *A. neomexicanus* from SNW, (iv) *A. neomexicanus* from SBluG, (v) *A. marmoratus* from SNW, and (vi) *A. inornatus* from SBluG. The average percentage of the skin microbiota made up by each of the most abundant genera is shown underneath the genus name.

|  | 1 | 2 | 3 | 4 | 5 |
| --- | --- | --- | --- | --- | --- |
| ALL | Geodermatophilus 10.014940 | Fodinibacter 7.637811 | Nocardioides 5.108983 | Streptomyces 4.755007 | Rubrobacter 3.411058 |
| NEO (ALL) | Geodermatophilus 10.918053 | Nocardioides 6.087803 | Streptomyces 5.921441 | Fodinibacter 4.164724 | Rubrobacter 3.716244 |
| NEO (SNW) | Geodermatophilus 13.974965 | Fodinibacter 9.438615 | Streptomyces 8.970475 | Nocardioides 3.675444 | Rubrobacter 3.252345 |
| NEO (SBluG) | Geodermatophilus 8.504701 | Nocardioides 7.992298 | Rubrobacter 4.082480 | Streptomyces 3.514308 | Microvirga 2.683994 |
| MARM (SNW) | Fodinibacter 23.614724 | Geodermatophilus 13.535333 | Rubrobacter 3.012660 | Arthrobacter 2.627475 | Streptomyces 2.620577 |
| INO (SBluG) | Nocardioides 5.601648 | Dietzia 4.816035 | Geodermatophilus 4.795456 | Streptomyces 4.277362 | unclassified_  Lachnospiraceae 3.925766 |
