## Additional file 9 for "Transgressive Hybrids as Hopeful Holobionts"

**Additional file 9: *Aspidoscelis* Habitat Niches**

Several pieces of information suggest that *A. neomexicanus* has a broader habitat niche (including novel habitat classes) from its progenitor species. First, in a study from the same general region of New Mexico where our research occurred, Axtell^1^ found that *A. neomexicanus* was distributed across the entire habitat gradient that he studied, ranging from juniper grasslands in the West to pinon juniper forests in the East. By contrast, *A. inornatus* was missing from the middle of the gradient that bisects the Rio Grande. Overall, this resulted in *A. neomexicanus* being present in two additional discrete habitat classes as compared to *A. inornatus* and 33% more aerial extent. Second, current NatureServe models^2^ indicate that *A. neomexicanus* occupies six distinct habitat classes, including two that are not inhabited by either progenitor. While this is a slightly narrower niche breadth than *A. inornatus*, which occupies seven habitat classes, it is much broader than *A. marmoratus*, which occupies three habitat classes.

Beyond this existing literature, we used georeferenced occurrence records from the Global Biodiversity Information Facility^3^ for *A. inornatus*, *A. marmoratus*, and *A. neomexicanus*, combined with Landfire vegetation classes^4^, to determine the niche breadth of each species. More specifically, we considered counts of the number of vegetation classes, as well as the Shannon index applied to the proportion of occurrences of each species in each vegetation class. The latter helps to address evenness of a species distribution across vegetation classes. Briefly, we considered four different levels of vegetation classification (landfire pixels, GROUP, MACROGROUP, and CLASS). For each level, we subsampled 500 individual lizards of each species and either counted the number of vegetation classes where the species occurred or calculated the Shannon index. We then repeated the subsampling process 100 times. To test for statistically significant differences between lizard species, we used a Kruskal-Wallis test applied to the three distinct species. This was done using the kruskal.test function from the stats package (version 4.2.1).^5^ When the Kruskal-Wallis test was significant, we did pairwise post-hoc testing by applying pairwise Wilcoxon rank sum also from the stats package. Results are shown in Figure 10.1. Like NatureServe, counts of habitat classes suggest that, except at the highest level of classification, *A. neomexicanus* is intermediate in habitat niche breadth (see Fig. 10.1A-D), generally much broader than *A. marmoratus* and almost as broad as *A. inornatus*. However, accounting for evenness like the study by Axtell,^1^ the Shannon Index suggests that at all levels of classification, *A. neomexicanus* is transgressive, exhibiting a broader habitat niche (see Fig. 10.1E-H). Comparing the identities of the habitats across the three lizard species further shows that, except at the coarsest level of classification, *A. neomexicanus* occupies numerous novel habitat classes where neither *A. neomexicanus* nor *A. marmoratus* are found (see Figure 10.2). For the same classification levels, *A. neomexicanus* occupies more unique habitat classes than either progenitor, another example of habitat niche transgression.

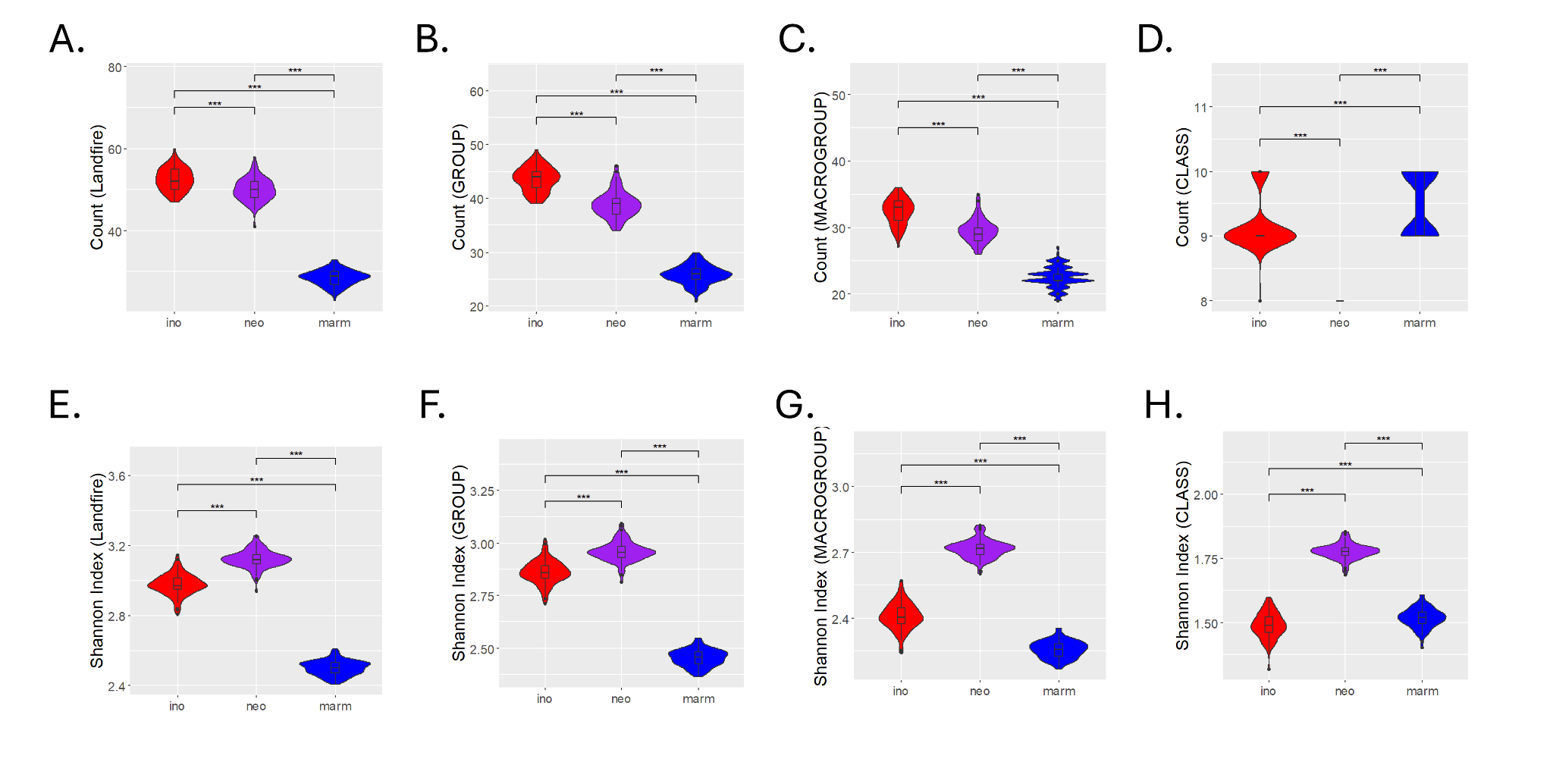

**Figure 9.1** Violin plots for the counts of vegetation classes (A-D) and the Shannon Index applied to the proportions of each species in each vegetation class (E-H) comparing *A. inornatus* (red), *A. neomexicanus* (purple) and *A. marmoratus* (blue). Statistical differences are based on 100 subsamples of 500 individual lizards from each species. From left to right, each column resembles a different resolution of Landfire data from the highest resolution, pixel values (A, E), to the lowest resolution, ‘CLASS’ (D, H). Significant differences in diversity between groups, as determined by a Kruskal-Wallis test followed by post hoc Pairwise Wilcox tests using a Benjamini-Hochberg correction, are indicated as follows: p-value ≤ 0.001 (***), p-value ≤ 0.01 (**), p-value ≤ 0.05 (*), p-value ≤ 0.1 (.)

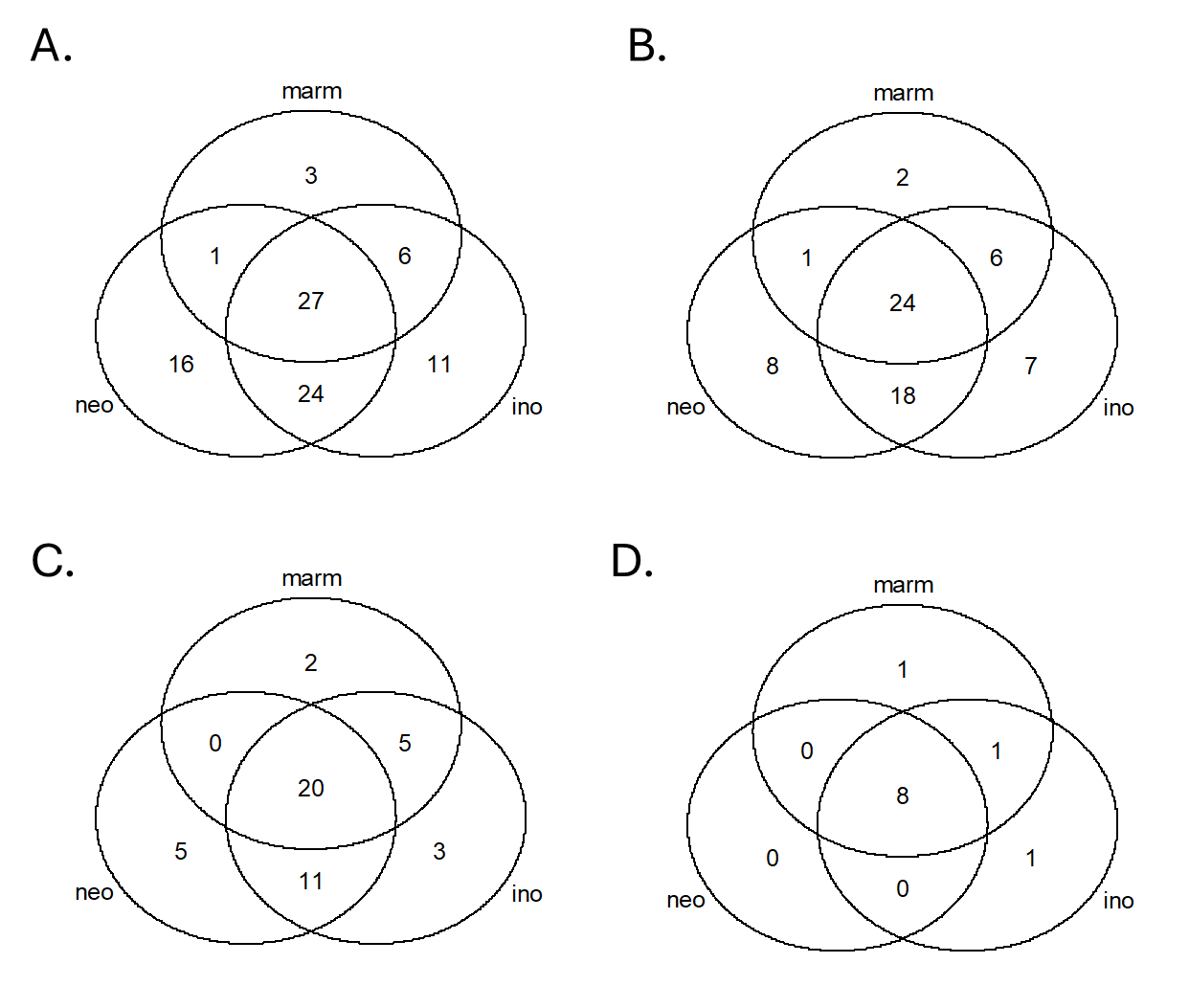

**Figure 9.2** Venn diagrams showing the shared and unique habitat classifications for each of our three lizard species based on Landfire vegetation classes at the (A) highest-resolution pixel values, (B) high-resolution ‘GROUP,’ (C) medium-resolution ‘MACROGROUP,’ and (D) low-resolution ‘CLASS’.
