## Additional file 10 for "Transgressive Hybrids as Hopeful Holobionts"

**Additional file 10: Marking *Aspidoscelis* Lizards**

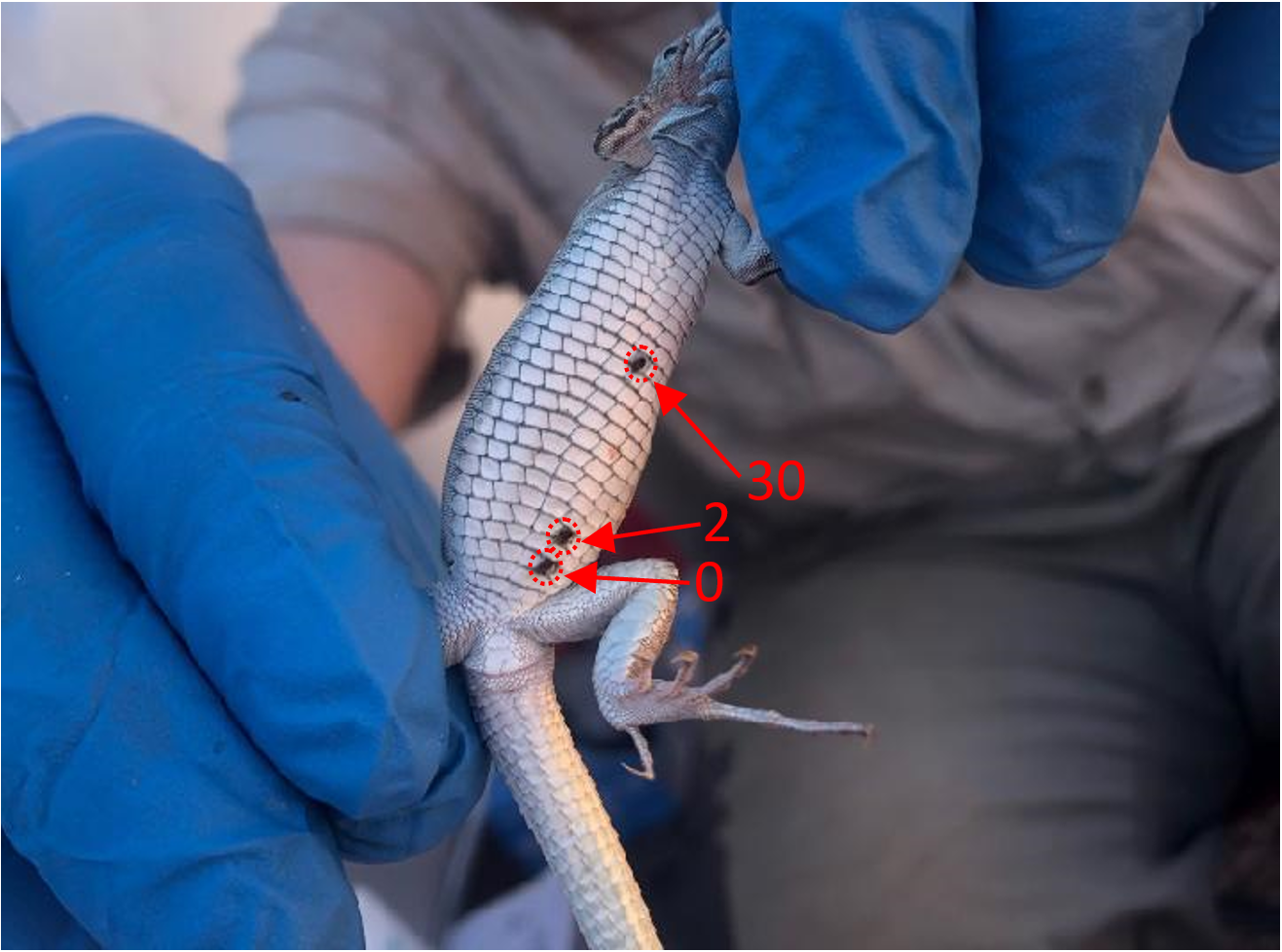

**Figure 10.1.** Ventral aspect of adult *Aspidoscelis neomexicanus* (#32) marked with a miniature medical cautery unit. The marking scheme consists of marking individuals on the same ventral scale column (along the anterior-posterior axis). The most posterior mark near the left hind-leg indicates ‘0’ and is used to calibrate the marks positioned anteriorly. Scale-by-scale, counting anterior to the calibration mark, scales indicated 1-10 by increments of one. At the 11^th^ scale anterior to the calibration mark, each scale indicates increments of 10 (i.e., 20, 30, 40, 50,… n).
