## Additional file 1 for "Transgressive Hybrids as Hopeful Holobionts"

**Additional File 1: Additional Diversity Metrics**

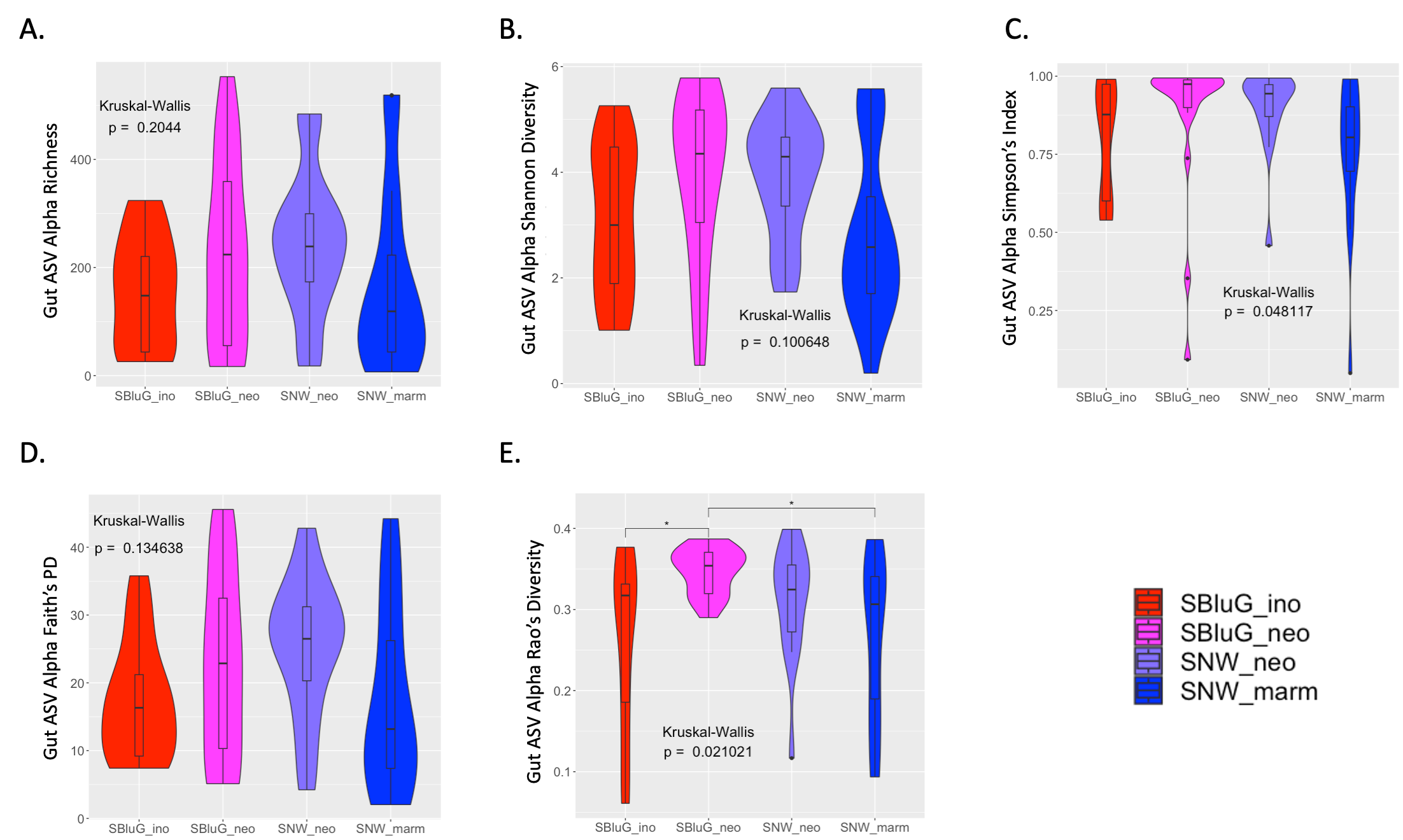

**Figure 1.1.** Comparison of gut microbiota 𝛼-diversity of amplicon sequence variants (ASVs) between populations of *Aspidoscelis inornatus* from SBluG (red), *A. neomexicanus* from SBluG (magenta), *A. neomexicanus* from SNW (purple), and *A. marmoratus* from SNW (blue) as measured by (A) richness (count of ASVs), (B) Shannon diversity, (C) Simpson’s index, (D) Faith’s phylogenetic diversity (PD), and (E) Rao’s diversity. Significant differences in diversity between groups, as determined by a Kruskal-Wallis test followed by post hoc pairwise Wilcox tests using a Benjamini-Hochberg correction, are indicated as follows: p-value ≤ 0.001 (***), p-value ≤ 0.01 (**), p-value ≤ 0.05 (*), p-value ≤ 0.1 (.).

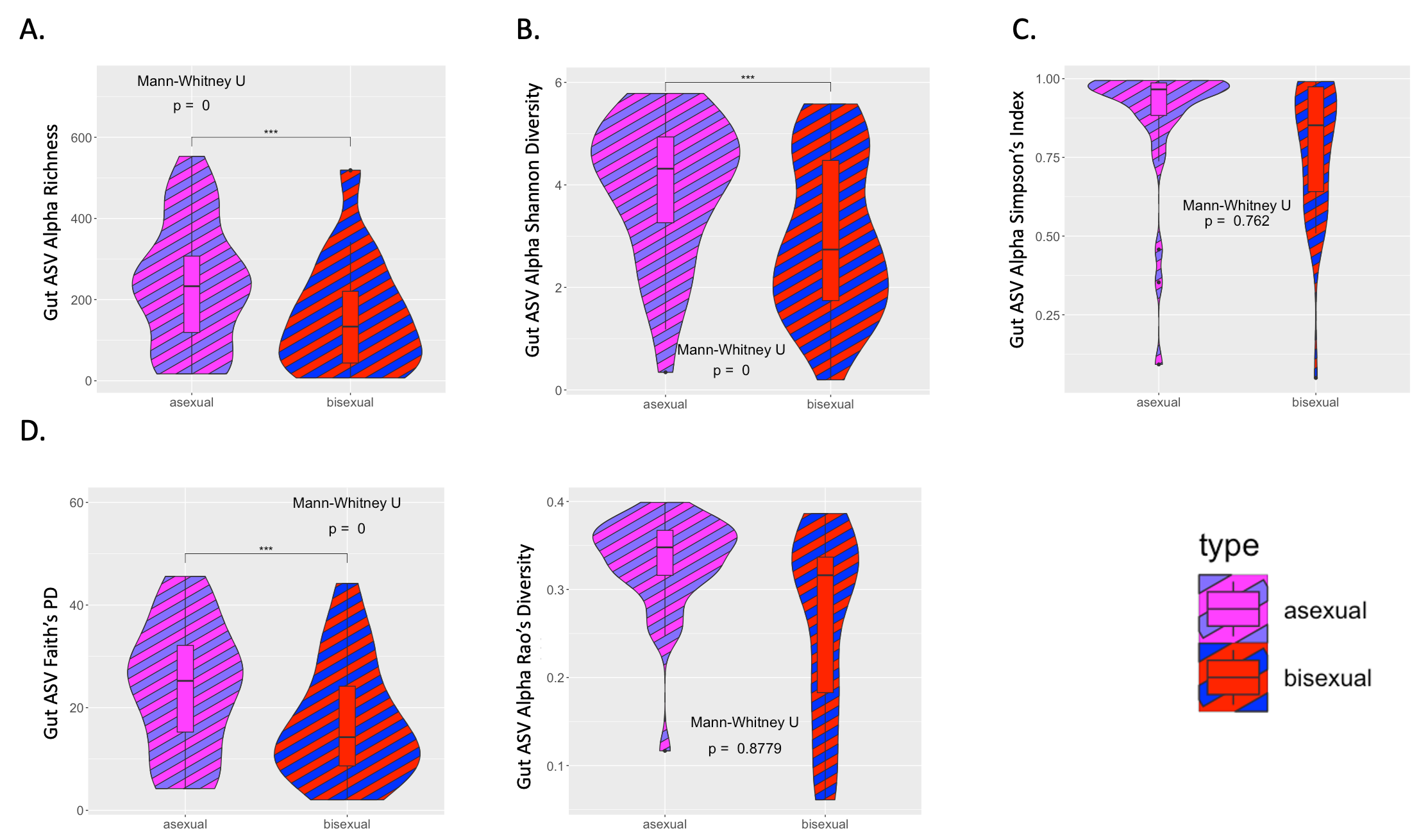

**Figure 1.2.** Comparison of gut microbiota 𝛼-diversity of amplicon sequence variants (ASVs) between parent sexual species (red and blue striped) and hybrid asexual *Aspidoscelis neomexicanus* (magenta and purple striped) as measured by (A) richness (count of ASVs), (B) Shannon diversity, (C) Simpson’s index, (D) Faith’s phylogenetic diversity (PD), and (E) Rao’s diversity. Significant differences in diversity between groups, as determined by a Mann-Whitney test, are indicated as follows: p-value ≤ 0.001 (***), p-value ≤ 0.01 (**), p-value ≤ 0.05 (*), p-value ≤ 0.1 (.).

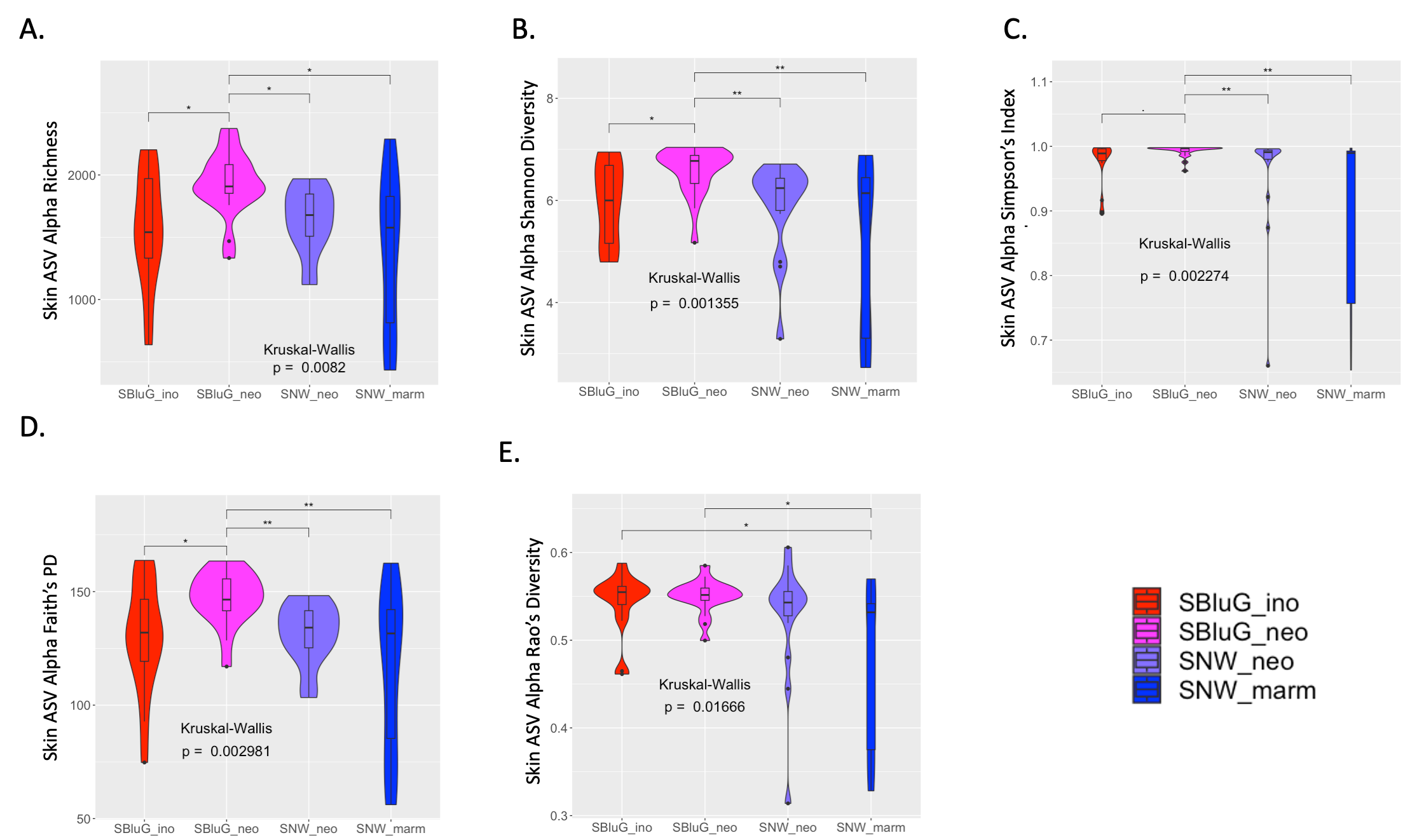

**Figure 1.3.** Comparison of skin microbiota 𝛼-diversity of amplicon sequence variants (ASVs) between populations of *Aspidoscelis inornatus* from SBluG (red), *A. neomexicanus* from SBluG (magenta), *A. neomexicanus* from SNW (purple), and *A. marmoratus* from SNW (blue) as measured by (A) richness (count of ASVs), (B) Shannon diversity, (C) Simpson’s index, (D) Faith’s phylogenetic diversity (PD), and (E) Rao’s diversity. Significant differences in diversity between groups, as determined by a Kruskal-Wallis test followed by post hoc pairwise Wilcox tests using a Benjamini-Hochberg correction, are indicated as follows: p-value ≤ 0.001 (***), p-value ≤ 0.01 (**), p-value ≤ 0.05 (*), p-value ≤ 0.1 (.).

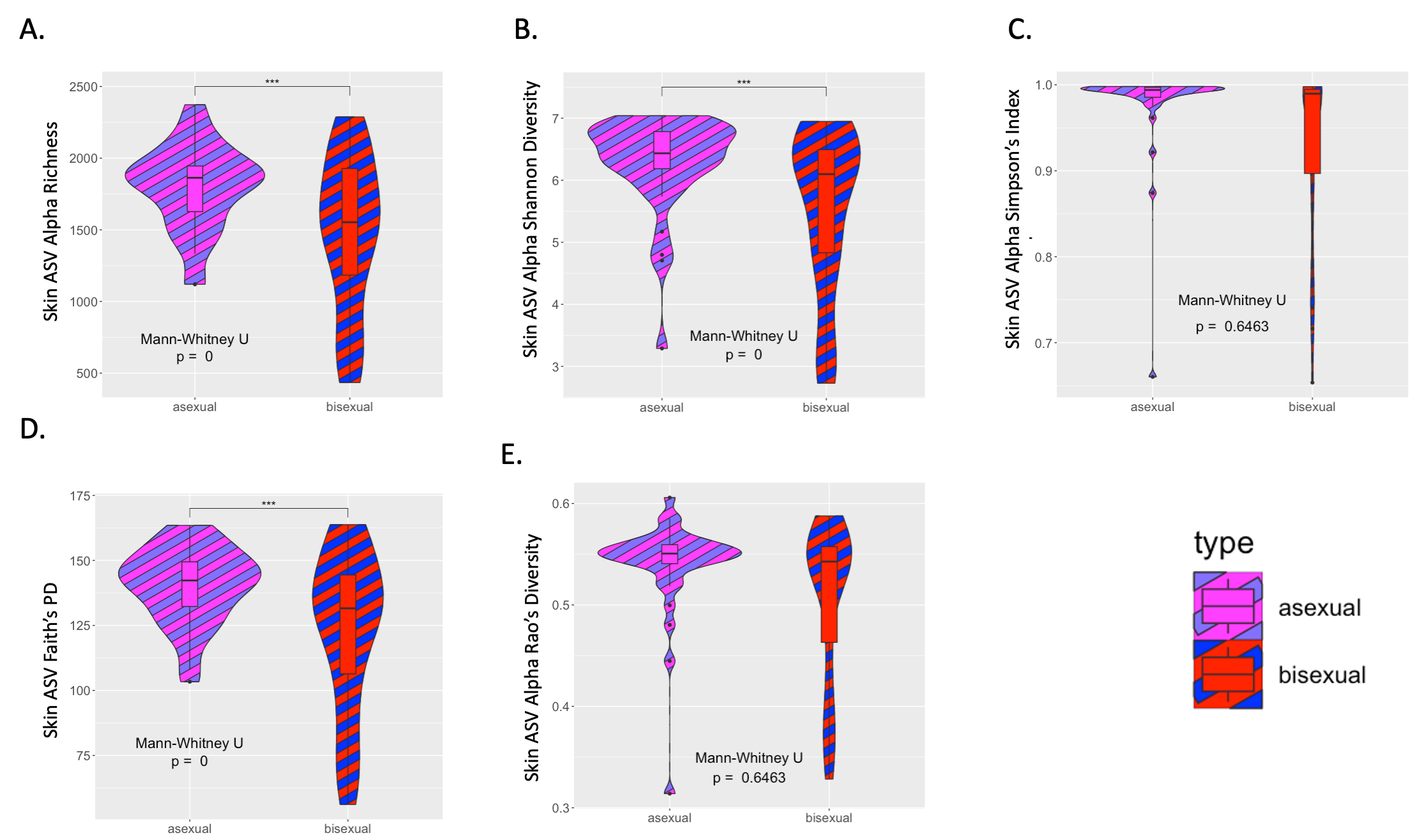

**Figure 1.4.** Comparison of skin microbiota 𝛼-diversity of amplicon sequence variants (ASVs) between parent sexual species (red and blue striped) and hybrid asexual *Aspidoscelis neomexicanus* (magenta and purple striped) as measured by (A) richness (count of ASVs), (B) Shannon diversity, (C) Simpson’s index, (D) Faith’s phylogenetic diversity (PD), and (E) Rao’s diversity. Significant differences in diversity between groups, as determined by a Mann-Whitney test, are indicated as follows: p-value ≤ 0.001 (***), p-value ≤ 0.01 (**), p-value ≤ 0.05 (*), p-value ≤ 0.1 (.).

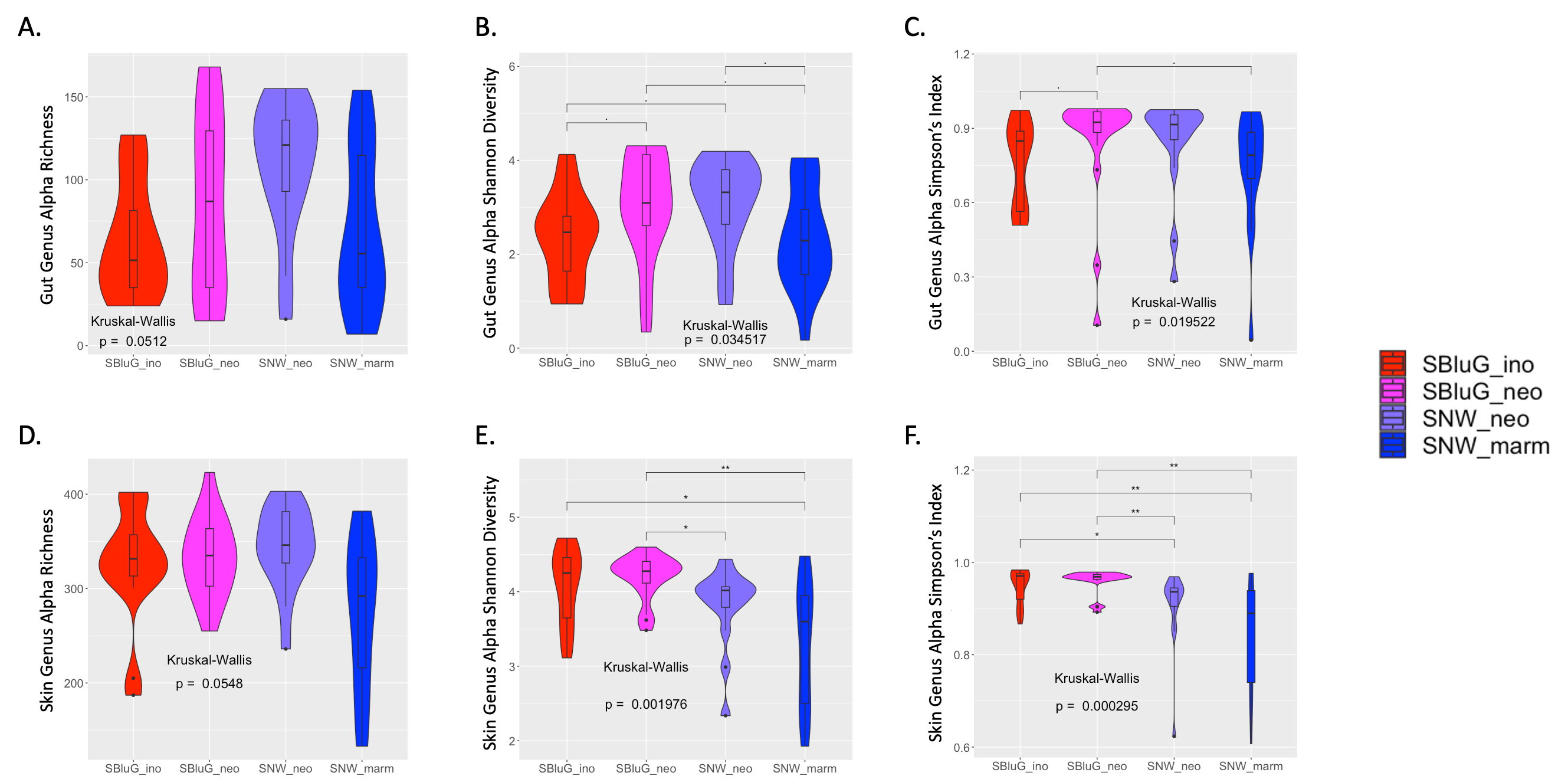

**Figure 1.5.** Comparison of gut (A-C) and skin (D-F) microbiota 𝛼-diversity of microbial genera between populations of *Aspidoscelis inornatus* from SBluG (red), *A. neomexicanus* from SBluG (magenta), *A. neomexicanus* from SNW (purple), and *A. marmoratus* from SNW (blue) as measured by (A, D) richness (count of genera), (B, E) Shannon diversity, and (C, F) Simpson’s index. Significant differences in diversity between groups, as determined by a Kruskal-Wallis test followed by post hoc pairwise Wilcox tests using a Benjamini-Hochberg correction, are indicated as follows: p-value ≤ 0.001 (***), p-value ≤ 0.01 (**), p-value ≤ 0.05 (*), p-value ≤ 0.1 (.).

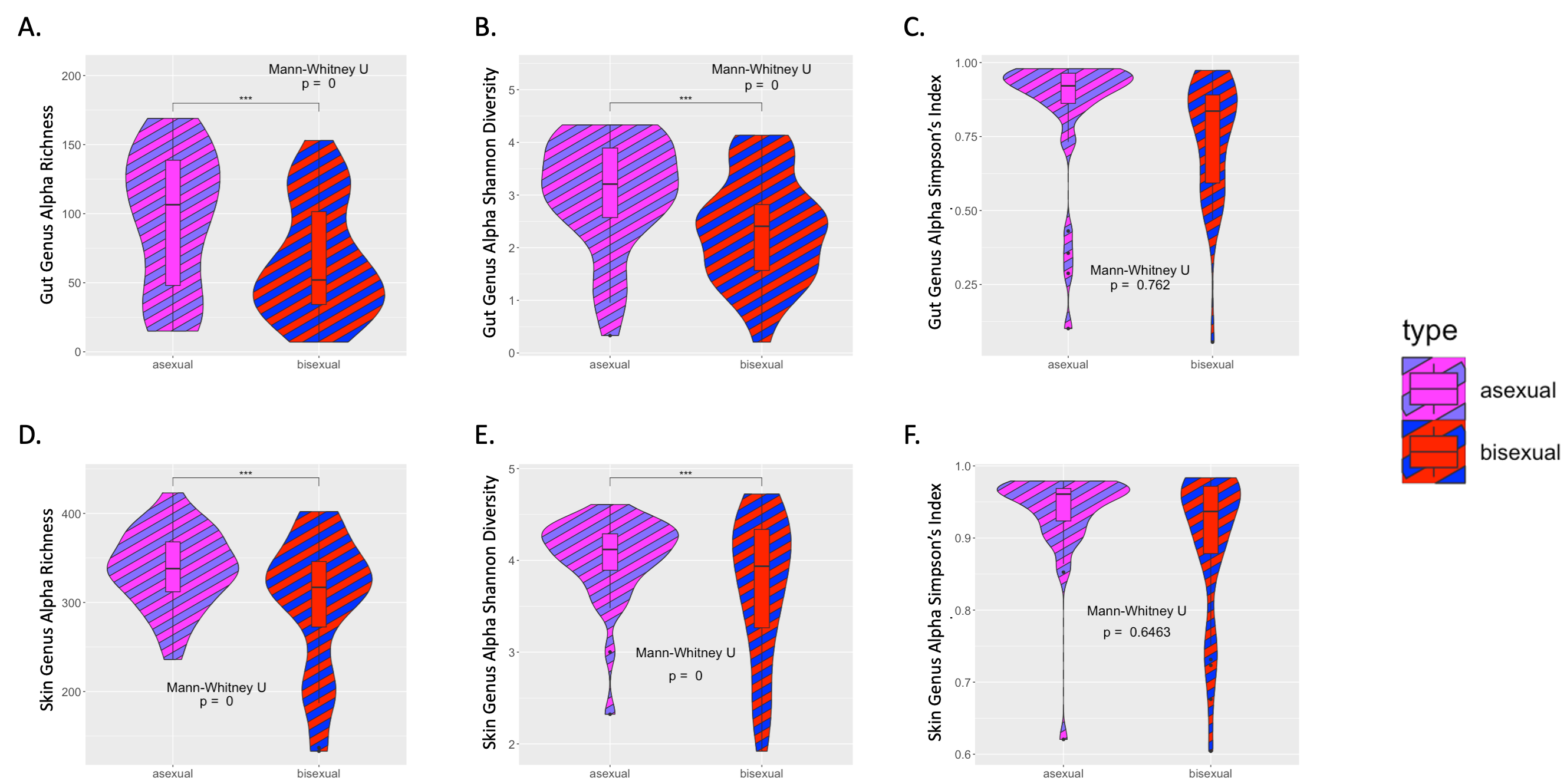

**Figure 1.6.** Comparison of gut (A-C) and skin (D-F) microbiota 𝛼-diversity of microbial genera between parent sexual species (red and blue striped) and hybrid asexual *Aspidoscelis neomexicanus* (magenta and purple striped) as measured by (A, D) richness (count of genera), (B, E) Shannon diversity, and (C, F) Simpson’s index. Significant differences in diversity between groups, as determined by a Mann-Whitney test, are indicated as follows: p-value ≤ 0.001 (***), p-value ≤ 0.01 (**), p-value ≤ 0.05 (*), p-value ≤ 0.1 (.).

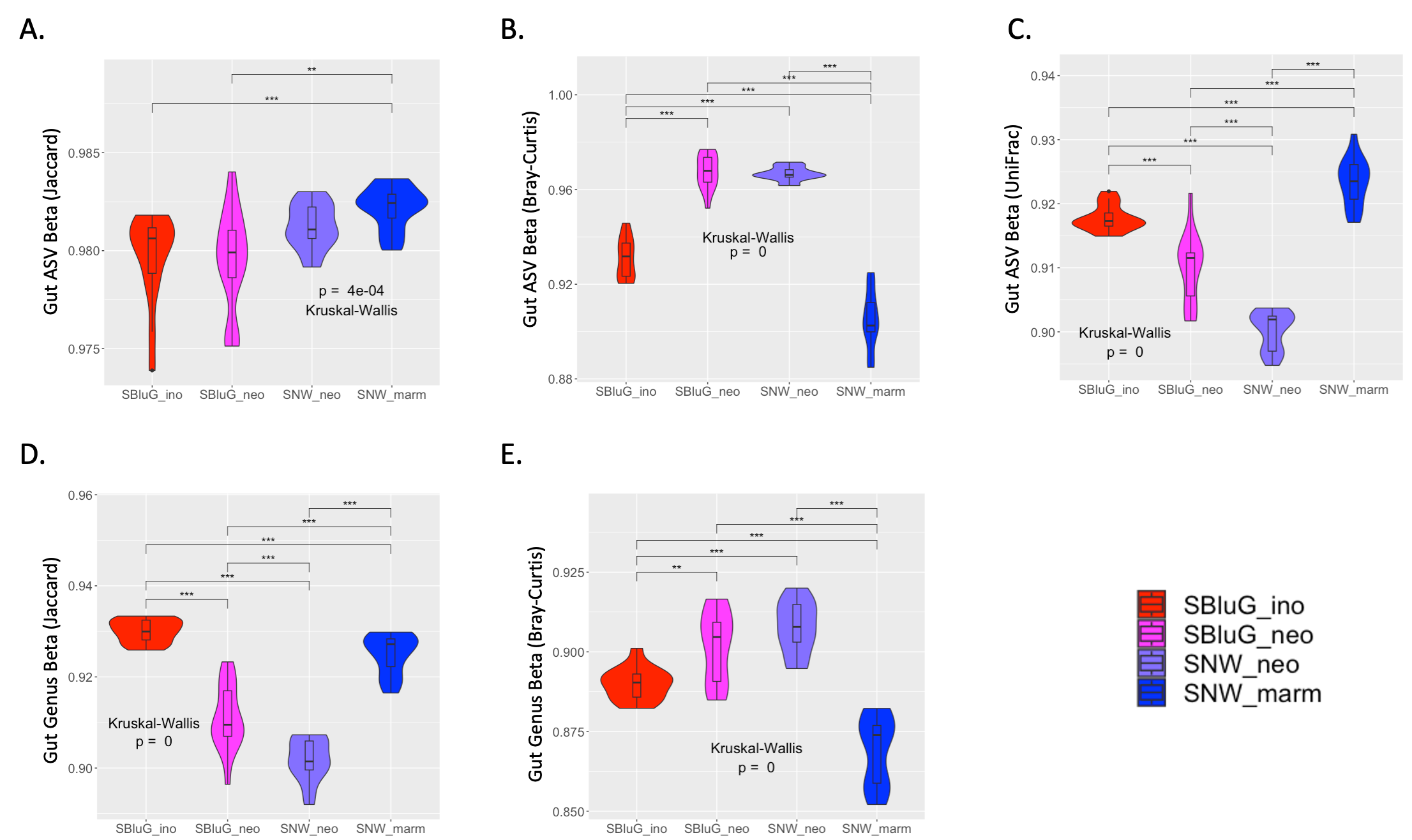

**Figure 1.7.** Comparison of gut microbiota 𝛽-diversity between populations of *Aspidoscelis inornatus* from SBluG (red), *A. neomexicanus* from SBluG (magenta), *A. neomexicanus* from SNW (purple), and *A. marmoratus* from SNW (blue) as measured using (A) Jaccard dissimilarity of amplicon sequence variants (ASVs), (B) Bray-Curtis dissimilarity of ASVs, (C) unweighted UniFrac dissimilarity of ASVs, (D) Jaccard dissimilarity of microbial genera, and (E) Bray-Curtis dissimilarity of microbial genera. Significant differences in diversity between groups, as determined by a Kruskal-Wallis test followed by posthoc pairwise Wilcox tests using a Benjamini-Hochberg correction, are indicated as follows: p-value ≤ 0.001 (***), p-value ≤ 0.01 (**), p-value ≤ 0.05 (*), p-value ≤ 0.1 (.). In all panels, multi-community 𝛽-diversity estimates were generated using 15 subsamples of 12 individuals from each population.

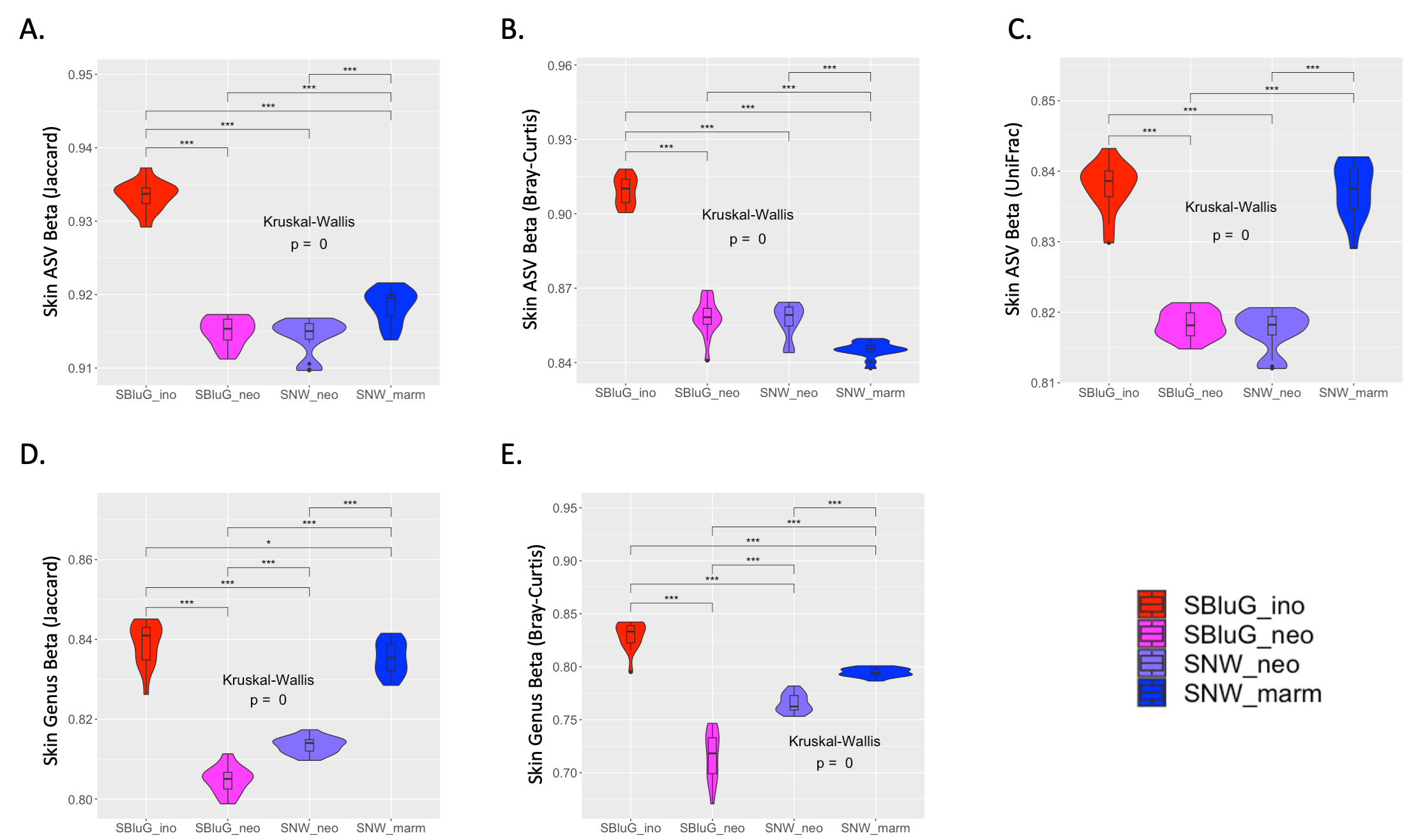

**Figure 1.8.** Comparison of skin microbiota 𝛽-diversity between populations of *Aspidoscelis inornatus* from SBluG (red), *A. neomexicanus* from SBluG (magenta), *A. neomexicanus* from SNW (purple), and *A. marmoratus* from SNW (blue) as measured using (A) Jaccard dissimilarity of amplicon sequence variants (ASVs), (B) Bray-Curtis dissimilarity of ASVs, (C) unweighted UniFrac dissimilarity of ASVs, (D) Jaccard dissimilarity of microbial genera, and (E) Bray-Curtis dissimilarity of microbial genera. Significant differences in diversity between groups, as determined by a Kruskal-Wallis test followed by post hoc pairwise Wilcox tests using a Benjamini-Hochberg correction, are indicated as follows: p-value ≤ 0.001 (***), p-value ≤ 0.01 (**), p-value ≤ 0.05 (*), p-value ≤ 0.1 (.). In all panels, multi-community 𝛽-diversity estimates were generated using 15 subsamples of 12 individuals from each population.

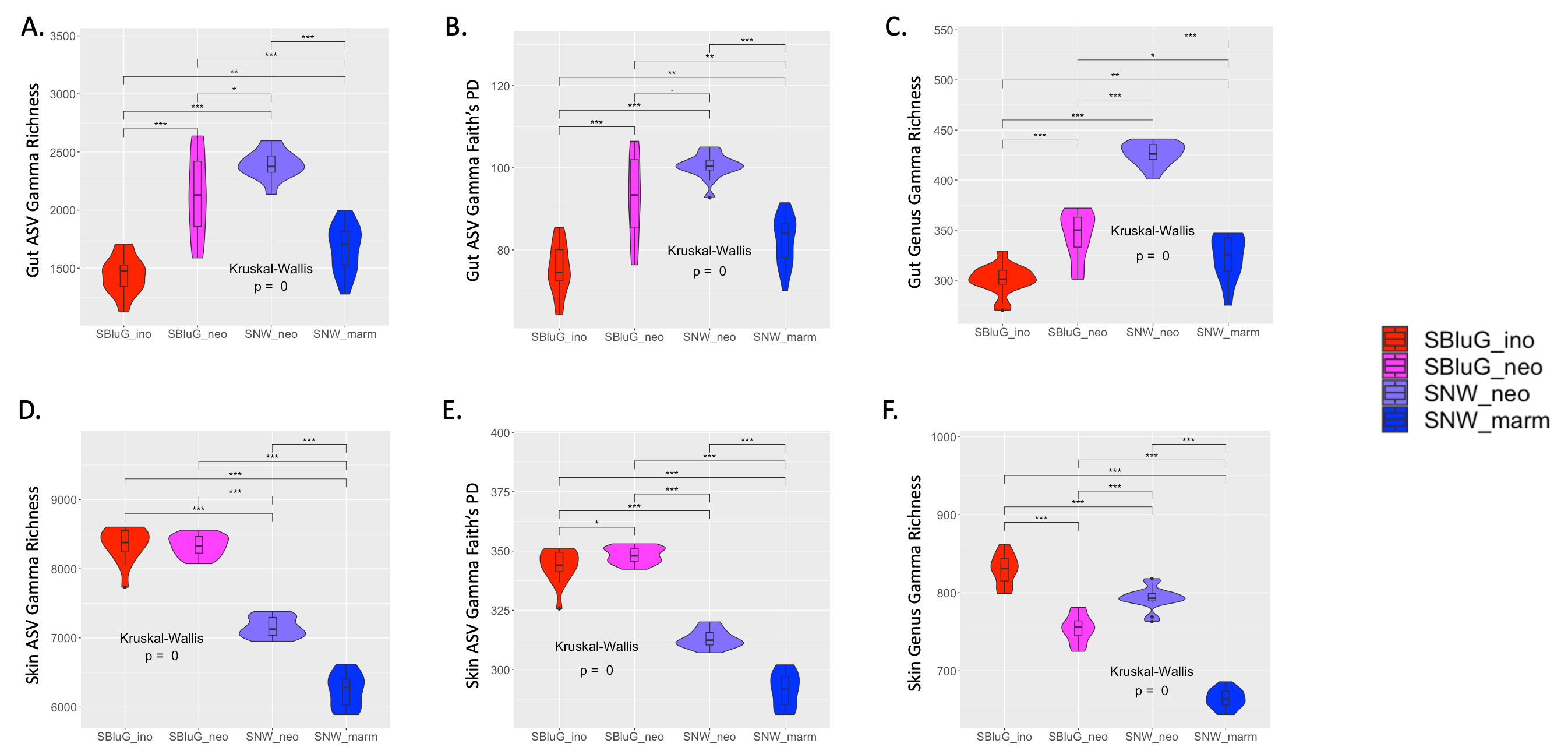

**Figure 1.9.** Comparison of gut (A-C) and skin (D-F) microbiota 𝛾-diversity between populations of *Aspidoscelis inornatus* from SBluG (red), *A. neomexicanus* from SBluG (magenta), *A. neomexicanus* from SNW (purple), and *A. marmoratus* from SNW (blue) as measured using (A, D) richness of amplicon sequence variants (ASV count), (B, E) Faith’s phylogenetic diversity (PD) of ASVs, and (C, F) richness of microbial genera (genera count). Significant differences in diversity between groups, as determined by a Kruskal-Wallis test followed by post hoc pairwise Wilcox tests using a Benjamini-Hochberg correction, are indicated as follows: p-value ≤ 0.001 (***), p-value ≤ 0.01 (**), p-value ≤ 0.05 (*), p-value ≤ 0.1 (.). In all panels, 𝛾-diversity estimates were generated using 15 subsamples of 12 individuals from each population.
