## Additional file 3 for "Transgressive Hybrids as Hopeful Holobionts"

**Additional file 3: Ordination**
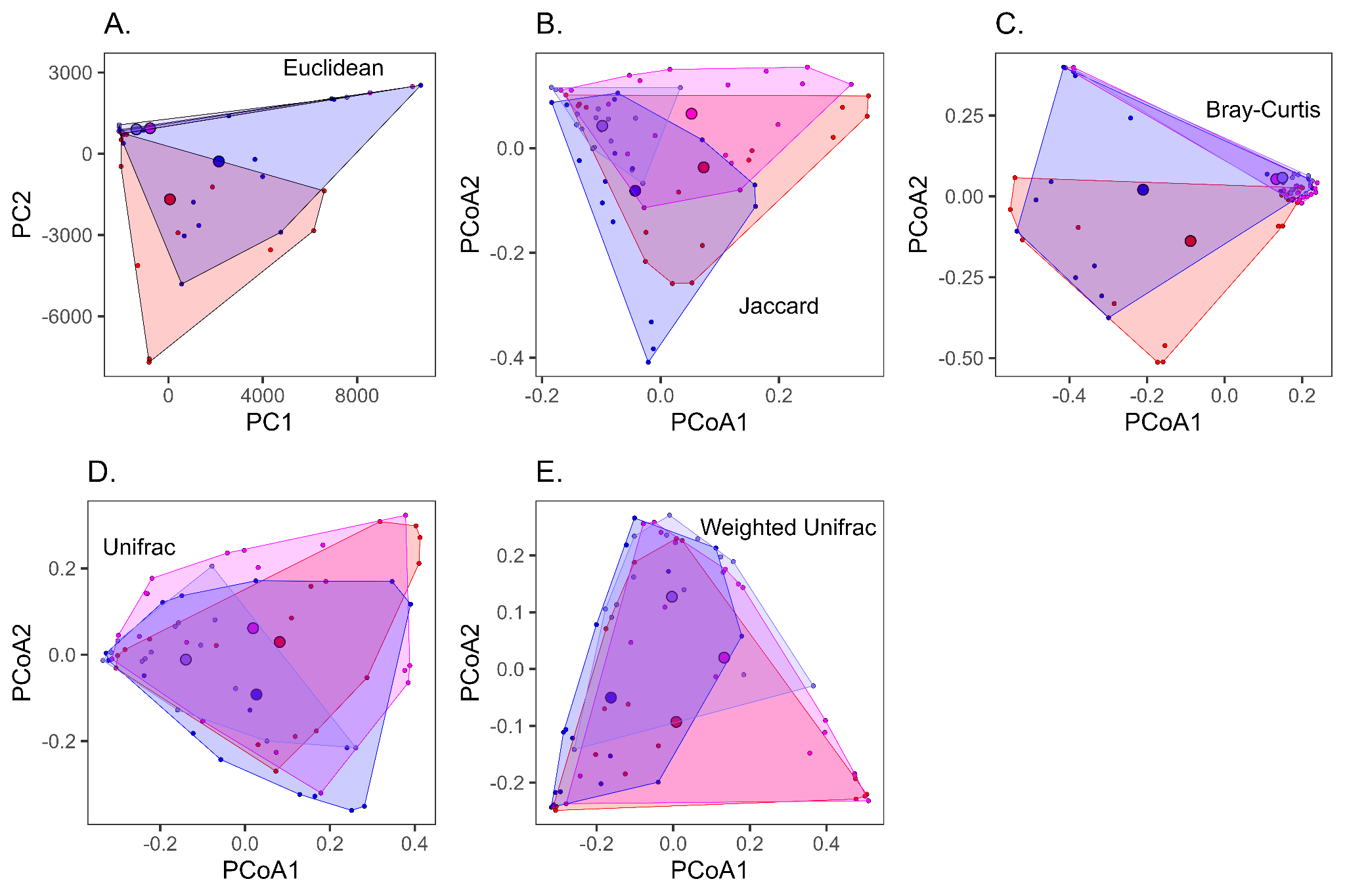

**Figure 3.1.** Two-dimensional Principal Coordinates Analysis (PCoA) of gut microbiota for populations of *A. inornatus* from SBluG (red), *A. neomexicanus* from SBluG (magenta), *A. neomexicanus* from SNW (purple), and *A. marmoratus* from SNW (blue) based on amplicon sequence variants (ASVs) in combination with (A) euclidean dissimilarity, (B) Jaccard dissimilarity, (C) Bray-Curtis dissimilarity, (D) UniFrac dissimilarity, and (E) weighted UniFrac dissimilarity.

**Table 3.1.** Percent variation explained by the first five Principal Coordinate Axes, both individually (%) and cumulatively (Cum%) for PCoA analyses of gut ASVs using Euclidean, Jaccard, Bray-Curtis, UniFrac, and weighted UniFrac distances.

|  | PCoA1 | | PCoA2 | | PCoA3 | | PCoA4 | | PCoA5 | |
| --- | --- | --- | --- | --- | --- | --- | --- | --- | --- | --- |
|  | % | Cum% | % | Cum% | % | Cum% | % | Cum% | % | Cum% |
| Euclidean  (PCA) | 49.98 | 49.98 | 17.53 | 67.51 | 4.27 | 71.78 | 3.38 | 75.16 | 2.87 | 78.03 |
| Jaccard | 4.39 | 4.39 | 3.5 | 7.89 | 2.4 | 10.29 | 2.09 | 12.37 | 1.92 | 14.3 |
| Bray-Curtis | 17.04 | 17.04 | 7.89 | 24.93 | 5.58 | 30.52 | 3.22 | 33.73 | 3 | 36.73 |
| UniFrac | 18.46 | 18.46 | 10.1 | 28.56 | 5.04 | 33.6 | 4 | 37.61 | 2.91 | 40.52 |
| w-UniFrac | 40.04 | 40.04 | 22.07 | 62.11 | 7.05 | 69.16 | 4.1 | 73.27 | 3.59 | 76.85 |

**Table 3.2.** ASVs with the five largest loadings on the first (PCA1) and second (PCA2) Principal Component Axes (Euclidean PCoA) for lizard gut microbiota.

|  | Rank | Gut ASV | Loading | %Loading |
| --- | --- | --- | --- | --- |
| P  C  A  1 | 1 | *Dietzia maris* | 0.987407 | 97.50 |
|  | 2 | *Corynebacterium testudinoris* | 0.134899 | 1.82 |
|  | 3 | *Brevibacterium linens-salitolerans"* | 0.042395 | 0.18 |
|  | 4 | *Salmonella enterica* | -0.025747 | 0.07 |
|  | 5 | *Corynebacterium glyciniphilum"* | 0.019802 | 0.04 |
| P  C  A  2 | 1 | *Corynebacterium testudinoris* | -0.970945 | 94.27 |
|  | 2 | Propionibacteriaceae sp7816-sp7836 | -0.149714 | 2.24 |
|  | 3 | *Dietzia maris* | 0.139948 | 1.96 |
|  | 4 | *Brevibacterium linens-salitolerans"* | -0.063281 | 0.40 |
|  | 5 | *Salmonella enterica* | 0.045693 | 0.21 |

**Table 3.3** Positions of the hybrid gut microbiota from SBluG along the first two ASV PCoA axes. Column 4-5 show p-values for Mann-Whitney U tests assessing whether the positions of hybrid animals along the first PCoA axis is significantly different than the positions of *A. inornatus* (ino) and *A. marmoratus* (marm) respectively. Column 9-10 show p-values for Mann-Whitney U tests assessing whether the position of the hybrid animals along the second PCoA axis is significantly different than the positions of *A. inornatus* (ino) and *A. marmoratus* (marm) respectively. Yellow is used for tests significantly different at *p*>0.05.

|  | PCoA axis 1  mean | | | p-value | | PCoA axis 2  mean | | | p-value | |
| --- | --- | --- | --- | --- | --- | --- | --- | --- | --- | --- |
|  | neo | ino | marm | ino | marm | neo | ino | marm | ino | marm |
| euclidean | 779 | -63 | -2138 | 0.0047 | 0.0042 | 942 | -1681 | -284 | 0 | 0.4221 |
| jaccard | 0.052 | 0.073 | -0.043 | 0.6827 | 0.0519 | 0.066 | -0.037 | -0.081 | 0.0081 | 0.001 |
| bray | 0.133 | -0.088 | -0.21 | 0.0012 | 8e-04 | 0.053 | -0.138 | 0.021 | 4e-04 | 0.8318 |
| unifrac | 0.019 | 0.082 | 0.027 | 0.5021 | 0.987 | 0.061 | 0.03 | -0.092 | 0.6121 | 0.0165 |
| wunifrac | 0.133 | 0.008 | -0.162 | 0.1332 | 2e-04 | 0.02 | -0.093 | -0.05 | 0.0656 | 0.2432 |

**Table 3.4** Positions of the hybrid gut microbiota from SNW along the first two ASV PCoA axes. Column 4-5 show p-values for Mann-Whitney U tests assessing whether the positions of hybrid animals along the first PCoA axis is significantly different than the positions of *A. inornatus* (ino) and *A. marmoratus* (marm) respectively. Column 9-10 show p-values for Mann-Whitney U tests assessing whether the position of the hybrid animals along the second PCoA axis is significantly different than the positions of *A. inornatus* (ino) and *A. marmoratus* (marm) respectively. Yellow is used for tests significantly different at *p*>0.05.

|  | PCoA axis 1  mean | | | p-value | | PCoA axis 2  mean | | | p-value | |
| --- | --- | --- | --- | --- | --- | --- | --- | --- | --- | --- |
|  | neo | ino | marm | ino | marm | neo | ino | marm | ino | marm |
| euclidean | 1362 | -63 | -2138 | 0.0012 | 9e-04 | 902 | -1681 | -284 | 0 | 0.3183 |
| jaccard | -0.098 | 0.073 | -0.043 | 0.0042 | 0.0597 | 0.043 | -0.037 | -0.081 | 0.2641 | 0.0171 |
| bray | 0.149 | -0.088 | -0.21 | 0.0042 | 0.0015 | 0.058 | -0.138 | 0.021 | 0 | 0.4232 |
| unifrac | -0.14 | 0.082 | 0.027 | 0.0135 | 0.0544 | -0.011 | 0.03 | -0.092 | 0.5196 | 0.2814 |
| wunifrac | -0.003 | 0.008 | -0.162 | 0.4945 | 0.0055 | 0.127 | -0.093 | -0.05 | 8e-04 | 0.0082 |

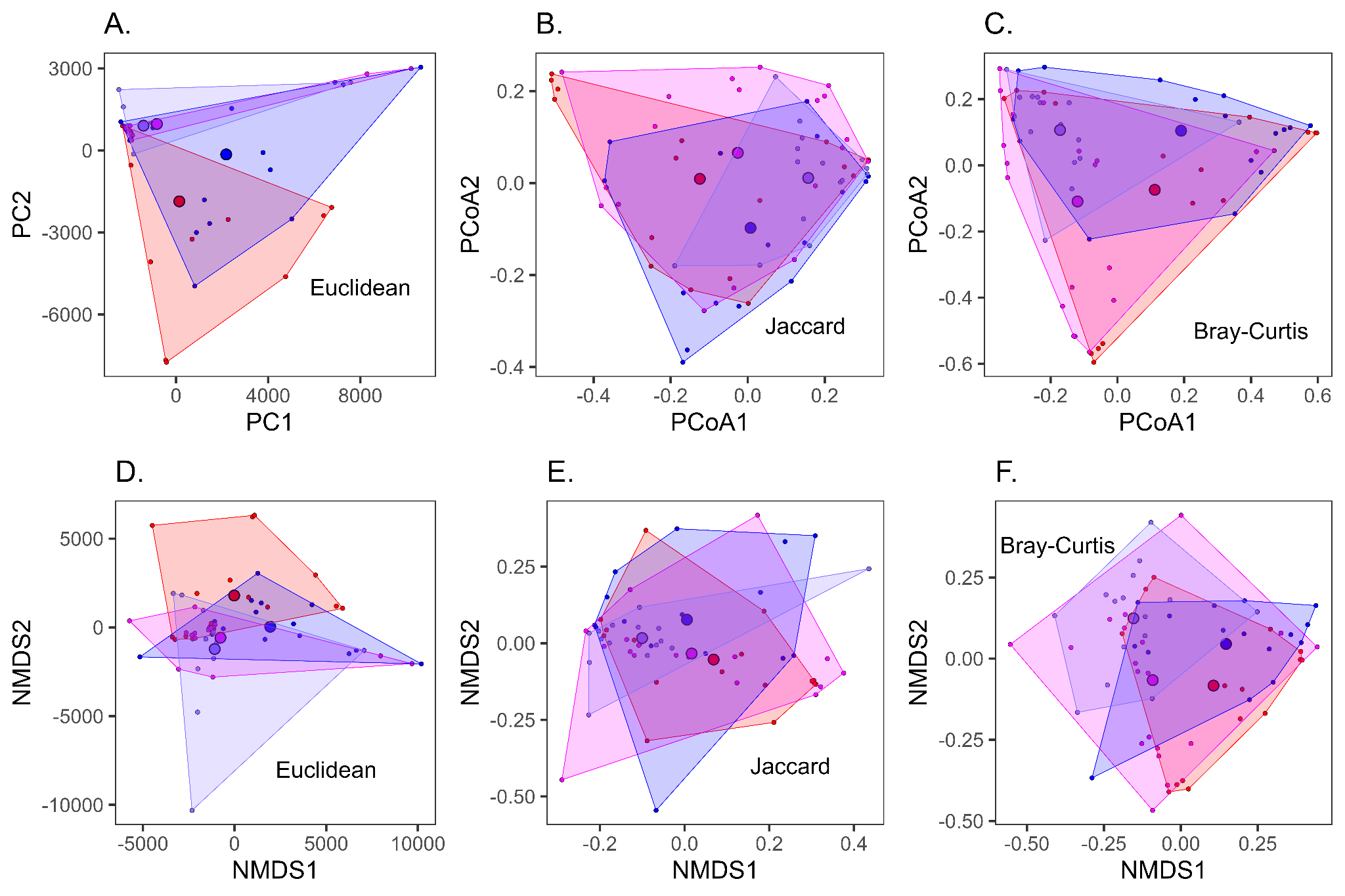

**Figure 3.2.** Two-dimensional Principal Coordinates Analysis of gut microbiota for populations of *A. inornatus* from SBluG (red), *A. neomexicanus* from SBluG (magenta), *A. neomexicanus* from SNW (purple), and *A. marmoratus* from SNW (blue) and based on microbial genera in combination with (A) euclidean dissimilarity, (B) Jaccard dissimilarity, and (C) Bray-Curtis dissimilarity.

**Table 3.5.** Percent variation explained by the first five Principal Coordinate Axes, both individually (%) and cumulatively (Cum%) for PCoA analyses of gut genera using Euclidean, Jaccard, and Bray-Curtis distances.

|  | PCoA1 | | PCoA2 | | PCoA3 | | PCoA4 | | PCoA5 | |
| --- | --- | --- | --- | --- | --- | --- | --- | --- | --- | --- |
|  | % | Cum% | % | Cum% | % | Cum% | % | Cum% | % | Cum% |
| Euclidean  (PCA) | 43.74 | 43.74 | 16.76 | 60.51 | 9.37 | 69.87 | 5.74 | 75.61 | 3.57 | 79.18 |
| Jaccard | 17.61 | 17.61 | 7.94 | 25.55 | 3.99 | 29.54 | 3.32 | 32.86 | 3.05 | 35.91 |
| Bray-Curtis | 25.41 | 25.41 | 17.41 | 42.82 | 9.47 | 52.28 | 5.77 | 58.05 | 4.4 | 62.46 |

**Table 3.6.** Genera with the five largest loadings on the first (PCA1) and second (PCA2) Principal Component Axes (Euclidean PCoA) for lizard gut microbiota.

|  | Rank | Gut Genus | Loading | %Loading |
| --- | --- | --- | --- | --- |
| P  C  A  1 | 1 | *Dietzia* | 0.970388 | 94.17 |
|  | 2 | *Corynebacterium* | 0.169186 | 2.86 |
|  | 3 | unclassified Lachnospiraceae | -0.085009 | 0.72 |
|  | 4 | *Geodermatophilus* | -0.069416 | 0.48 |
|  | 5 | *Salmonella* | -0.055386 | 0.31 |
| P  C  A  2 | 1 | *Corynebacterium* | -0.946047 | 89.50 |
|  | 2 | *Dietzia* | 0.196495 | 3.86 |
|  | 3 | *Salmonella* | 0.149812 | 2.24 |
|  | 4 | unclassified Propionibacteriaceae | -0.141657 | 2.01 |
|  | 5 | unclassified Lachnospiraceae | 0.074217 | 0.55 |

**Table 3.7** Positions of the hybrid gut microbiota from SBluG along the first two genus PCoA axes. Column 4-5 show p-values for Mann-Whitney U tests assessing whether the positions of hybrid animals along the first PCoA axis is significantly different than the positions of *A. inornatus* (ino) and *A. marmoratus* (marm) respectively. Column 9-10 show p-values for Mann-Whitney U tests assessing whether the position of the hybrid animals along the second PCoA axis is significantly different than the positions of *A. inornatus* (ino) and *A. marmoratus* (marm) respectively. Yellow is used for tests significantly different at *p*>0.05.

|  | **PCoA axis 1**  **mean** | | | **p-value** | | **PCoA axis 2**  **mean** | | | **p-value** | |
| --- | --- | --- | --- | --- | --- | --- | --- | --- | --- | --- |
|  | **neo** | **ino** | **marm** | **ino** | **marm** | **neo** | **ino** | **marm** | **ino** | **marm** |
| **euclidean** | -832 | 140 | 2176 | 0.1421 | 0.0111 | 971 | -1859 | -144 | 0.0023 | 0.3498 |
| **jaccard** | -0.026 | -0.124 | 0.007 | 0.3331 | 0.8318 | 0.066 | 0.009 | -0.098 | 0.3331 | 0.01 |
| **bray** | -0.119 | 0.112 | 0.191 | 0.0219 | 0.0073 | -0.109 | -0.074 | 0.104 | 0.7314 | 0.0123 |

**Table 3.8** Positions of the hybrid gut microbiota from SNW along the first two genus PCoA axes. Column 4-5 show p-values for Mann-Whitney U tests assessing whether the positions of hybrid animals along the first PCoA axis is significantly different than the positions of *A. inornatus* (ino) and *A. marmoratus* (marm) respectively. Column 9-10 show p-values for Mann-Whitney U tests assessing whether the position of the hybrid animals along the second PCoA axis is significantly different than the positions of *A. inornatus* (ino) and *A. marmoratus* (marm) respectively. Yellow is used for tests significantly different at *p*>0.05.

|  | **PCoA axis 1**  **mean** | | | **p-value** | | **PCoA axis 2**  **mean** | | | **p-value** | |
| --- | --- | --- | --- | --- | --- | --- | --- | --- | --- | --- |
|  | **neo** | **ino** | **marm** | **ino** | **marm** | **neo** | **ino** | **marm** | **ino** | **marm** |
| **euclidean** | -1417 | 140 | 2176 | 0.151 | 0.012 | 907 | -1859 | -144 | 0.0106 | 0.4463 |
| **jaccard** | 0.157 | -0.124 | 0.007 | 0.0106 | 0.0856 | 0.011 | 0.009 | -0.098 | 0.9224 | 0.0856 |
| **bray** | -0.172 | 0.112 | 0.191 | 0.0055 | 0.0072 | 0.106 | -0.074 | 0.104 | 0.1102 | 0.9845 |

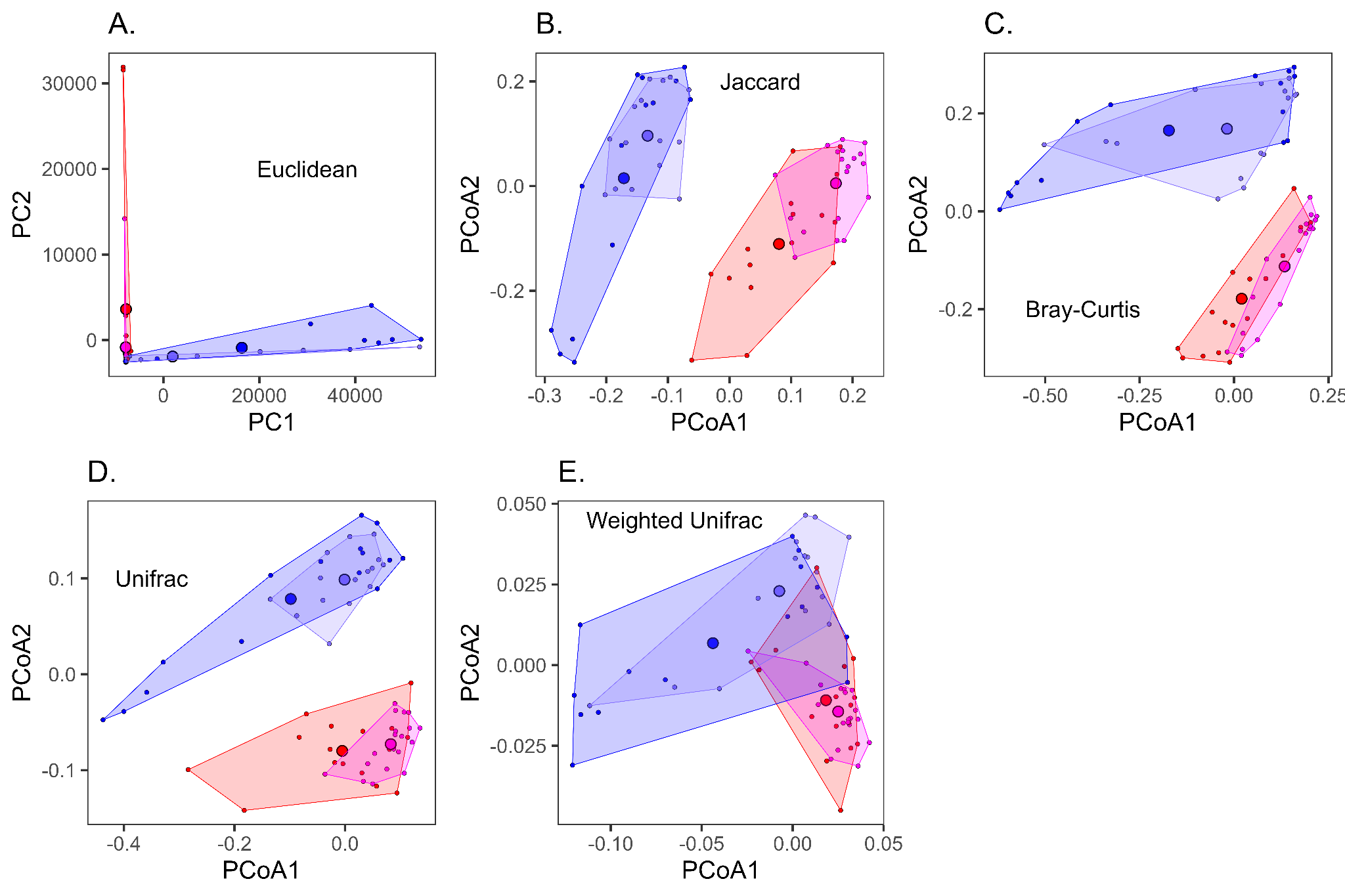

**Figure 3.3.** Two-dimensional Principal Coordinates Analysis (PCoA) of skin microbiota for populations of *A. inornatus* from SBluG (red), *A. neomexicanus* from SBluG (magenta), *A. neomexicanus* from SNW (purple), and *A. marmoratus* from SNW (blue) and based on amplicon sequence variants (ASVs) in combination with (A) euclidean dissimilarity, (B) Jaccard dissimilarity, (C) Bray-Curtis dissimilarity, (D) UniFrac dissimilarity, and (E) weighted UniFrac dissimilarity.

**Table 3.9.** Percent variation explained by the first five Principal Coordinate Axes, both individually (%) and cumulatively (Cum%) for PCoA analyses of skin ASVs using Euclidean, Jaccard, Bray-Curtis, UniFrac, and weighted UniFrac distances.

|  | PCoA1 | | PCoA2 | | PCoA3 | | PCoA4 | | PCoA5 | |
| --- | --- | --- | --- | --- | --- | --- | --- | --- | --- | --- |
|  | % | Cum% | % | Cum% | % | Cum% | % | Cum% | % | Cum% |
| Euclidean  (PCA) | 73.99 | 73.99 | 8.78 | 82.78 | 3.55 | 86.33 | 2.31 | 88.64 | 1.49 | 90.13 |
| Jaccard | 7.5 | 7.5 | 6.65 | 14.15 | 3.77 | 17.92 | 3.14 | 21.06 | 2.56 | 23.62 |
| Bray-Curtis | 18.62 | 18.62 | 12.52 | 31.14 | 7.29 | 38.43 | 4.56 | 42.99 | 3.62 | 46.61 |
| UniFrac | 11.57 | 11.57 | 6.05 | 17.62 | 3.59 | 21.22 | 2.86 | 24.07 | 2.43 | 26.51 |
| w-UniFrac | 45.57 | 45.57 | 11.12 | 56.69 | 5.21 | 61.9 | 4.67 | 66.56 | 4.17 | 70.74 |

**Table 3.10.** ASVs with the five largest loadings on the first (PCA1) and second (PCA2) Principal Component Axes (Euclidean PCoA) for lizard skin microbiota.

|  | Rank | Skin ASV | Loading | %Loading |
| --- | --- | --- | --- | --- |
| P  C  A  1 | 1 | *Fodinibacter luteus* | 0.995745 | 99.15 |
|  | 2 | *Arthrobacter agilis* | 0.056460 | 0.32 |
|  | 3 | *Nocardioides mesophilus* | -0.020697 | 0.04 |
|  | 4 | *Kocuria polaris* | 0.020384 | 0.04 |
|  | 5 | *Dietzia maris* | -0.019132 | 0.04 |
| P  C  A  2 | 1 | *Dietzia maris* | 0.986550 | 97.33 |
|  | 2 | *Geodermatophilus africanus-obscurus* | -0.074128 | 0.55 |
|  | 3 | *Corynebacterium testudinoris* | 0.058295 | 0.34 |
|  | 4 | *Geodermatophilus africanus-obscurus-siccatus* | -0.043706 | 0.19 |
|  | 5 | *Aeromicrobium massiliense* | 0.042662 | 0.18 |

**Table 3.11** Positions of the hybrid skin microbiota from SNW along the first two ASV PCoA axes. Column 4-5 show p-values for Mann-Whitney U tests assessing whether the positions of hybrid animals along the first PCoA axis is significantly different than the positions of *A. inornatus* (ino) and *A. marmoratus* (marm) respectively. Column 9-10 show p-values for Mann-Whitney U tests assessing whether the position of the hybrid animals along the second PCoA axis is significantly different than the positions of *A. inornatus* (ino) and *A. marmoratus* (marm) respectively. Yellow is used for tests significantly different at *p*>0.05.

|  | PCoA axis 1  mean | | | p-value | | PCoA axis 2  mean | | | p-value | |
| --- | --- | --- | --- | --- | --- | --- | --- | --- | --- | --- |
|  | neo | ino | marm | ino | marm | neo | ino | marm | ino | marm |
| euclidean | -1922 | 7799 | -16353 | 0.0023 | 0.5125 | -1914 | 3615 | -885 | 4e-04 | 0.3245 |
| jaccard | -0.133 | 0.08 | -0.172 | 0 | 0.2169 | 0.096 | -0.111 | 0.015 | 0 | 0.8063 |
| bray | -0.019 | 0.02 | -0.173 | 0.8916 | 0.3453 | 0.168 | -0.179 | 0.165 | 0 | 0.8063 |
| unifrac | -0.001 | -0.005 | -0.098 | 0.5986 | 0.5125 | 0.099 | -0.08 | 0.079 | 0 | 0.9674 |
| wunifrac | -0.007 | 0.018 | -0.044 | 0.0048 | 0.0742 | 0.023 | -0.011 | 0.007 | 1e-04 | 0.0453 |

**Table 3.12** Positions of the hybrid skin microbiota from SBluG along the first two ASV PCoA axes. Column 4-5 show p-values for Mann-Whitney U tests assessing whether the positions of hybrid animals along the first PCoA axis is significantly different than the positions of *A. inornatus* (ino) and *A. marmoratus* (marm) respectively. Column 9-10 show p-values for Mann-Whitney U tests assessing whether the position of the hybrid animals along the second PCoA axis is significantly different than the positions of *A. inornatus* (ino) and *A. marmoratus* (marm) respectively. Yellow is used for tests significantly different at *p*>0.05.

|  | PCoA axis 1  mean | | | p-value | | PCoA axis 2  mean | | | p-value | |
| --- | --- | --- | --- | --- | --- | --- | --- | --- | --- | --- |
|  | neo | ino | marm | ino | marm | neo | ino | marm | ino | marm |
| euclidean | 7860 | 7799 | -16353 | 0.2569 | 8e-04 | -835 | 3615 | -885 | 0.0441 | 0.5836 |
| jaccard | 0.173 | 0.08 | -0.172 | 0 | 0 | 0.005 | -0.111 | 0.015 | 0.0026 | 0.2561 |
| bray | 0.134 | 0.02 | -0.173 | 0.0014 | 0.0039 | -0.113 | -0.179 | 0.165 | 0.0883 | 0 |
| unifrac | 0.083 | -0.005 | -0.098 | 0.0037 | 1e-04 | -0.073 | -0.08 | 0.079 | 0.4611 | 0 |
| wunifrac | 0.025 | 0.018 | -0.044 | 0.3331 | 0 | -0.014 | -0.011 | 0.007 | 0.5669 | 8e-04 |

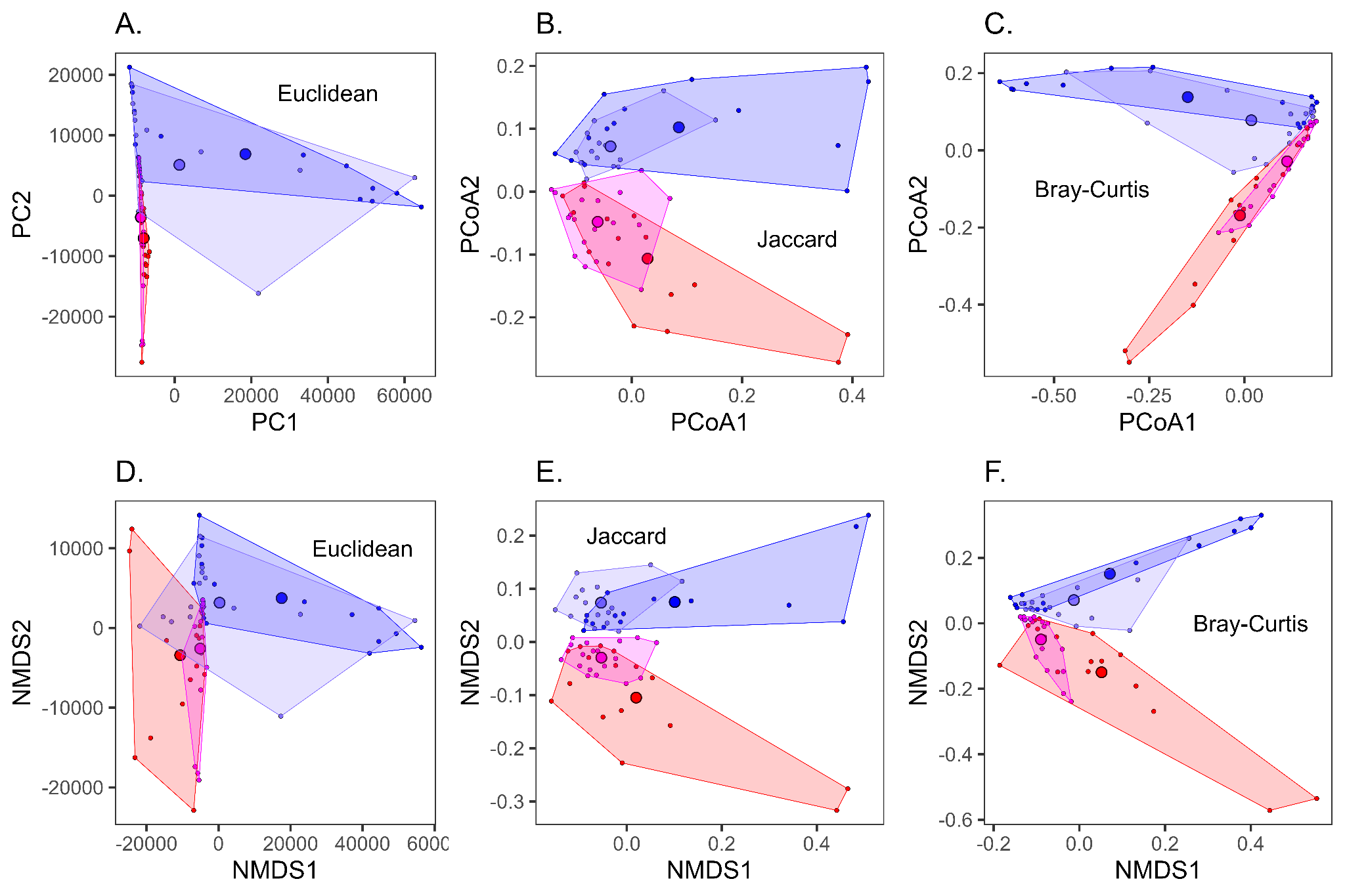
**Figure 3.4.** Two-dimensional Principal Coordinates Analysis (PCoA, A-C) or Non-metric MultiDimensional Scaling (NMDS, D-F) of skin microbiota for populations of *A. inornatus* from SBluG (red), *A. neomexicanus* from SBluG (magenta), *A. neomexicanus* from SNW (purple) and *A. marmoratus* from SNW (blue) based on microbial genera in combination with (A) euclidean dissimilarity, (B) Jaccard dissimilarity, and (C) Bray-Curtis dissimilarity.

**Table 3.13.** Percent variation explained by the first five Principal Coordinate Axes, both individually (%) and cumulatively (Cum%) for PCoA analyses of skin genera using Euclidean, Jaccard, and Bray-Curtis distances.

|  | PCoA1 | | PCoA2 | | PCoA3 | | PCoA4 | | PCoA5 | |
| --- | --- | --- | --- | --- | --- | --- | --- | --- | --- | --- |
|  | % | Cum% | % | Cum% | % | Cum% | % | Cum% | % | Cum% |
| Euclidean  (PCA) | 56.58 | 56.58 | 14.53 | 71.11 | 8.38 | 79.49 | 5.93 | 85.42 | 5.23 | 90.65 |
| Jaccard | 14.79 | 14.79 | 7.62 | 22.41 | 5.63 | 28.04 | 3.87 | 31.91 | 3.36 | 35.27 |
| Bray-Curtis | 33.98 | 33.98 | 17.64 | 51.62 | 11.08 | 62.69 | 7.23 | 69.93 | 4.48 | 74.41 |

**Table 3.14.** Genera with the five largest loadings on the first (PCA1) and second (PCA2) Principal Component Axes (Euclidean PCoA) for lizard skin microbiota.

|  | Rank | Skin Genus | Loading | %Loading |
| --- | --- | --- | --- | --- |
| P  C  A  1 | 1 | *Fodinibacter* | 0.979675 | 95.98 |
|  | 2 | *Geodermatophilus* | -0.120809 | 1.46 |
|  | 3 | *Streptomyces* | -0.066442 | 0.44 |
|  | 4 | *Arthrobacter* | 0.054813 | 0.30 |
|  | 5 | *Rubrobacter* | -0.046422 | 0.22 |
| P  C  A  2 | 1 | *Geodermatophilus* | 0.705442 | 49.76 |
|  | 2 | *Nocardioides* | -0.623731 | 38.90 |
|  | 3 | *Dietzia* | -0.143556 | 2.06 |
|  | 4 | unclassified Lachnospiraceae | -0.117034 | 1.37 |
|  | 5 | *Acinetobacter* | -0.097352 | 0.95 |

**Table 3.15** Positions of the hybrid skin microbiota from SBluG along the first two genus PCoA axes. Column 4-5 show p-values for Mann-Whitney U tests assessing whether the positions of hybrid animals along the first PCoA axis is significantly different than the positions of *A. inornatus* (ino) and *A. marmoratus* (marm) respectively. Column 9-10 show p-values for Mann-Whitney U tests assessing whether the position of the hybrid animals along the second PCoA axis is significantly different than the positions of *A. inornatus* (ino) and *A. marmoratus* (marm) respectively. Yellow is used for tests significantly different at *p*>0.05.

|  | **PCoA axis 1**  **mean** | | | **p-value** | | **PCoA axis 2**  **mean** | | | **p-value** | |
| --- | --- | --- | --- | --- | --- | --- | --- | --- | --- | --- |
|  | **neo** | **ino** | **marm** | **ino** | **marm** | **neo** | **ino** | **marm** | **ino** | **marm** |
| **euclidean** | 1262 | -8056 | 18469 | 0.2814 | 0.3453 | -5108 | 7025 | -6899 | 8.00E-04 | 0.8063 |
| **jaccard** | -0.038 | 0.029 | 0.085 | 0.2641 | 0.3046 | 0.072 | -0.107 | 0.102 | 0 | 0.1485 |
| **bray** | 0.018 | -0.011 | -0.149 | 0.2995 | 0.4864 | 0.078 | -0.169 | 0.138 | 0 | 0.0164 |

**Table 3.16** Positions of the hybrid skin microbiota from SBluG along the first two genus PCoA axes. Column 4-5 show p-values for Mann-Whitney U tests assessing whether the positions of hybrid animals along the first PCoA axis is significantly different than the positions of *A. inornatus* (ino) and *A. marmoratus* (marm) respectively. Column 9-10 show p-values for Mann-Whitney U tests assessing whether the position of the hybrid animals along the second PCoA axis is significantly different than the positions of *A. inornatus* (ino) and *A. marmoratus* (marm) respectively. Yellow is used for tests significantly different at *p*>0.05.

|  | **PCoA axis 1**  **mean** | | | **p-value** | | **PCoA axis 2**  **mean** | | | **p-value** | |
| --- | --- | --- | --- | --- | --- | --- | --- | --- | --- | --- |
|  | **neo** | **ino** | **marm** | **ino** | **marm** | **neo** | **ino** | **marm** | **ino** | **marm** |
| **euclidean** | -8793 | -8056 | 18469 | 0.0023 | 0.3024 | 3563 | 7025 | -6899 | 0.0656 | 0.0187 |
| **jaccard** | -0.062 | 0.029 | 0.085 | 0.0479 | 0.0557 | -0.048 | -0.107 | 0.102 | 0.0948 | 0 |
| **bray** | 0.112 | -0.011 | -0.149 | 0.0037 | 0.1675 | -0.029 | -0.169 | 0.138 | 0.015 | 0 |
