## Additional file 4 for "Transgressive Hybrids as Hopeful Holobionts"

**Additional file 4: PERMANOVA**

PERMANOVA was performed on Euclidean, Jaccard, Bray-Curtis, unweighted UniFrac, and weighted Unifrac distances for microbial ASVs, and Euclidean, Jaccard, and Bray-Curtis distances for microbial genera. All distances were calculated using the vegdist function from the vegan package. PERMANOVA was performed simultaneously on all four lizard populations using the PERMANOVA function (10,000 permutations) from the PERMANOVA package (version 0.2.0)^1^. When this indicated a significant difference between groups, post-hoc testing was performed based on pairwise multilevel comparisons using the pairwise.adonis function from the pairwiseAdonis package (version 0.4.1)^2^ with a Benjamini-Hochberg correction for multiple comparisons.

**Table 4.1.** Significance of PERMANOVA results for population comparisons of ASV gut microbiota. Significance was first assessed using a global test on all four populations followed by post-hoc pairwise comparisons using a Benjamini-Hochberg correction. Significance levels are indicated as follows: p-value ≤ 0.001 (***), p-value ≤ 0.01 (**), p-value ≤ 0.05 (*), p-value ≤ 0.1 (.). Dark grey cells indicate significant differences p-value ≤ 0.05, light grey cells indicate marginally significant differences p-value ≤ 0.1, and white squares indicate non-significant differences.

|  | Euclidean | Jaccard | Bray-Curtis | UniFrac | w-UniFrac |
| --- | --- | --- | --- | --- | --- |
| SBluG_ino vs SNW_marm | - | * | - | - | . |
| SBluG_ino vs SNW_neo | - | ** | ** | . | . |
| SBluG_ino vs SBluG_neo | - | - | ** | - | . |
| SNW_marm vs SNW_neo | * | ** | ** | . | ** |
| SNW_marm vs SBluG_neo | * | ** | ** | - | ** |
| SNW_neo vs SBluG_neo | - | ** | - | . | . |

**Table 4.2.** Significance of PERMANOVA results for population comparisons of gut microbiota based on microbial genus. Significance was first assessed using a global test on all four populations followed by post-hoc pairwise comparisons using a Benjamini-Hochberg correction. Significance levels are indicated as follows: p-value ≤ 0.001 (***), p-value ≤ 0.01 (**), p-value ≤ 0.05 (*), p-value ≤ 0.1 (.). Dark grey cells indicate significant differences p-value ≤ 0.05, light grey cells indicate marginally significant differences p-value ≤ 0.1, and white squares indicate non-significant differences.

|  | Euclidean | Jaccard | Bray-Curtis |
| --- | --- | --- | --- |
| SBluG_ino vs SNW_marm | - | - | - |
| SBluG_ino vs SNW_neo | - | * | * |
| SBluG_ino vs SBluG_neo | - | - | - |
| SNW_marm vs SNW_neo | - | . | * |
| SNW_marm vs SBluG_neo | - | - | * |
| SNW_neo vs SBluG_neo | - | . | - |

**Table 4.3.** Significance of PERMANOVA results for population comparisons of ASV skin microbiota. Significance was first assessed using a global test on all four populations followed by post-hoc pairwise comparisons using a Benjamini-Hochberg correction. Significance levels are indicated as follows: p-value ≤ 0.001 (***), p-value ≤ 0.01 (**), p-value ≤ 0.05 (*), p-value ≤ 0.1 (.). Dark grey cells indicate significant differences p-value ≤ 0.05, light grey cells indicate marginally significant differences p-value ≤ 0.1, and white squares indicate non-significant differences.

|  | Euclidean | Jaccard | Bray-Curtis | UniFrac | w-UniFrac |
| --- | --- | --- | --- | --- | --- |
| SBluG_ino vs SNW_marm | - | ** | * | - | . |
| SBluG_ino vs SNW_neo | - | ** | ** | ** | . |
| SBluG_ino vs SBluG_neo | - | ** | * | ** | * |
| SNW_marm vs SNW_neo | - | - | - | - | - |
| SNW_marm vs SBluG_neo | * | ** | ** | ** | ** |
| SNW_neo vs SBluG_neo | - | ** | ** | ** | ** |

**Table 4.4.** Significance of PERMANOVA results for population comparisons of skin microbiota based on microbial genus. Significance was first assessed using a global test on all four populations followed by post-hoc pairwise comparisons using a Benjamini-Hochberg correction. Significance levels are indicated as follows: p-value ≤ 0.001 (***), p-value ≤ 0.01 (**), p-value ≤ 0.05 (*), p-value ≤ 0.1 (.). Dark grey cells indicate significant differences p-value ≤ 0.05, light grey cells indicate marginally significant differences p-value ≤ 0.1, and white squares indicate non-significant differences.

|  | Euclidean | Jaccard | Bray-Curtis |
| --- | --- | --- | --- |
| SBluG_ino vs SNW_marm | . | - | - |
| SBluG_ino vs SNW_neo | - | * | - |
| SBluG_ino vs SBluG_neo | * | * | * |
| SNW_marm vs SNW_neo | - | - | - |
| SNW_marm vs SBluG_neo | * | * | * |
| SNW_neo vs SBluG_neo | * | ** | . |

**Table 4.5.** p-values from ANOVA tests for differences in dispersion between groups.

|  | Euclidean | Jaccard | Bray-Curtis | UniFrac | w-UniFrac |
| --- | --- | --- | --- | --- | --- |
| Gut ASV | 0.1609 | 0.9556 | 0.01227* | 0.1144 | 0.04956* |
| Skin ASV | 0.006** | 0.0006162*** | 0.00486** | 0.0127* | 0.007493** |
| Gut Genus | 0.5613 | 0.1171 | 0.7002 | - | - |
| Skin Genus | 0.001142 | 0.02097 | 0.008787 | - | - |

**Table 4.6.** PERMANOVA results for asexual vs. sexual comparison for gut and skin microbiota. Significance was assessed using a Mann-Whitney test. Significance levels are indicated as follows: p-value ≤ 0.001 (***), p-value ≤ 0.01 (**), p-value ≤ 0.05 (*), p-value ≤ 0.1 (.). Dark grey cells indicate significant differences p-value ≤ 0.05, light grey cells indicate marginally significant differences p-value ≤ 0.1, and white squares indicate non-significant differences or comparisons that were not considered.

|  | Euclidean | Jaccard | Bray-Curtis | UniFrac | w-UniFrac |
| --- | --- | --- | --- | --- | --- |
| PERMANOVA  (gut ASV) | *** | *** | *** | * | *** |
| PERMANOVA  (gut Genus) | ** | * | *** | NA | NA |
| PERMANOVA  (skin ASV) | - | ** | * | ** | . |
| PERMANOVA  (skin Genus) | . | ** | * | NA | NA |

**Table 4.7.** p-values for dispersion differences across asexual vs. sexual species for gut and skin microbiota.

|  | Euclidean | Jaccard | Bray-Curtis | UniFrac | w-UniFrac |
| --- | --- | --- | --- | --- | --- |
| Gut ASV | 0.01414* | 0.3784 | 0.01082* | 0.196 | 0.4258 |
| Skin ASV | 0.02824* | 0.006152** | 0.01972* | 0.001426** | 0.007117** |
| Gut Genus | 0.1433 | 0.06367 | 0.8271 | - | - |
| Skin Genus | 0.01895 | 0.002665 | 0.002473 | - | - |

**Table 4.8.** PERMANOVA results for species comparisons for gut and skin microbiota. Significance was first assessed using a global test on all three species followed by post-hoc pairwise comparisons using a Benjamini-Hochberg correction. Significance levels are indicated as follows: p-value ≤ 0.001 (***), p-value ≤ 0.01 (**), p-value ≤ 0.05 (*), p-value ≤ 0.1 (.). Dark grey cells indicate significant differences p-value ≤ 0.05, light grey cells indicate marginally significant differences p-value ≤ 0.1, and white squares indicate non-significant differences.

|  | Euclidean | | | Jaccard | | | Bray-Curtis | | | UniFrac | | | w-UniFrac | | |
| --- | --- | --- | --- | --- | --- | --- | --- | --- | --- | --- | --- | --- | --- | --- | --- |
|  | I  N | M  N | M  I | I  N | M  N | M  I | I  N | M  N | M  I | I  N | M  N | M  I | I  N | M  N | M  I |
| PERMANOVA (gut ASV) | . | ** | - | ** | ** | * | ** | ** | - | - | - | - | . | ** | . |
| PERMANOVA (gut Genus) | - | . | - | . | - | - | ** | ** | - | NA | NA | NA | NA | NA | NA |
| PERMANOVA (skin ASV) | - | * | . | ** | ** | ** | * | * | * | ** | * | - | . | * | . |
| PERMANOVA (skin Genus) | - | * | . | ** | ** | - | * | * | - | NA | NA | NA | NA | NA | NA |

**Table 4.9.** p-values for dispersion differences across species for gut and skin microbiota. Significance levels are indicated as follows: p-value ≤ 0.001 (***), p-value ≤ 0.01 (**), p-value ≤ 0.05 (*).

|  | Euclidean | Jaccard | Bray-Curtis | UniFrac | w-UniFrac |
| --- | --- | --- | --- | --- | --- |
| Gut ASV | 0.1097 | 0.0005672*** | 0.0008962*** | 0.4598 | 0.02293* |
| Skin ASV | 0.007267** | 0.008357** | 0.01262* | 0.2485 | 0.06513 |
| Gut Genus | 0.4799 | 0.5653 | 0.3219 | - | - |
| Skin Genus | 0.007322 | 0.1399 | 0.03978 | - | - |

**Table 4.10.** PERMANOVA statistics for gut ASV analyses (see Table 4.1). Significance levels are indicated as follows: p-value ≤ 0.001 (***), p-value ≤ 0.01 (**), p-value ≤ 0.05 (*).

|  | Sums Of Sqs | F.Model | R^2^ | p.value | p.adjusted |
| --- | --- | --- | --- | --- | --- |
| Euclidean | | | | | |
| SBluG_ino vs SNW_marm | 0.04141904 | 1.28522195 | 0.04388637 | 0.264 | 0.3168 |
| SBluG_ino vs SNW_neo | 0.05453879 | 2.24146632 | 0.07411897 | 0.115 | 0.1725 |
| SBluG_ino vs SBluG_neo | 0.0867081 | 3.40580687 | 0.10844513 | 0.062 | 0.124 |
| SNW_marm vs SNW_neo | 0.1523795 | 5.97535603 | 0.17587324 | 0.009** | 0.027* |
| SNW_marm vs SBluG_neo | 0.20857182 | 7.8326594 | 0.21858995 | 0.005** | 0.027* |
| SNW_neo vs SBluG_neo | 0.00724921 | 0.38697395 | 0.0136321 | 0.622 | 0.622 |
| Jaccard | | | | | |
| SBluG_ino vs SNW_marm | 0.00083556 | 2.06586504 | 0.06871131 | 0.034* | 0.0408* |
| SBluG_ino vs SNW_neo | 0.00146353 | 4.24254923 | 0.13158231 | 0.002** | 0.004** |
| SBluG_ino vs SBluG_neo | 0.00049153 | 1.25863254 | 0.04301748 | 0.198 | 0.198 |
| SNW_marm vs SNW_neo | 0.00075464 | 2.23674165 | 0.0739743 | 0.006** | 0.009** |
| SNW_marm vs SBluG_neo | 0.00105412 | 2.75263237 | 0.08950884 | 0.001** | 0.004** |
| SNW_neo vs SBluG_neo | 0.00109329 | 3.3800086 | 0.10771216 | 0.002** | 0.004** |
| Bray-Curtis | | | | | |
| SBluG_ino vs SNW_marm | 0.00747972 | 2.51738902 | 0.08249031 | 0.107 | 0.1284 |
| SBluG_ino vs SNW_neo | 0.01328212 | 6.57450935 | 0.19015481 | 0.005** | 0.0075** |
| SBluG_ino vs SBluG_neo | 0.01236017 | 5.95667677 | 0.1754199 | 0.005** | 0.0075** |
| SNW_marm vs SNW_neo | 0.02693784 | 13.9600423 | 0.33269848 | 0.001** | 0.003** |
| SNW_marm vs SBluG_neo | 0.02927207 | 14.7510554 | 0.34504541 | 0.001** | 0.003** |
| SNW_neo vs SBluG_neo | 0.00155189 | 1.50169345 | 0.05090194 | 0.248 | 0.248 |
| UniFrac | | | | | |
| SBluG_ino vs SNW_marm | 0.00443588 | 1.07261798 | 0.03689444 | 0.366 | 0.4392 |
| SBluG_ino vs SNW_neo | 0.01467193 | 4.71885133 | 0.14422424 | 0.015* | 0.072 |
| SBluG_ino vs SBluG_neo | 0.00178089 | 0.44090201 | 0.01550239 | 0.707 | 0.707 |
| SNW_marm vs SNW_neo | 0.01324868 | 4.15449697 | 0.12920423 | 0.024* | 0.072 |
| SNW_marm vs SBluG_neo | 0.00447777 | 1.08710689 | 0.03737418 | 0.326 | 0.4392 |
| SNW_neo vs SBluG_neo | 0.00916175 | 2.96244507 | 0.09567865 | 0.047* | 0.094 |
| weighted UniFrac | | | | | |
| SBluG_ino vs SNW_marm | 0.05032898 | 2.98483683 | 0.09633218 | 0.057 | 0.0684 |
| SBluG_ino vs SNW_neo | 0.05382376 | 3.31521042 | 0.10586582 | 0.036* | 0.066 |
| SBluG_ino vs SBluG_neo | 0.04648164 | 2.48511759 | 0.08151904 | 0.094 | 0.094 |
| SNW_marm vs SNW_neo | 0.08161288 | 7.56136063 | 0.21262855 | 0.003** | 0.009** |
| SNW_marm vs SBluG_neo | 0.15908131 | 11.9952594 | 0.29991703 | 0.001** | 0.006** |
| SNW_neo vs SBluG_neo | 0.04387976 | 3.47263725 | 0.1103383 | 0.044* | 0.066 |

**Table 4.11.** PERMANOVA statistics for gut genera analyses (see Table 4.2). Significance levels are indicated as follows: p-value ≤ 0.001 (***), p-value ≤ 0.01 (**), p-value ≤ 0.05 (*).

|  | Sums Of Sqs | F.Model | R^2^ | p.value | p.adjusted |
| --- | --- | --- | --- | --- | --- |
| Euclidean | | | | | |
| SNW_marm vs SNW_neo | 0.08650074 | 3.37177162 | 0.10747788 | 0.055 | 0.165 |
| SNW_marm vs SBluG_ino | 0.02126703 | 0.82295127 | 0.02855194 | 0.41 | 0.492 |
| SNW_marm vs SBluG_neo | 0.11328755 | 3.92644205 | 0.12298402 | 0.048* | 0.165 |
| SNW_neo vs SBluG_ino | 0.04313415 | 2.0661626 | 0.06872053 | 0.135 | 0.2025 |
| SNW_neo vs SBluG_neo | 0.00899793 | 0.3766946 | 0.01327479 | 0.611 | 0.611 |
| SBluG_ino vs SBluG_neo | 0.06176811 | 2.56570324 | 0.08394059 | 0.121 | 0.2025 |
| Jaccard | | | | | |
| SNW_marm vs SNW_neo | 0.01166802 | 4.05093834 | 0.12639063 | 0.027* | 0.081 |
| SNW_marm vs SBluG_ino | 0.00454871 | 1.15891465 | 0.03974478 | 0.325 | 0.381 |
| SNW_marm vs SBluG_neo | 0.00373469 | 0.92686771 | 0.03204176 | 0.381 | 0.381 |
| SNW_neo vs SBluG_ino | 0.02391125 | 7.85566372 | 0.21909129 | 0.006** | 0.036* |
| SNW_neo vs SBluG_neo | 0.01045317 | 3.32034895 | 0.10601251 | 0.042* | 0.084 |
| SBluG_ino vs SBluG_neo | 0.00421906 | 1.00624835 | 0.03469074 | 0.335 | 0.381 |
| Bray-Curtis | | | | | |
| SNW_marm vs SNW_neo | 0.0453224 | 7.23244878 | 0.20527806 | 0.005** | 0.018* |
| SNW_marm vs SBluG_ino | 0.01229819 | 1.39776138 | 0.04754652 | 0.241 | 0.241 |
| SNW_marm vs SBluG_neo | 0.04587498 | 5.38331648 | 0.16125769 | 0.013* | 0.026* |
| SNW_neo vs SBluG_ino | 0.03537377 | 5.90934651 | 0.17426896 | 0.006** | 0.018* |
| SNW_neo vs SBluG_neo | 0.0106935 | 1.87300502 | 0.06269892 | 0.14 | 0.168 |
| SBluG_ino vs SBluG_neo | 0.0231077 | 2.80391439 | 0.09102461 | 0.069 | 0.1035 |

**Table 4.12.** PERMANOVA statistics for skin ASV analyses (see Table 4.3). Significance levels are indicated as follows: p-value ≤ 0.001 (***), p-value ≤ 0.01 (**), p-value ≤ 0.05 (*).

|  | Sums Of Sqs | F.Model | R^2^ | p.value | p.adjusted |
| --- | --- | --- | --- | --- | --- |
| Euclidean | | | | | |
| SNW_neo vs SNW_marm | 0.19664448 | 2.47755295 | 0.08129107 | 0.129 | 0.1548 |
| SNW_neo vs SBluG_ino | 0.01727338 | 0.44089928 | 0.0155023 | 0.547 | 0.547 |
| SNW_neo vs SBluG_neo | 0.09106028 | 2.96539912 | 0.09576492 | 0.08 | 0.12 |
| SNW_marm vs SBluG_ino | 0.25801384 | 3.96948385 | 0.12416478 | 0.054 | 0.112 |
| SNW_marm vs SBluG_neo | 0.52590484 | 9.3032194 | 0.24939454 | 0.004** | 0.024* |
| SBluG_ino vs SBluG_neo | 0.05621489 | 3.44106213 | 0.10944484 | 0.056 | 0.112 |
| Jaccard | | | | | |
| SNW_neo vs SNW_marm | 0.001637 | 1.57634263 | 0.05329742 | 0.192 | 0.192 |
| SNW_neo vs SBluG_ino | 0.01157207 | 14.1958632 | 0.33642784 | 0.001** | 0.002** |
| SNW_neo vs SBluG_neo | 0.01040793 | 19.0554172 | 0.40495693 | 0.001** | 0.002** |
| SNW_marm vs SBluG_ino | 0.0079291 | 6.23919494 | 0.18222376 | 0.002** | 0.003** |
| SNW_marm vs SBluG_neo | 0.0098815 | 9.86301271 | 0.26049202 | 0.001** | 0.002** |
| SBluG_ino vs SBluG_neo | 0.00383281 | 4.92291122 | 0.14952843 | 0.004** | 0.0048** |
| Bray-Curtis | | | | | |
| SNW_neo vs SNW_marm | 0.01006297 | 1.71371813 | 0.05767431 | 0.199 | 0.199 |
| SNW_neo vs SBluG_ino | 0.03292475 | 8.5074267 | 0.23303277 | 0.001** | 0.003** |
| SNW_neo vs SBluG_neo | 0.02596222 | 8.71914778 | 0.23745507 | 0.001** | 0.003** |
| SNW_marm vs SBluG_ino | 0.0281825 | 4.73490808 | 0.144644 | 0.01* | 0.015* |
| SNW_marm vs SBluG_neo | 0.03884995 | 7.67852289 | 0.21521415 | 0.004** | 0.008** |
| SBluG_ino vs SBluG_neo | 0.01276091 | 4.17341073 | 0.12971614 | 0.033* | 0.0396* |
| UniFrac | | | | | |
| SNW_neo vs SNW_marm | 0.00740221 | 3.01451328 | 0.09719686 | 0.098 | 0.1176 |
| SNW_neo vs SBluG_ino | 0.00988644 | 8.41207885 | 0.2310244 | 0.001** | 0.003** |
| SNW_neo vs SBluG_neo | 0.00839851 | 12.5375063 | 0.30928164 | 0.001** | 0.003** |
| SNW_marm vs SBluG_ino | 0.00548348 | 1.91027259 | 0.06386677 | 0.138 | 0.138 |
| SNW_marm vs SBluG_neo | 0.01328438 | 5.61677854 | 0.16708259 | 0.004** | 0.006** |
| SBluG_ino vs SBluG_neo | 0.00486036 | 4.48014294 | 0.13793483 | 0.003** | 0.006** |
| weighted UniFrac | | | | | |
| SNW_neo vs SNW_marm | 0.06913107 | 2.52970206 | 0.08286036 | 0.124 | 0.124 |
| SNW_neo vs SBluG_ino | 0.0468076 | 3.64717748 | 0.11524495 | 0.044* | 0.0624 |
| SNW_neo vs SBluG_neo | 0.07915039 | 7.39171736 | 0.20885444 | 0.002** | 0.006** |
| SNW_marm vs SBluG_ino | 0.07932571 | 3.31896621 | 0.10597304 | 0.052 | 0.0624 |
| SNW_marm vs SBluG_neo | 0.20088678 | 9.2256589 | 0.24783064 | 0.002** | 0.006** |
| SBluG_ino vs SBluG_neo | 0.03907796 | 5.36714363 | 0.16085116 | 0.007** | 0.014* |

**Table 4.13.** PERMANOVA statistics for skin genera analyses (see Table 4.4). Significance levels are indicated as follows: p-value ≤ 0.001 (***), p-value ≤ 0.01 (**), p-value ≤ 0.05 (*).

|  | Sums Of Sqs | F.Model | R^2^ | p.value | p.adjusted |
| --- | --- | --- | --- | --- | --- |
| Euclidean | | | | | |
| SBluG_ino vs SNW_marm | 0.19804334 | 4.61853848 | 0.14159244 | 0.045* | 0.0675 |
| SBluG_ino vs SBluG_neo | 0.07750271 | 5.22865463 | 0.15735379 | 0.024* | 0.048* |
| SBluG_ino vs SNW_neo | 0.03735685 | 1.35727822 | 0.04623311 | 0.253 | 0.253 |
| SNW_marm vs SBluG_neo | 0.47042381 | 12.7575351 | 0.31301047 | 0.004** | 0.018* |
| SNW_marm vs SNW_neo | 0.11721662 | 2.36443656 | 0.07786861 | 0.121 | 0.1452 |
| SBluG_neo vs SNW_neo | 0.14519356 | 6.74770935 | 0.19419149 | 0.006** | 0.018* |
| Jaccard | | | | | |
| SBluG_ino vs SNW_marm | 0.00714262 | 1.22945499 | 0.04206219 | 0.259 | 0.259 |
| SBluG_ino vs SBluG_neo | 0.01440114 | 5.04210065 | 0.15259625 | 0.007** | 0.018* |
| SBluG_ino vs SNW_neo | 0.01360729 | 4.38657314 | 0.13544419 | 0.009** | 0.018* |
| SNW_marm vs SBluG_neo | 0.0210731 | 5.32475111 | 0.15978367 | 0.018* | 0.027* |
| SNW_marm vs SNW_neo | 0.00972965 | 2.31469563 | 0.07635556 | 0.102 | 0.1224 |
| SBluG_neo vs SNW_neo | 0.00889538 | 7.11617762 | 0.2026467 | 0.001** | 0.006** |
| Bray-Curtis | | | | | |
| SBluG_ino vs SNW_marm | 0.04060713 | 1.7226889 | 0.05795872 | 0.176 | 0.176 |
| SBluG_ino vs SBluG_neo | 0.07663524 | 6.01116523 | 0.17674094 | 0.014* | 0.042* |
| SBluG_ino vs SNW_neo | 0.04287272 | 2.71577849 | 0.0884164 | 0.098 | 0.147 |
| SNW_marm vs SBluG_neo | 0.14241206 | 8.0536649 | 0.22337992 | 0.008** | 0.042* |
| SNW_marm vs SNW_neo | 0.04673829 | 2.25564328 | 0.07455281 | 0.124 | 0.1488 |
| SBluG_neo vs SNW_neo | 0.04501936 | 4.54857014 | 0.13974716 | 0.026* | 0.052 |

**Table 4.14.** Significance of PERMANOVA results for population comparisons of ASV gut and skin microbiota combined by population comparison. Yellow indicates that there are significant differences between EITHER skin microbiota, gut microbiota or both. Orange indicates that there are marginally significant differences (*p* < 0.1) between EITHER skin microbiota, gut microbiota or both. All else is as described for Tables 4.1 and 4.3.

|  |  | Euclidean | Jaccard | Bray-Curtis | UniFrac | w-UniFrac |
| --- | --- | --- | --- | --- | --- | --- |
| SBluG_ino  vs  SNW_neo | GUT | - | ** | ** | . | . |
|  | SKIN | - | ** | ** | ** | . |
| SBluG_ino  vs  SBluG_neo | GUT | - | - | ** | - | . |
|  | SKIN | - | ** | * | ** | * |
| SNW_marm  vs  SNW_neo | GUT | * | ** | ** | . | ** |
|  | SKIN | - | - | - | - | - |
| SNW_marm  vs  SBluG_neo | GUT | * | ** | ** | - | ** |
|  | SKIN | * | ** | ** | ** | ** |
