## Additional file 5 for "Transgressive Hybrids as Hopeful Holobionts"

**Additional file 5: Triangle Plots**

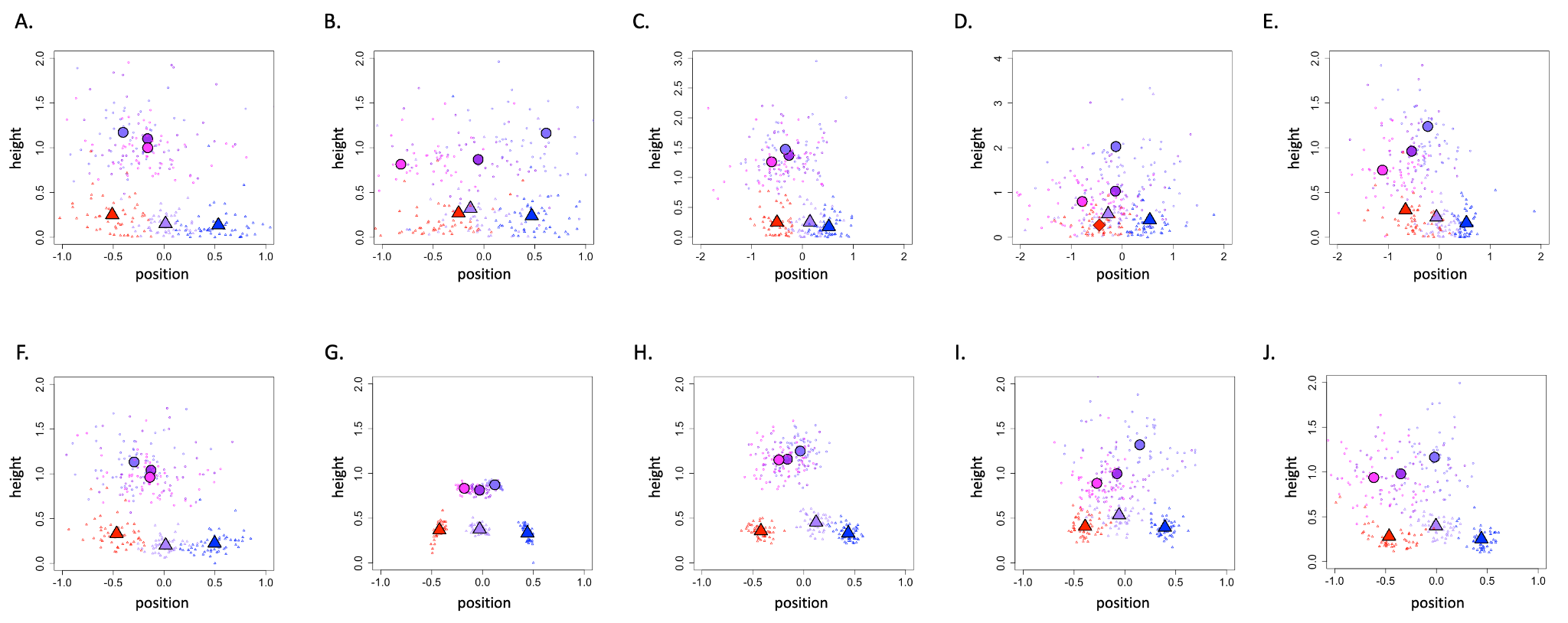

**Figure 5.1.** Position of the hybrid microbiota along the main parental axis of variation and height of the hybrid microbiota perpendicular to the parental axis of variation for gut ASVs (circles) of all hybrids (purple), hybrids at SBluG (magenta), and hybrids at SNW (blue-purple). Null models (triangles) are also shown for random subsamples of the *Aspidoscelis inornatus* population at SBluG (red), the *A. marmoratus* population at SNW (blue), and microbiota derived from ‘breeding’ (see Materials and Methods) *A. inornatus* at SBluG and *A. marmoratus* at SNW (light purple). In all panels, small open circles/triangles represent individual subsamples, while large filled circles/triangles represent means over 50 subsamples. The large diamond in panel (D) represents a median, because the mean was strongly influenced by one outlier. Panels (A-E) use the first two PCA/PCoA axes, panels (F-I) use the first 20 PCA/PCoA axes and panel (J) uses the first 15 PCoA axes due to convergence problems with 20 dimensions. Distance metrics for each panel are: Euclidean (A, F), Jaccard (B, G), Bray-Curtis (C, H), unweighted UniFrac (D, I), and weighted UniFrac (E, J).

**Table 5.1.** Position of the overall (i.e., both locations) hybrid microbiota along (median x) and perpendicular to (median y) the main parental axis of variation for gut ASVs. Median hybrid positions are calculated over 15 subsamples of 12 randomly selected *A. neomexicanus* individuals. Column 2-4 show p-values for Mann-Whitney U tests assessing whether the median position of the hybrid along the x-axis is significantly different than null models generated (1) using 12 randomly selected *A. inornatus* (ino) (2) using 12 randomly selected *A. marmoratus* (marm) and (3) summing the microbiota of 12 randomly selected *A. inornatus* and 12 randomly selected *A. marmoratus*. Column 6-8 show p-values for Mann-Whitney U tests assessing whether the median position of the hybrid along the y-axis is significantly greater than the same three null models. Yellow is used for tests significantly different at *p*>0.05.

|  | mean x | ino | marm | ino + marm | mean y | ino | marm | ino + marm |
| --- | --- | --- | --- | --- | --- | --- | --- | --- |
| euclidean_2 | 0.0149 | 0.0004 | 0.0003 | 1.0000 | 1.0454 | 0.0000 | 0.0000 | 0.0000 |
| euclidean_20 | 0.0046 | 0.0001 | 0.0000 | 1.0000 | 0.9877 | 0.0000 | 0.0000 | 0.0000 |
| jaccard_2 | 0.0525 | 0.0502 | 0.0675 | 0.4864 | 0.8752 | 0.0001 | 0.0000 | 0.0001 |
| jaccard_20 | -0.0067 | 0.0000 | 0.0000 | 0.3453 | 0.8223 | 0.0000 | 0.0000 | 0.0000 |
| bray_2 | -0.0442 | 0.0007 | 0.0000 | 0.1736 | 1.3412 | 0.0000 | 0.0000 | 0.0000 |
| bray_20 | -0.1739 | 0.0000 | 0.0000 | 0.0000 | 1.1648 | 0.0000 | 0.0000 | 0.0000 |
| unifrac_2 | -0.1055 | 0.3669 | 0.0075 | 0.4124 | 1.2398 | 0.0006 | 0.0000 | 0.0008 |
| unifrac_20 | -0.0269 | 0.0000 | 0.0002 | 0.1607 | 0.9408 | 0.0000 | 0.0000 | 0.0000 |
| wunifrac_2 | -0.5162 | 0.7130 | 0.0000 | 0.0003 | 1.0405 | 0.0000 | 0.0000 | 0.0000 |
| wunifrac_20 | -0.3390 | 0.3245 | 0.0000 | 0.0049 | 0.9621 | 0.0000 | 0.0000 | 0.0000 |

**Table 5.2.** Position of hybrid microbiota from SNW along (median x) and perpendicular to (median y) the main parental axis of variation for gut ASVs. Median hybrid positions are calculated over 15 subsamples of 12 randomly selected *A. neomexicanus* individuals. Column 2-4 show p-values for Mann-Whitney U tests assessing whether the median position of the hybrid along the x-axis is significantly different than null models generated (1) using 12 randomly selected *A. inornatus* (ino) (2) using 12 randomly selected *A. marmoratus* (marm) and (3) summing the microbiota of 12 randomly selected *A. inornatus* and 12 randomly selected *A. marmoratus*. Column 6-8 show p-values for Mann-Whitney U tests assessing whether the median position of the hybrid along the y-axis is significantly greater than the same three null models. Yellow is used for tests significantly different at *p*>0.05.

|  | mean x | ino | marm | ino + marm | mean y | ino | marm | ino +  marm |
| --- | --- | --- | --- | --- | --- | --- | --- | --- |
| euclidean_2 | -0.4484 | 0.6236 | 0.0000 | 0.0005 | 1.2246 | 0.0000 | 0.0000 | 0.0000 |
| euclidean_20 | -0.3862 | 0.8702 | 0.0000 | 0.0003 | 1.1688 | 0.0000 | 0.0000 | 0.0000 |
| jaccard_2 | 0.6186 | 0.0000 | 0.1607 | 0.0000 | 1.0827 | 0.0000 | 0.0000 | 0.0000 |
| jaccard_20 | 0.134 | 0.0000 | 0.0000 | 0.0000 | 0.8768 | 0.0000 | 0.0000 | 0.0000 |
| bray_2 | -0.4239 | 0.8063 | 0.0000 | 0.0006 | 1.396 | 0.0000 | 0.0000 | 0.0000 |
| bray_20 | -0.0547 | 0.0000 | 0.0000 | 0.0020 | 1.2357 | 0.0000 | 0.0000 | 0.0000 |
| unifrac_2 | 0.4114 | 0.0066 | 0.2328 | 0.0209 | 1.6918 | 0.0000 | 0.0000 | 0.0000 |
| unifrac_20 | 0.2816 | 0.0000 | 0.1160 | 0.0613 | 1.2652 | 0.0000 | 0.0000 | 0.0000 |
| wunifrac_2 | -0.181 | 0.0209 | 0.0000 | 0.0814 | 0.9764 | 0.0000 | 0.0000 | 0.0000 |
| wunifrac_20 | 0.0381 | 0.0000 | 0.0000 | 0.9674 | 1.1903 | 0.0000 | 0.0000 | 0.0000 |

**Table 5.3.** Position of hybrid microbiota from SBluG along (median x) and perpendicular to (median y) the main parental axis of variation for gut ASVs. Median hybrid positions are calculated over 15 subsamples of 12 randomly selected *A. neomexicanus* individuals. Column 2-4 show p-values for Mann-Whitney U tests assessing whether the median position of the hybrid along the x-axis is significantly different than null models generated (1) using 12 randomly selected *A. inornatus* (ino) (2) using 12 randomly selected *A. marmoratus* (marm) and (3) summing the microbiota of 12 randomly selected *A. inornatus* and 12 randomly selected *A. marmoratus*. Column 6-8 show p-values for Mann-Whitney U tests assessing whether the median position of the hybrid along the y-axis is significantly greater than the same three null models. Yellow is used for tests significantly different at *p*>0.05.

|  | mean x | ino | marm | ino + marm | mean y | ino | marm | ino + marm |
| --- | --- | --- | --- | --- | --- | --- | --- | --- |
| euclidean_2 | -0.1613 | 0.0027 | 0.0000 | 0.3892 | 0.9175 | 0.0000 | 0.0000 | 0.0000 |
| euclidean_20 | -0.1066 | 0.0010 | 0.0000 | 0.3046 | 0.8806 | 0.0000 | 0.0000 | 0.0000 |
| jaccard_2 | -0.6498 | 0.1485 | 0.0000 | 0.0057 | 0.8475 | 0.0001 | 0.0001 | 0.0001 |
| jaccard_20 | -0.1725 | 0.0000 | 0.0000 | 0.0000 | 0.8254 | 0.0000 | 0.0000 | 0.0000 |
| bray_2 | -0.4653 | 0.6529 | 0.0000 | 0.0017 | 1.2191 | 0.0000 | 0.0000 | 0.0000 |
| bray_20 | -0.2009 | 0.0002 | 0.0000 | 0.0000 | 1.2083 | 0.0000 | 0.0000 | 0.0000 |
| unifrac_2 | -0.6557 | 0.8381 | 0.0000 | 0.3245 | 0.8588 | 0.0007 | 0.0000 | 0.0010 |
| unifrac_20 | -0.2374 | 0.0086 | 0.0000 | 0.0006 | 0.896 | 0.0000 | 0.0000 | 0.0000 |
| wunifrac_2 | -0.9096 | 0.0408 | 0.0000 | 0.0000 | 0.7722 | 0.0002 | 0.0000 | 0.0000 |
| wunifrac_20 | -0.5897 | 0.4864 | 0.0000 | 0.0000 | 1.0508 | 0.0000 | 0.0000 | 0.0000 |

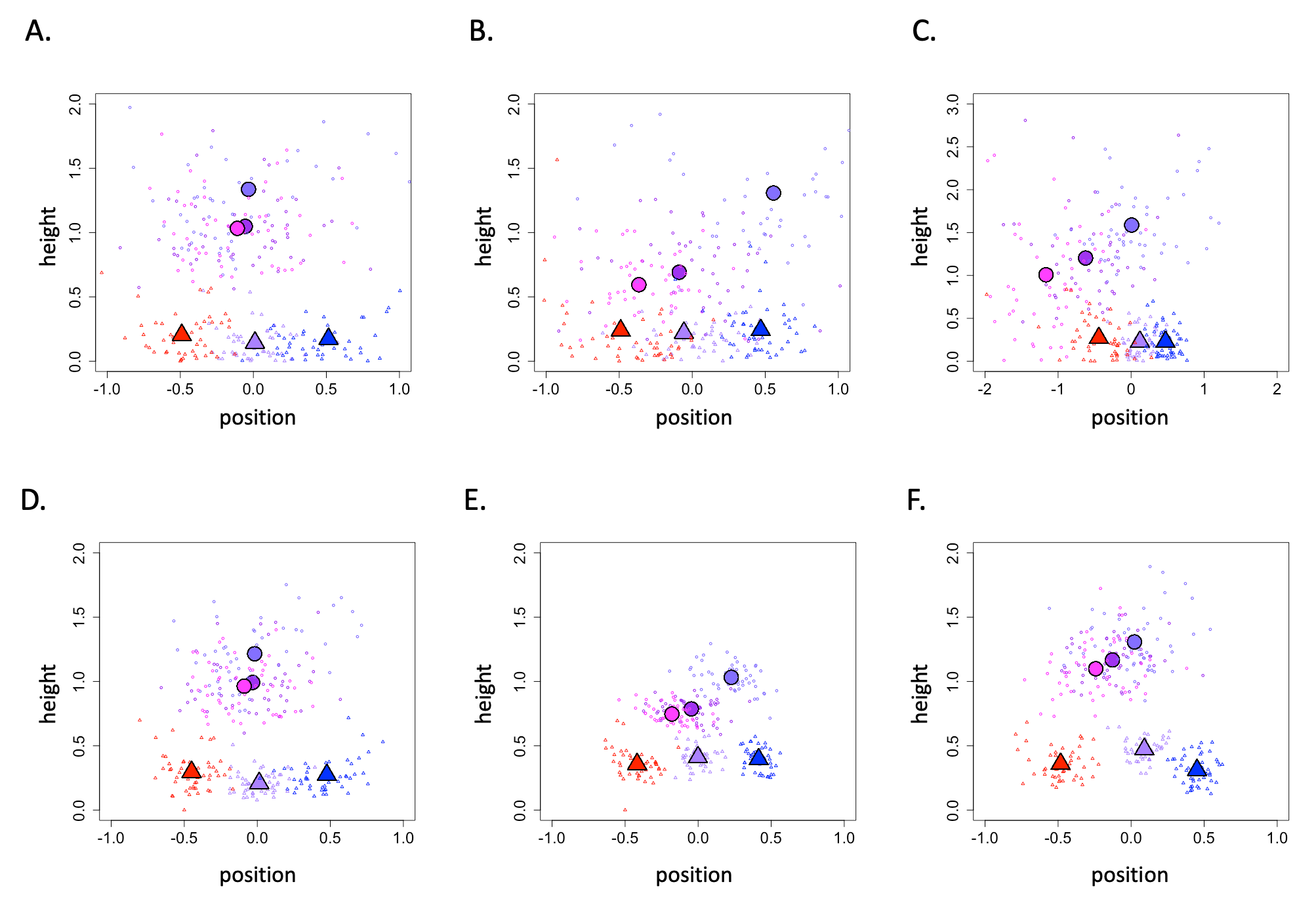

**Figure 5.2.** Position of the hybrid microbiota along the main parental axis of variation and height of the hybrid microbiota perpendicular to the parental axis of variation for gut genera (circles) of all hybrids (purple), hybrids at SBluG (magenta), and hybrids at SNW (blue-purple). Null models (triangles) are also shown for random subsamples of the *Aspidoscelis inornatus* population at SBluG (red), the *A. marmoratus* population at SNW (blue), and microbiota derived from ‘breeding’ (see Materials and Methods) *A. inornatus* at SBluG and *A. marmoratus* at `SNW (light purple). In all panels, small open circles/triangles represent individual subsamples, while large filled circles/triangles represent means over 50 subsamples. Panels (A-C) use the first two PCA/PCoA axes, while panels (D-F) use the first 20 PCA/PCoA axes. Distance metrics for each panel are: Euclidean (A, D), Jaccard (B, E), and Bray-Curtis (C, F).

**Table 5.4.** Position of the overall (i.e., both locations) hybrid microbiota along (median x) and perpendicular to (median y) the main parental axis of variation for gut genera. Median hybrid positions are calculated over 15 subsamples of 12 randomly selected *A. neomexicanus* individuals. Column 2-4 show p-values for Mann-Whitney U tests assessing whether the median position of the hybrid along the x-axis is significantly different than null models generated (1) using 12 randomly selected *A. inornatus* (ino) (2) using 12 randomly selected *A. marmoratus* (marm) and (3) summing the microbiota of 12 randomly selected *A. inornatus* and 12 randomly selected *A. marmoratus*. Column 6-8 show p-values for Mann-Whitney U tests assessing whether the median position of the hybrid along the y-axis is significantly greater than the same three null models. Yellow is used for tests significantly different at *p*>0.05.

|  | mean x | ino | marm | ino + marm | mean y | ino | marm | ino + marm |
| --- | --- | --- | --- | --- | --- | --- | --- | --- |
| euclidean_2 | 0.021 | 0.0020 | 0.0113 | 0.8063 | 1.0185 | 0.0000 | 0.0000 | 0.0000 |
| euclidean_20 | 0.0415 | 0.0001 | 0.0010 | 0.5393 | 0.9736 | 0.0000 | 0.0000 | 0.0000 |
| jaccard_2 | -0.1116 | 0.0164 | 0.0007 | 0.5949 | 0.6556 | 0.0001 | 0.0001 | 0.0001 |
| jaccard_20 | -0.0442 | 0.0000 | 0.0000 | 0.7437 | 0.7776 | 0.0000 | 0.0000 | 0.0000 |
| bray_2 | -0.5792 | 0.4124 | 0.0000 | 0.0001 | 1.1544 | 0.0000 | 0.0000 | 0.0000 |
| bray_15 | -0.1665 | 0.0001 | 0.0000 | 0.0023 | 1.2058 | 0.0000 | 0.0000 | 0.0000 |

**Table 5.5.** Position of hybrid microbiota from SNW along (median x) and perpendicular to (median y) the main parental axis of variation for gut genera. Median hybrid positions are calculated over 15 subsamples of 12 randomly selected *A. neomexicanus* individuals. Column 2-4 show p-values for Mann-Whitney U tests assessing whether the median position of the hybrid along the x-axis is significantly different than null models generated (1) using 12 randomly selected *A. inornatus* (ino) (2) using 12 randomly selected *A. marmoratus* (marm) and (3) summing the microbiota of 12 randomly selected *A. inornatus* and 12 randomly selected *A. marmoratus*. Column 6-8 show p-values for Mann-Whitney U tests assessing whether the median position of the hybrid along the y-axis is significantly greater than the same three null models. Yellow is used for tests significantly different at *p*>0.05.

|  | mean x | ino | marm | ino + marm | mean y | ino | marm | ino + marm |
| --- | --- | --- | --- | --- | --- | --- | --- | --- |
| euclidean_2 | 0.003 | 0.0023 | 0.0066 | 0.6529 | 1.2425 | 0.0000 | 0.0000 | 0.0000 |
| euclidean_20 | 0.0289 | 0.0000 | 0.0007 | 0.8381 | 1.188 | 0.0000 | 0.0000 | 0.0000 |
| jaccard_2 | 0.5884 | 0.0000 | 0.5949 | 0.0329 | 1.4931 | 0.0000 | 0.0000 | 0.0000 |
| jaccard_20 | 0.2108 | 0.0000 | 0.0014 | 0.0007 | 1.0875 | 0.0000 | 0.0000 | 0.0000 |
| bray_2 | -0.0393 | 0.0099 | 0.1160 | 0.6236 | 1.5826 | 0.0000 | 0.0000 | 0.0000 |
| bray_15 | 0.0366 | 0.0000 | 0.0000 | 0.2854 | 1.3485 | 0.0000 | 0.0000 | 0.0000 |

**Table 5.6.** Position of hybrid microbiota from SBluG along (median x) and perpendicular to (median y) the main parental axis of variation for gut genera. Median hybrid positions are calculated over 15 subsamples of 12 randomly selected *A. neomexicanus* individuals. Column 2-4 show p-values for Mann-Whitney U tests assessing whether the median position of the hybrid along the x-axis is significantly different than null models generated (1) using 12 randomly selected *A. inornatus* (ino) (2) using 12 randomly selected *A. marmoratus* (marm) and (3) summing the microbiota of 12 randomly selected *A. inornatus* and 12 randomly selected *A. marmoratus*. Column 6-8 show p-values for Mann-Whitney U tests assessing whether the median position of the hybrid along the y-axis is significantly greater than the same three null models. Yellow is used for tests significantly different at *p*>0.05.

|  | mean x | ino | marm | ino + marm | mean y | ino | marm | ino + marm |
| --- | --- | --- | --- | --- | --- | --- | --- | --- |
| euclidean_2 | -0.0621 | 0.0049 | 0.0012 | 0.5393 | 0.9903 | 0.0000 | 0.0000 | 0.0000 |
| euclidean_20 | -0.0649 | 0.0000 | 0.0000 | 0.3245 | 0.9181 | 0.0000 | 0.0000 | 0.0000 |
| jaccard_2 | -0.431 | 0.4610 | 0.0000 | 0.0023 | 0.5842 | 0.0001 | 0.0003 | 0.0005 |
| jaccard_20 | -0.2166 | 0.0000 | 0.0000 | 0.0000 | 0.7662 | 0.0000 | 0.0000 | 0.0000 |
| bray_2 | -1.1021 | 0.0001 | 0.0000 | 0.0000 | 0.7948 | 0.0073 | 0.0132 | 0.0082 |
| bray_15 | -0.2459 | 0.0032 | 0.0000 | 0.0000 | 1.0804 | 0.0000 | 0.0000 | 0.0000 |

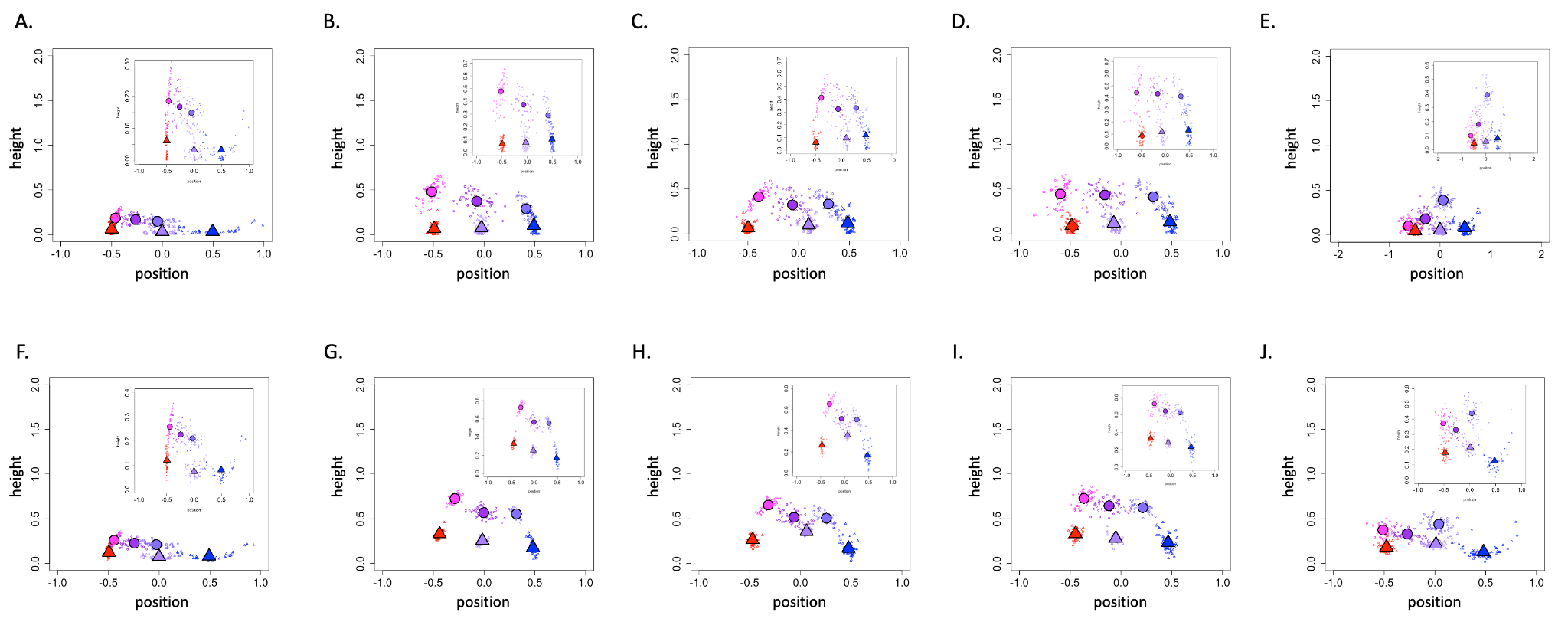

**Figure 5.3.** Position of the hybrid microbiota along the main parental axis of variation and height of the hybrid microbiota perpendicular to the parental axis of variation for skin ASVs (circles) of all hybrids (purple), hybrids at SBluG (magenta), and hybrids at SNW (blue-purple). Null models (triangles) are also shown for random subsamples of the *Aspidoscelis inornatus* population at SBluG (red), the *A. marmoratus* population at SNW (blue), and microbiota derived from ‘breeding’ (see Materials and Methods) *A. inornatus* at SBluG and *A. marmoratus* at SNW (light purple). In all panels, small open circles/triangles represent individual subsamples, while large filled circles/triangles represent means over 50 subsamples. The large diamond in panel (D) represents a median because the mean was strongly influenced by one outlier. Panels (A-E) use the first two PCA/PCoA axes, panels (F-I) use the first 20 PCA/PCoA axes and panel (J) uses the first 15 PCA/PCoA axes. Distance metrics for each panel are: Euclidean (A, F), Jaccard (B, G), Bray-Curtis (C, H), unweighted UniFrac (D, I), and weighted UniFrac (E, J). Insets show close-ups of the regions of interest in each panel.

**Table 5.7.** Position of the overall (i.e., both locations) hybrid microbiota along (median x) and perpendicular to (median y) the main parental axis of variation for skin ASVs. Median hybrid positions are calculated over 15 subsamples of 12 randomly selected *A. neomexicanus* individuals. Column 2-4 show p-values for Mann-Whitney U tests assessing whether the median position of the hybrid along the x-axis is significantly different than null models generated (1) using 12 randomly selected *A. inornatus* (ino) (2) using 12 randomly selected *A. marmoratus* (marm) and (3) summing the microbiota of 12 randomly selected *A. inornatus* and 12 randomly selected *A. marmoratus*. Column 6-8 show p-values for Mann-Whitney U tests assessing whether the median position of the hybrid along the y-axis is significantly greater than the same three null models. Yellow is used for tests significantly different at p>0.05.

|  | mean x | ino | marm | ino + marm | mean y | ino | marm | ino + marm |
| --- | --- | --- | --- | --- | --- | --- | --- | --- |
| euclidean_2 | -0.2890 | 0.0000 | 0.0000 | 0.0000 | 0.1779 | 0.0000 | 0.0000 | 0.0000 |
| euclidean_20 | -0.2680 | 0.0000 | 0.0000 | 0.0000 | 0.2337 | 0.0000 | 0.0000 | 0.0000 |
| jaccard_2 | -0.0801 | 0.0000 | 0.0000 | 0.2328 | 0.3653 | 0.0000 | 0.0000 | 0.0000 |
| jaccard_20 | -0.0091 | 0.0000 | 0.0000 | 0.7437 | 0.5622 | 0.0000 | 0.0000 | 0.0000 |
| bray_2 | -0.0809 | 0.0000 | 0.0000 | 0.0000 | 0.3343 | 0.0000 | 0.0003 | 0.0000 |
| bray_20 | -0.0229 | 0.0000 | 0.0000 | 0.0164 | 0.5005 | 0.0000 | 0.0000 | 0.0000 |
| unifrac_2 | -0.1717 | 0.0000 | 0.0000 | 0.1160 | 0.4294 | 0.0000 | 0.0000 | 0.0000 |
| unifrac_20 | -0.0690 | 0.0000 | 0.0000 | 0.5125 | 0.6373 | 0.0000 | 0.0000 | 0.0000 |
| wunifrac_2 | -0.3144 | 0.0000 | 0.0000 | 0.0000 | 0.2120 | 0.0000 | 0.0043 | 0.0001 |
| wunifrac_20 | -0.2296 | 0.0000 | 0.0000 | 0.0000 | 0.3342 | 0.0000 | 0.0000 | 0.0000 |

**Table 5.8.** Position of hybrid microbiota from SNW along (median x) and perpendicular to (median y) the main parental axis of variation for skin ASVs. Median hybrid positions are calculated over 15 subsamples of 12 randomly selected *A. neomexicanus* individuals. Column 2-4 show p-values for Mann-Whitney U tests assessing whether the median position of the hybrid along the x-axis is significantly different than null models generated (1) using 12 randomly selected *A. inornatus* (ino) (2) using 12 randomly selected *A. marmoratus* (marm) and (3) summing the microbiota of 12 randomly selected *A. inornatus* and 12 randomly selected *A. marmoratus*. Column 6-8 show p-values for Mann-Whitney U tests assessing whether the median position of the hybrid along the y-axis is significantly greater than the same three null models. Yellow is used for tests significantly different at *p*>0.05.

|  | mean x | ino | marm | ino + marm | mean y | ino | marm | ino +  marm |
| --- | --- | --- | --- | --- | --- | --- | --- | --- |
| euclidean_2 | -0.0581 | 0.0000 | 0.0000 | 0.1485 | 0.1509 | 0.0000 | 0.0000 | 0.0000 |
| euclidean_20 | -0.0376 | 0.0000 | 0.0000 | 0.2671 | 0.2112 | 0.0000 | 0.0000 | 0.0000 |
| jaccard_2 | 0.3951 | 0.0000 | 0.0000 | 0.0000 | 0.3282 | 0.0000 | 0.0000 | 0.0000 |
| jaccard_20 | 0.3131 | 0.0000 | 0.0000 | 0.0000 | 0.5646 | 0.0000 | 0.0000 | 0.0000 |
| bray_2 | 0.2703 | 0.0000 | 0.0000 | 0.0000 | 0.3635 | 0.0000 | 0.0000 | 0.0000 |
| bray_20 | 0.2692 | 0.0000 | 0.0000 | 0.0000 | 0.5032 | 0.0000 | 0.0000 | 0.0000 |
| unifrac_2 | 0.2863 | 0.0000 | 0.0000 | 0.0000 | 0.4538 | 0.0000 | 0.0000 | 0.0000 |
| unifrac_20 | 0.2283 | 0.0000 | 0.0000 | 0.0000 | 0.6159 | 0.0000 | 0.0000 | 0.0000 |
| wunifrac_2 | 0.0257 | 0.0000 | 0.0000 | 0.7130 | 0.3830 | 0.0000 | 0.0000 | 0.0000 |
| wunifrac_20 | 0.0406 | 0.0000 | 0.0000 | 0.2671 | 0.4501 | 0.0000 | 0.0000 | 0.0000 |

**Table 5.9.** Position of hybrid microbiota from SBluG along (median x) and perpendicular to (median y) the main parental axis of variation for skin ASVs. Median hybrid positions are calculated over 15 subsamples of 12 randomly selected *A. neomexicanus* individuals. Column 2-4 show p-values for Mann-Whitney U tests assessing whether the median position of the hybrid along the x-axis is significantly different than null models generated (1) using 12 randomly selected *A. inornatus* (ino) (2) using 12 randomly selected *A. marmoratus* (marm) and (3) summing the microbiota of 12 randomly selected *A. inornatus* and 12 randomly selected *A. marmoratus*. Column 6-8 show p-values for Mann-Whitney U tests assessing whether the median position of the hybrid along the y-axis is significantly greater than the same three null models. Yellow is used for tests significantly different at *p*>0.05.

|  | mean x | ino | marm | ino + marm | mean y | ino | marm | ino + mar1 |
| --- | --- | --- | --- | --- | --- | --- | --- | --- |
| euclidean_2 | -0.4637 | 0.0007 | 0.0000 | 0.0000 | 0.1840 | 0.0000 | 0.0000 | 0.0000 |
| euclidean_20 | -0.4439 | 0.0000 | 0.0000 | 0.0000 | 0.2588 | 0.0000 | 0.0000 | 0.0000 |
| jaccard_2 | -0.5341 | 0.1370 | 0.0000 | 0.0000 | 0.4559 | 0.0000 | 0.0000 | 0.0000 |
| jaccard_20 | -0.2933 | 0.0000 | 0.0000 | 0.0000 | 0.7179 | 0.0000 | 0.0000 | 0.0000 |
| bray_2 | -0.4103 | 0.0086 | 0.0000 | 0.0000 | 0.3977 | 0.0000 | 0.0000 | 0.0000 |
| bray_20 | -0.3355 | 0.0000 | 0.0000 | 0.0000 | 0.6413 | 0.0000 | 0.0000 | 0.0000 |
| unifrac_2 | -0.6125 | 0.0000 | 0.0000 | 0.0000 | 0.4078 | 0.0000 | 0.0001 | 0.0000 |
| unifrac_20 | -0.3812 | 0.0043 | 0.0000 | 0.0000 | 0.7242 | 0.0000 | 0.0000 | 0.0000 |
| wunifrac_2 | -0.6302 | 0.0001 | 0.0000 | 0.0000 | 0.1011 | 0.0093 | 0.6281 | 0.1164 |
| wunifrac_20 | -0.5285 | 0.0057 | 0.0000 | 0.0000 | 0.3720 | 0.0000 | 0.0000 | 0.0000 |

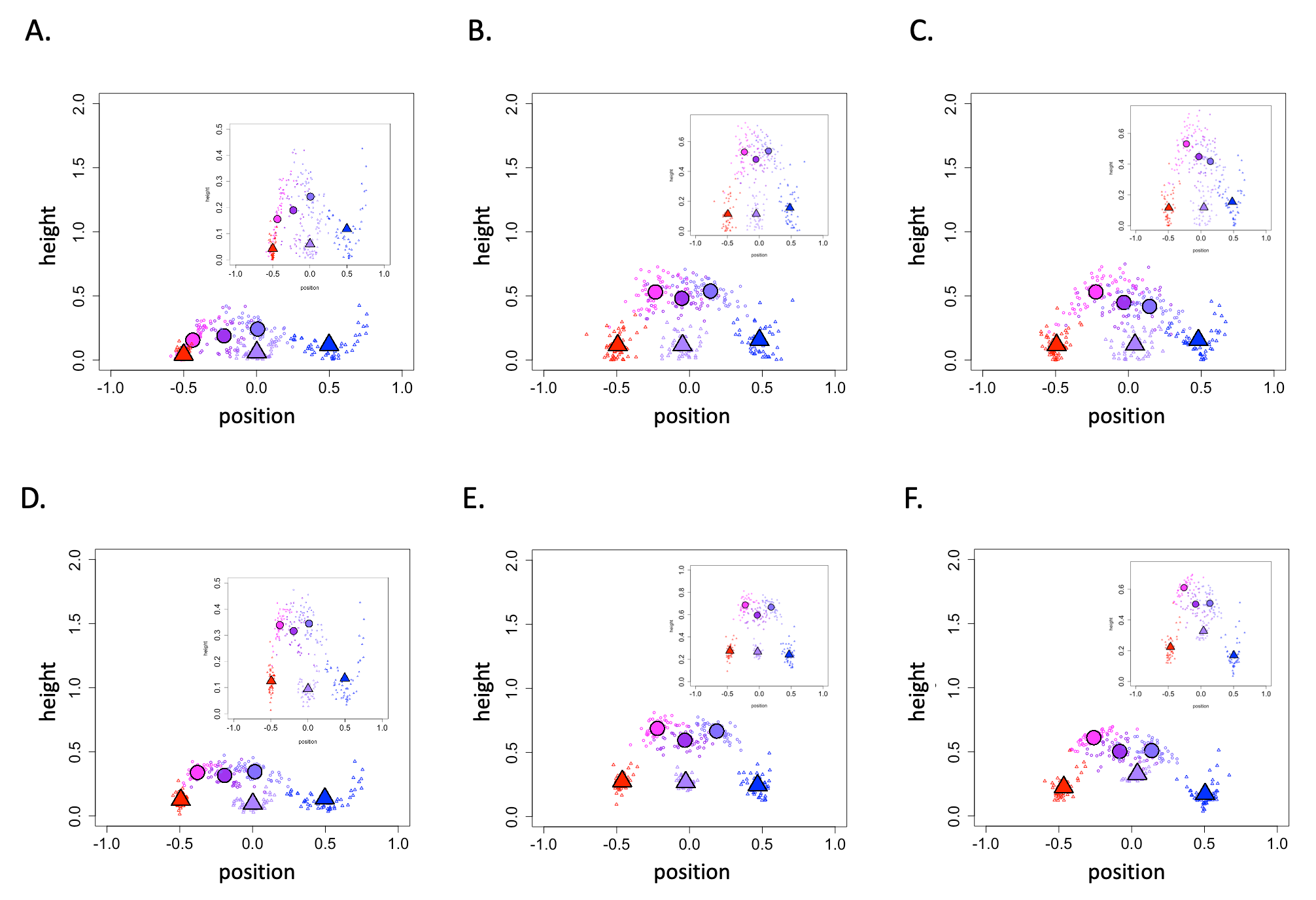

**Figure 5.4.** Position of the hybrid microbiota along the main parental axis of variation and height of the hybrid microbiota perpendicular to the parental axis of variation for skin genera (circles) of all hybrids (purple), hybrids at SBluG (magenta), and hybrids at SNW (blue-purple). Null models (triangles) are also shown for random subsamples of the *Aspidoscelis inornatus* population at SBluG (red), the *A. marmoratus* population at SNW (blue), and microbiota derived from ‘breeding’ (see Materials and Methods) *A. inornatus* at SBluG and *A. marmoratus* at `SNW (light purple). In all panels, small open circles/triangles represent individual subsample samples, while large filled circles/triangles represent means over 50 subsample samples. Panels (A-C) use the first two PCA/PCoA axes, while panels (D-F) use the first 20 PCA/PCoA axes. Distance metrics for each panel are: Euclidean (A, D), Jaccard (B, E), and Bray-Curtis (C, F).

**Table 5.10.** Position of the overall (i.e., both locations) hybrid microbiota along (median x) and perpendicular to (median y) the main parental axis of variation for skin genera. Median hybrid positions are calculated over 15 subsamples of 12 randomly selected *A. neomexicanus* individuals. Column 2-4 show p-values for Mann-Whitney U tests assessing whether the median position of the hybrid along the x-axis is significantly different than null models generated (1) using 12 randomly selected *A. inornatus* (ino) (2) using 12 randomly selected *A. marmoratus* (marm) and (3) summing the microbiota of 12 randomly selected *A. inornatus* and 12 randomly selected *A. marmoratus*. Column 6-8 show p-values for Mann-Whitney U tests assessing whether the median position of the hybrid along the y-axis is significantly greater than the same three null models. Yellow is used for tests significantly different at *p*>0.05.

|  | mean x | ino | marm | ino + marm | mean y | ino | marm | ino + marm |
| --- | --- | --- | --- | --- | --- | --- | --- | --- |
| euclidean_2 | -0.197 | 0.0000 | 0.0000 | 0.0006 | 0.1927 | 0.0001 | 0.0936 | 0.0002 |
| euclidean_20 | -0.1756 | 0.0000 | 0.0000 | 0.0004 | 0.3065 | 0.0000 | 0.0000 | 0.0000 |
| jaccard_2 | -0.0573 | 0.0000 | 0.0000 | 0.5668 | 0.4834 | 0.0000 | 0.0000 | 0.0000 |
| jaccard_20 | -0.0326 | 0.0000 | 0.0000 | 0.4610 | 0.6026 | 0.0000 | 0.0000 | 0.0000 |
| bray_2 | -0.0092 | 0.0000 | 0.0000 | 0.4124 | 0.4272 | 0.0000 | 0.0000 | 0.0000 |
| bray_15 | -0.0552 | 0.0000 | 0.0000 | 0.0367 | 0.5046 | 0.0000 | 0.0000 | 0.0000 |

**Table 5.11.** Position of hybrid microbiota from SNW along (median x) and perpendicular to (median y) the main parental axis of variation for skin genera. Median hybrid positions are calculated over 15 subsamples of 12 randomly selected *A. neomexicanus* individuals. Column 2-4 show p-values for Mann-Whitney U tests assessing whether the median position of the hybrid along the x-axis is significantly different than null models generated (1) using 12 randomly selected *A. inornatus* (ino) (2) using 12 randomly selected *A. marmoratus* (marm) and (3) summing the microbiota of 12 randomly selected *A. inornatus* and 12 randomly selected *A. marmoratus*. Column 6-8 show p-values for Mann-Whitney U tests assessing whether the median position of the hybrid along the y-axis is significantly greater than the same three null models. Yellow is used for tests significantly different at *p*>0.05.

|  | median x | ino | marm | ino + marm | median y | ino | marm | ino + marm |
| --- | --- | --- | --- | --- | --- | --- | --- | --- |
| euclidean_2 | 0.0397 | 0.0000 | 0.0000 | 0.3892 | 0.2294 | 0.0000 | 0.0014 | 0.0000 |
| euclidean_20 | 0.0432 | 0.0000 | 0.0000 | 0.3669 | 0.326 | 0.0000 | 0.0000 | 0.0000 |
| jaccard_2 | 0.1744 | 0.0000 | 0.0000 | 0.0000 | 0.5311 | 0.0000 | 0.0000 | 0.0000 |
| jaccard_20 | 0.2131 | 0.0000 | 0.0000 | 0.0000 | 0.6511 | 0.0000 | 0.0000 | 0.0000 |
| bray_2 | 0.1793 | 0.0000 | 0.0000 | 0.0164 | 0.4054 | 0.0000 | 0.0000 | 0.0000 |
| bray_15 | 0.1512 | 0.0000 | 0.0000 | 0.0000 | 0.5033 | 0.0000 | 0.0000 | 0.0000 |

**Table 5.12.** Position of hybrid microbiota from SBluG along (median x) and perpendicular to (median y) the main parental axis of variation for skin genera. Median hybrid positions are calculated over 15 subsamples of 12 randomly selected *A. neomexicanus* individuals. Column 2-4 show p-values for Mann-Whitney U tests assessing whether the median position of the hybrid along the x-axis is significantly different than null models generated (1) using 12 randomly selected *A. inornatus* (ino) (2) using 12 randomly selected *A. marmoratus* (marm) and (3) summing the microbiota of 12 randomly selected *A. inornatus* and 12 randomly selected *A. marmoratus*. Column 6-8 show p-values for Mann-Whitney U tests assessing whether the median position of the hybrid along the y-axis is significantly greater than the same three null models. Yellow is used for tests significantly different at *p*>0.05.

|  | mean x | ino | marm | ino + marm | mean y | ino | marm | ino + marm |
| --- | --- | --- | --- | --- | --- | --- | --- | --- |
| euclidean_2 | -0.4353 | 0.0408 | 0.0000 | 0.0000 | 0.1552 | 0.0033 | 0.3565 | 0.0043 |
| euclidean_20 | -0.3771 | 0.0000 | 0.0000 | 0.0000 | 0.3293 | 0.0000 | 0.0000 | 0.0000 |
| jaccard_2 | -0.2454 | 0.0000 | 0.0000 | 0.0003 | 0.5164 | 0.0000 | 0.0000 | 0.0000 |
| jaccard_20 | -0.2279 | 0.0000 | 0.0000 | 0.0000 | 0.6859 | 0.0000 | 0.0000 | 0.0000 |
| bray_2 | -0.2332 | 0.0000 | 0.0000 | 0.0001 | 0.5308 | 0.0000 | 0.0000 | 0.0000 |
| bray_15 | -0.2535 | 0.0000 | 0.0000 | 0.0000 | 0.6152 | 0.0000 | 0.0000 | 0.0000 |

**Table 5.13.** Largest loadings on the parental and perpendicular axes using a PCA for gut ASVs. Average reads per individual of each species for each microbial taxon.

| Rank | Taxon | Mean Loading | *A. inornatus* | *A. neomexicanus* | *A. marmoratus* |
| --- | --- | --- | --- | --- | --- |
| Parental Axis (2D) | | | | | |
| 1 | *Dietzia maris* | 0.813381 | 4236 | 3308 | 12709 |
| 2 | *Corynebacterium testudinoris* | 0.5317166 | 6919 | 0 | 4418 |
| 3 | undescribed Propionibacteriaceae | 0.1098811 | 3362 | 0 | 781 |
| 4 | *Brevibacterium linens-salitolerans* | 0.02689861 | 0 | 0 | 2296 |
| 5 | *Acinetobacter baumannii-calcoaceticus* | 0.02401746 | 203 | 50 | 0 |
| Parental Axis (20D) | | | | | |
| 1 | *Dietzia maris* | 0.7345964 | 4236 | 3308 | 12709 |
| 2 | *Corynebacterium testudinoris* | 0.466264 | 6919 | 0 | 4418 |
| 3 | undescribed Propionibacteriaceae | 0.2090886 | 3362 | 0 | 781 |
| 4 | *Brevibacterium linens-salitolerans* | 0.1700369 | 0 | 0 | 2296 |
| 5 | *Fodinibacter luteus* | 0.1011516 | 0 | 968 | 541 |
| Perpendicular Axis (2D) | | | | | |
| 1 | *Dietzia maris* | 0.6720214 | 4236 | 3308 | 12709 |
| 2 | *Corynebacterium testudinoris* | 0.6495901 | 6919 | 0 | 4418 |
| 3 | undescribed Propionibacteriaceae | 0.1103276 | 3362 | 0 | 781 |
| 4 | *Brevibacterium linens-salitolerans* | 0.03624099 | 0 | 0 | 2296 |
| 5 | *Acinetobacter baumannii-calcoaceticus* | 0.02302417 | 203 | 50 | 0 |
| Perpendicular Axis (20D) | | | | | |
| 1 | *Dietzia maris* | 0.632756 | 4236 | 3308 | 12709 |
| 2 | *Corynebacterium testudinoris* | 0.5942032 | 6919 | 0 | 4418 |
| 3 | undescribed Propionibacteriaceae | 0.1876039 | 3362 | 0 | 781 |
| 4 | undescribed Lachnospiraceae | 0.09250316 | 0 | 0 | 1013 |
| 5 | *Fodinibacter luteus* | 0.0709338 | 0 | 968 | 541 |

**Table 5.14** Largest loadings on the parental and perpendicular axes using a PCA for skin ASVs.

Average reads per individual of each species for each microbial taxon.

| Rank | Taxon | Mean Loading | *A. inornatus* | *A. neomexicanus* | *A. marmoratus* |
| --- | --- | --- | --- | --- | --- |
| Parental Axis (2D) | | | | | |
| 1 | *Fodinibacter luteus* | 0.9745524 | 81 | 6100 | 30723 |
| 2 | *Dietzia maris* | 0.1884224 | 6554 | 615 | 947 |
| 3 | *Arthrobacter agilis* | 0.06280382 | 29 | 7 | 2514 |
| 4 | *Exiguobacterium acetylicum* | 0.0253033 | 1634 | 155 | 949 |
| 5 | *Kocuria polaris* | 0.02081149 | 115 | 216 | 971 |
| Parental Axis (20D) | | | | | |
| 1 | *Fodinibacter luteus* | 0.9526559 | 81 | 6100 | 30723 |
| 2 | *Dietzia maris* | 0.185721 | 6554 | 615 | 947 |
| 3 | *Nocardioides aquiterrae-deserti* | 0.1001629 | 3302 | 310 | 807 |
| 4 | *Arthrobacter agilis* | 0.06807111 | 29 | 7 | 2514 |
| 5 | *Nocardioides mesophilus* | 0.05575261 | 927 | 1869 | 26 |
| Perpendicular Axis (2D) | | | | | |
| 1 | *Dietzia maris* | 0.9650886 | 6554 | 615 | 947 |
| 2 | *Fodinibacter luteus* | 0.1879046 | 81 | 6100 | 30723 |
| 3 | *Geodermatophilus africanus-obscurus* | 0.07275101 | 451 | 2123 | 4348 |
| 4 | *Nocardioides aquiterrae-deserti* | 0.05574435 | 3302 | 311 | 807 |
| 5 | *Corynebacterium testudinoris* | 0.04246251 | 586 | 0 | 159 |
| Perpendicular Axis (20D) | | | | | |
| 1 | *Dietzia maris* | 0.6963135 | 6554 | 615 | 947 |
| 2 | *Nocardioides aquiterrae-deserti* | 0.3195602 | 3302 | 311 | 807 |
| 3 | *Fodinibacter luteus* | 0.2198578 | 81 | 6100 | 30723 |
| 4 | *Nocardioides mesophilus* | 0.1443251 | 927 | 1869 | 26 |
| 5 | Unclassified Lachnospiraceae | 0.1255619 | 1204 | 10 | 0 |
